# A simple method to separate birds and insects in single-pol weather radar data

**DOI:** 10.1101/2021.03.08.434434

**Authors:** Raphaël Nussbaumer, Baptiste Schmid, Silke Bauer, Felix Liechti

## Abstract

Recent and archived data from weather radar networks are extensively used for quantification of continent-wide bird migration pattern. While discriminating birds from weather signals is well established, insect contamination is still a problem. We present a simple method combining two doppler radar products within a single Gaussian-mixture model to estimate the proportions of birds and insects within a single measurement, as well as the density and speed of birds and insects. The method can be applied to any existing archives of vertical bird profiles, such as the ENRAM repository (enram.eu) with no need to recalculate the huge amount of original polar volume data, which often are not available.

## 1 Introduction

Weather radar data are increasingly used for quantifying the flow of nocturnal bird migration (Chilson, Stepanian, & Kelly, 2017; Gauthreaux, 1970; Jacobsen & Lakshmanan, 2017). In the past, the reflectivity from birds in the air and mean radial velocity has been used to quantify bird biomass (Gauthreaux & Belser, 1998), before algorithms were developed to extract birds automatically (Dokter et al., 2019, 2011; Lin et al., 2019; Sheldon, Jeffrey, Winner, Bhambhani, & Bernstein, 2019). These algorithms distinguish between bio-scatterers and precipitation but perform less well in distinguishing between the two main bio-scatterer, birds, and insects. To date, these algorithms have mainly been applied to nightly weather radar data during spring and autumn migration, when birds by far outweigh insects (e.g., Dokter et al., 2018; Nilsson et al., 2019; Van Doren & Horton, 2018). Nonetheless, in some nights large insect masses can also migrate and interfere with or even mask the bird echoes in the radar signal (Leskinen et al., 2011; Westbrook & Eyster, 2017). Consequently, classifying and distinguishing insects and birds remains a major challenge (Bauer et al., 2017). With the progressive replacement of single-polarization weather radar by dual-polarization systems, this classification could be improved considerably (e.g., Bachmann & Zrnic, 2005; Leskinen et al., 2011; Stepanian, Horton, Melnikov, Zrnić, & Gauthreaux, 2016). However, today almost all archived weather radar data consist of single-pol weather radar data (ENRAM, 2020; Istok et al., 2009). Additionally, in Europe weather radar data publicly available to ecologists consist only of vertical profiles of bird migration intensities derived from single-pol weather radar data.

To date, insects are usually filtered using a threshold on the standard deviation of radial velocity and/or an additional threshold on absolute airspeed (e.g., Cohen et al., 2020; Dokter et al., 2019; Farnsworth et al., 2016; Horton et al., 2020, 2019; Horton, Shriver, & Buler, 2015; Horton, Van Doren, Stepanian, Farnsworth, & Kelly, 2016; La Sorte et al., 2015; Nilsson et al., 2019; Nussbaumer et al., 2019). Here, we combine these features within a Gaussian-mixture model to estimate the proportions of birds and insects within a single measurement, as well as their density and speed.

## 2 Data

### 2.1 Weather data

We used vertical profiles of reflectivity [cm^2^/km^3^], ground speed [m/s], direction [°] and standard deviation of the radial velocity [m/s], generated with the vol2bird software (Dokter et al., 2011) from single-polarization polar volumes archived in a publicly accessible ENRAM repository (ENRAM, 2020). We used data from 37 weather radars in France, Germany, the Netherlands and Belgium operating between 13 February 2018 and 1 January 2019. Each radar provides a spatial and temporal datapoint, with a vertical resolution of 200m (0-5000m a.s.l.) and a temporal resolution of 5 min. Because we were primarily interested in quantifying nocturnal bird migration, we limit the analysis to night-time as defined by the local sunrise and sunset at each radar location.

### 2.2 Cleaning

We manually cleaned the vertical profile of reflectivity of each radar using a MATLAB dedicated GUI program (detailed in Appendix A of (Nussbaumer et al., 2019). We then only kept the ground speed values where reflectivity values were available, thus removing erroneous speeds. While high-reflectivity weather events (i.e., mainly rain) were easily removed during this procedure, slow moving weather events with low reflectivity (e.g., snow, mist) were difficult to identify visually and therefore were still present at this stage.

### 2.3 Airspeed

Airspeed was computed by taking the ground speed obtained from weather radar and subtracting the wind speed provided by the ERA reanalysis product interpolated at the same location and time (Copernicus Climate Change Service (C3S), 2017).

## 3 Methodology

We estimate the proportion of birds and insects in the sampled volume by comparing the measured airspeed and standard deviation of the radial velocity to the typical signature produced by birds and insects (**Figure 1**). This contrast with previous approaches that applied fixed thresholds in standard deviation of the radial velocity and/or airspeed.

**Figure 1.**
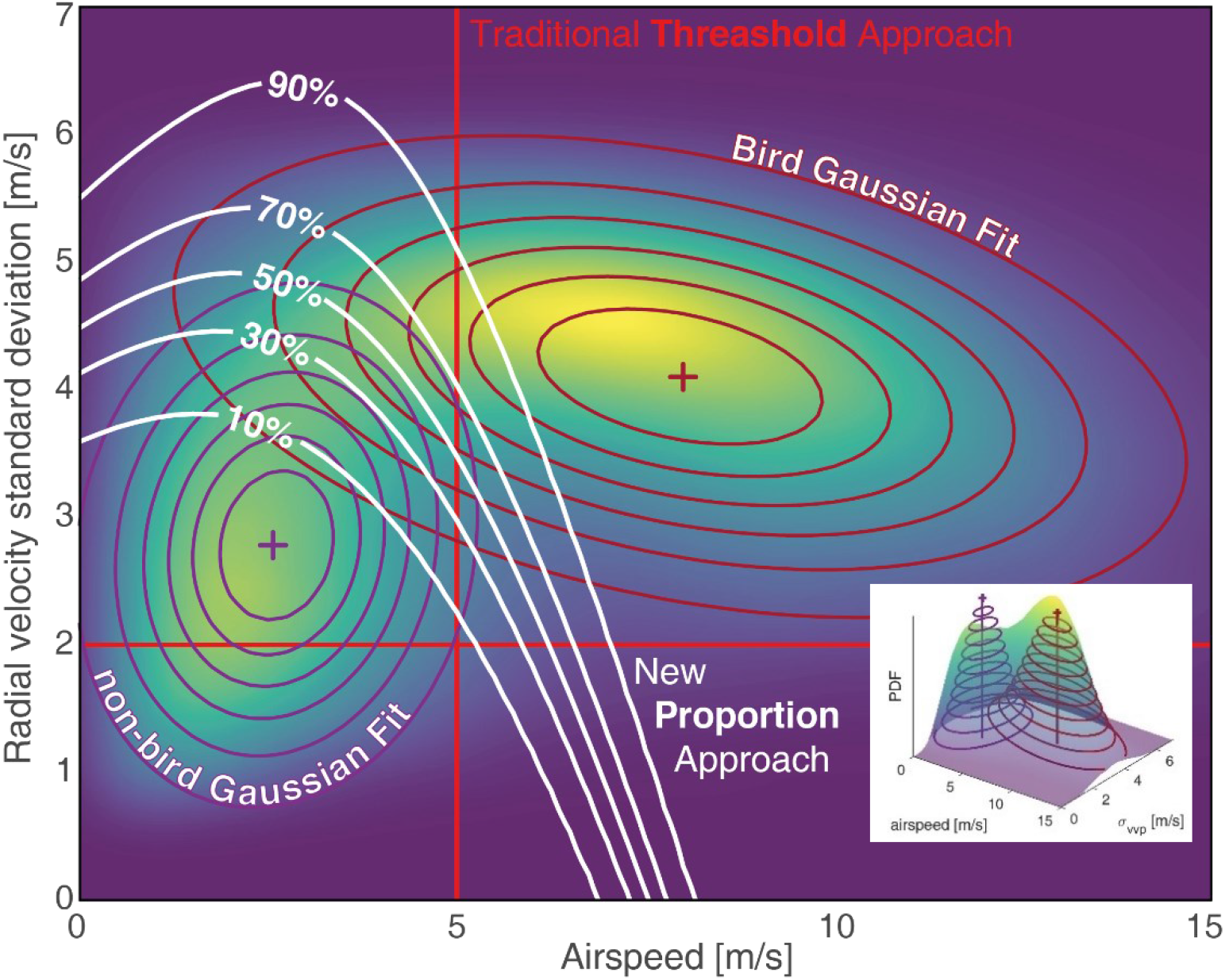
Joint probability distribution of airspeed and radial velocity standard deviation. The empirical probability density function (surface in color scale) shows two peaks: the one on the top-right corresponds to birds and the one on the bottom-left to insects. The two Gaussian distribution fitted to the data are shown in contour lines (red and purple). The previous approaches (red thick straight lines) filtered out all sampled volumes with an airspeed lower than 5m/s and/or radial velocity standard deviation lower than 2 m/s as insect. The new method determines the proportion of bird (white contour line) based on the ratio of the two Gaussian distribution adjusted for the spatio-temporal variation.

We first learn the typical signature of birds and insects by fitting a two-component Gaussian mixture model to the dataset (i.e., one component represents birds and the other insects) (section 3.1). Realizing that relevant insect-like echoes were present during winter, we separate the insect-like echoes into insect and weather-related signals (mainly snowfall) based on the timing of their presence (section 3.2). We then use the fitted model to determine the proportion of bird and insect for all datapoints (section 3.3). Finally, we also correct bird density (section 3.4) and speed of birds (section 3.5) according to the proportion of birds estimated.

### 3.1 Model fitting

The model used to estimate the proportion of birds and insects is fitted in two steps.

In the first step, we learn the distribution of airspeed and radial velocity standard deviation signature of birds and insects (**Figure 1**). Assuming that birds and insects fly in a similar manner everywhere, we use all datapoints (i.e., all radars, all altitudes, all times) to compute the empirical probability density function (PDF) of airspeed and standard deviation of radial velocity from kernel density smoother (ksdensity in MATLAB). We then fitted a two components Gaussian mixture model on that empirical PDF with the Expectation-Maximization (fitgmdist on MATLAB). We extract the location (mean) and shape (covariance) parameters from the two Gaussian components (one for the bird and one for the non-bird).

In this second step, we account for spatial and temporal variation in the presence of birds and insects (**Figure 2** and **Figure 3**). We build the empirical PDF for each month and each radar separately (see Supplementary Material 1) and fitted for each the amplitude of the two Gaussian components, using the location and shape parameters estimated in the first step. Because the empirical probability distribution function is normalized, the sum of the two Gaussian amplitudes is one. Therefore, we can fit it with a single parameter -the amplitude ratio, which ranges from 0 (only insects) to 1 (only birds) (**Figure 3**). Finally, to smooth the transitions from one month to the next, we temporally interpolated the amplitude ratios of each radar with a shape-preserving piecewise cubic interpolation (pchip on MATLAB) (**Figure 4**).

**Figure 2.**
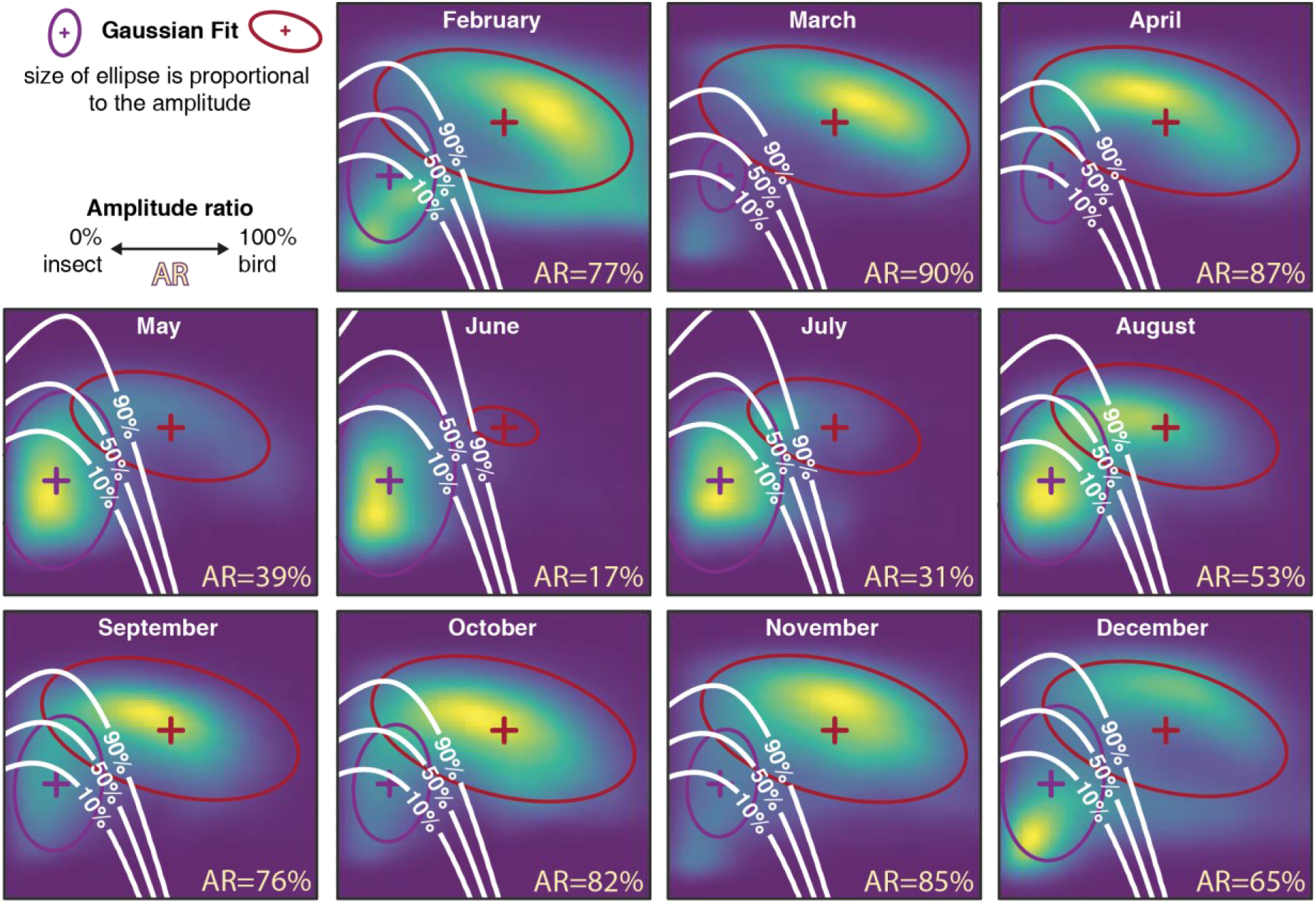
Monthly joint probability density function of airspeed and radial velocity standard deviation for the empirical (color-scaled surface) and fitted Gaussian distributions of birds (red lines) and insects (purple lines). The amplitude ratio quantifies the relative abundance of bird versus insect and is computed as the relative amplitude of the two Gaussians. The white contour line depicts the mapping of the proportion of birds and insect resulting from the fitted model. The x-and y-axis unit and range are the same than in **Figure 1**, but the color-scale is normalized each month.

**Figure 3.**
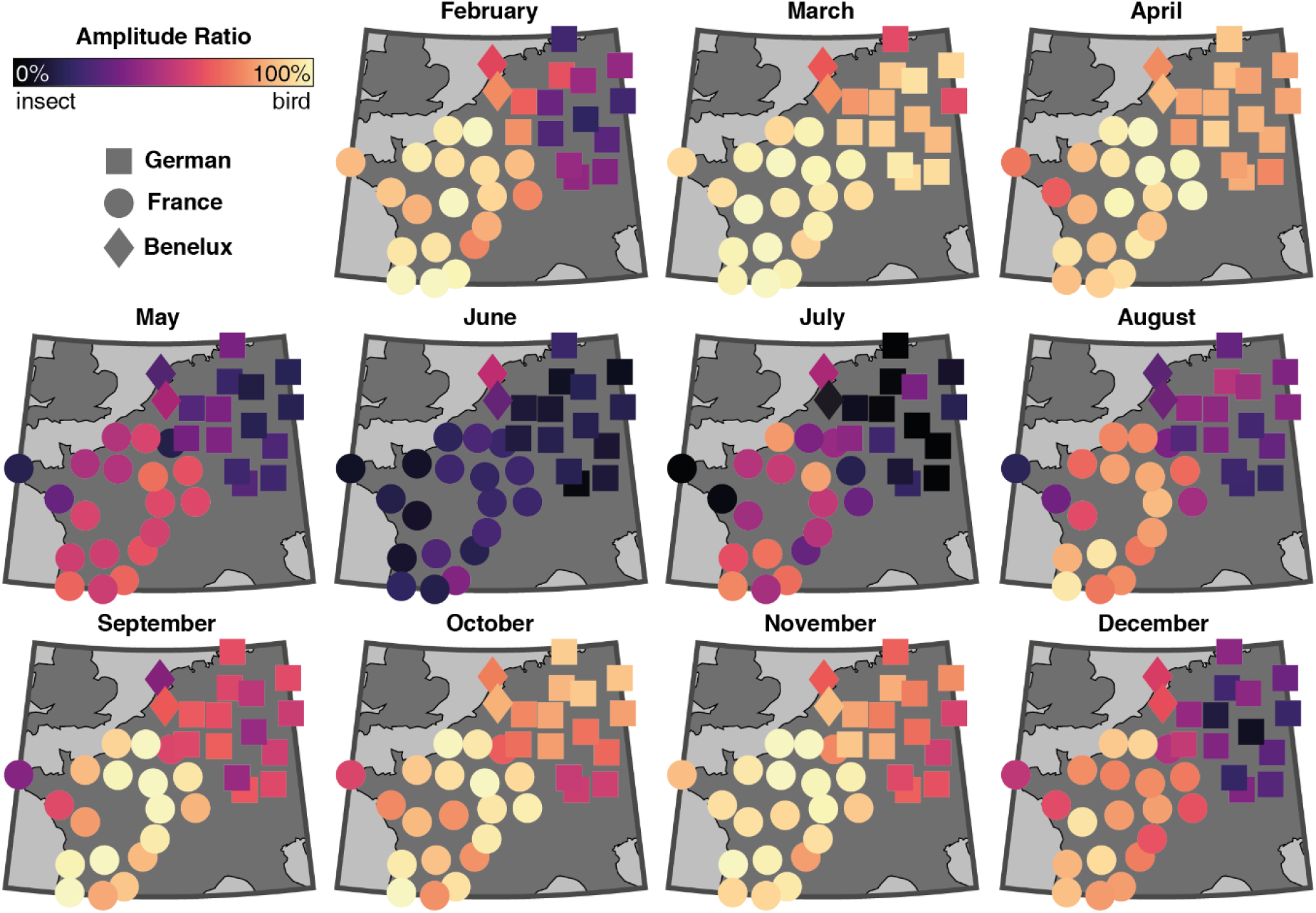
Amplitude ratio fitted for each radar and each month. In addition to the main seasonal trend (already visible in **Figure 2**), a smaller effect appears mainly during winter in the northern radars (mainly, but no only German), where slow-moving weather events (e.g., snow or mist) are picked up as insects by the radars.

**Figure 4.**
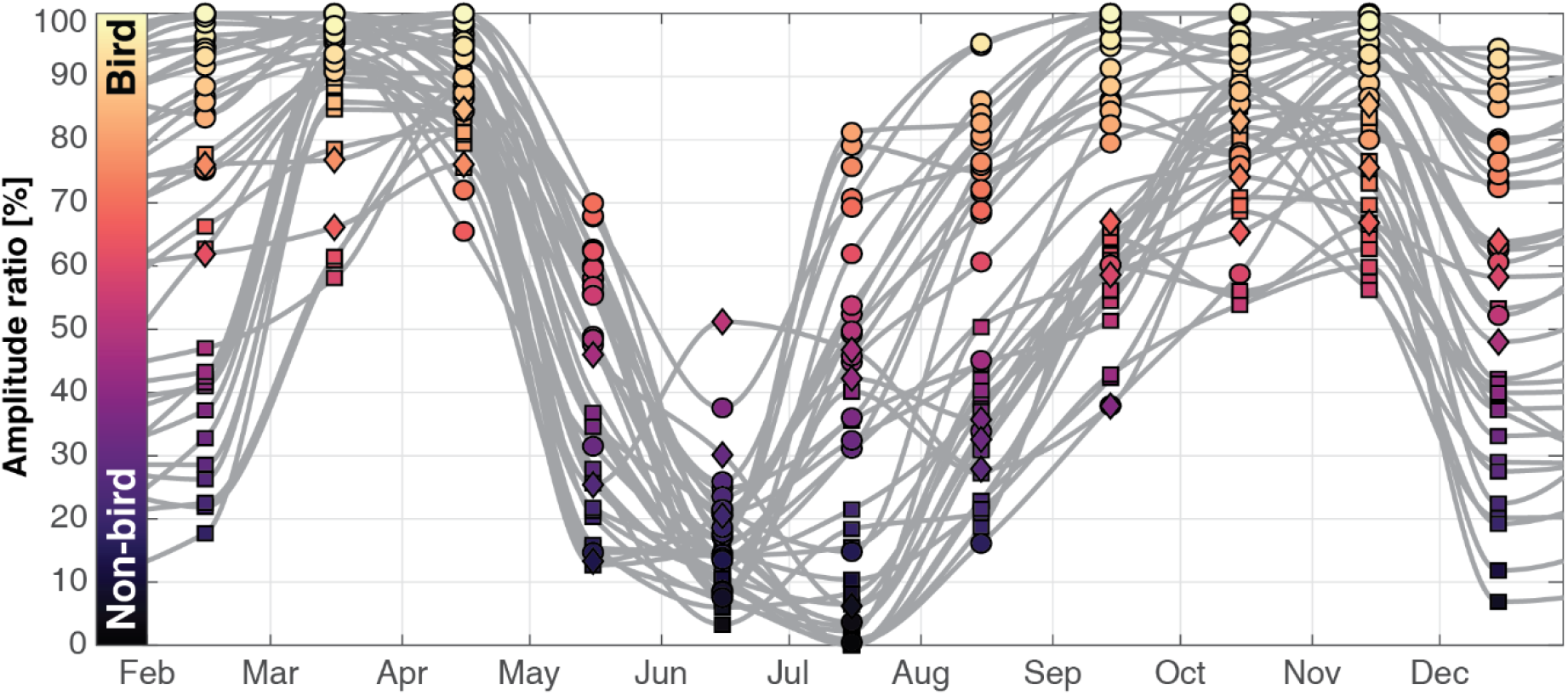
Seasonal variation in amplitude ratio for each radar. The monthly amplitude ratios of each radar (circle for France radar, square for German and diamond for Benelux) were temporally interpolated (grey lines).

### 3.2 Separation of insect and weather

While rain is easily recognized for its high reflectivity signal and therefore correctly removed in the initial cleaning step, slow moving weather events with low reflectivity such as snow and mist (low airspeed and low radial velocity standard deviation) are harder to remove from the dataset (see winter insect-like signal from Germany in **Figure 2** and **Figure 3**). Yet, because snow and mist generally occur in winter and insects mainly occur in summer, we can separate them based on their temporal occurrence.

For each radar, we fitted the complement of the amplitude ratio (i.e., 1-amplitude ratio) with the sum of two skewed normal density functions over time, one matching insects (peaking in summer) and the other weather (peaking in winter, **Figure 5a**). The insect ratio is finally determined as the ratio of the insect normal density function over the sum of the normal density functions (**Figure 5b**).

**Figure 5.**
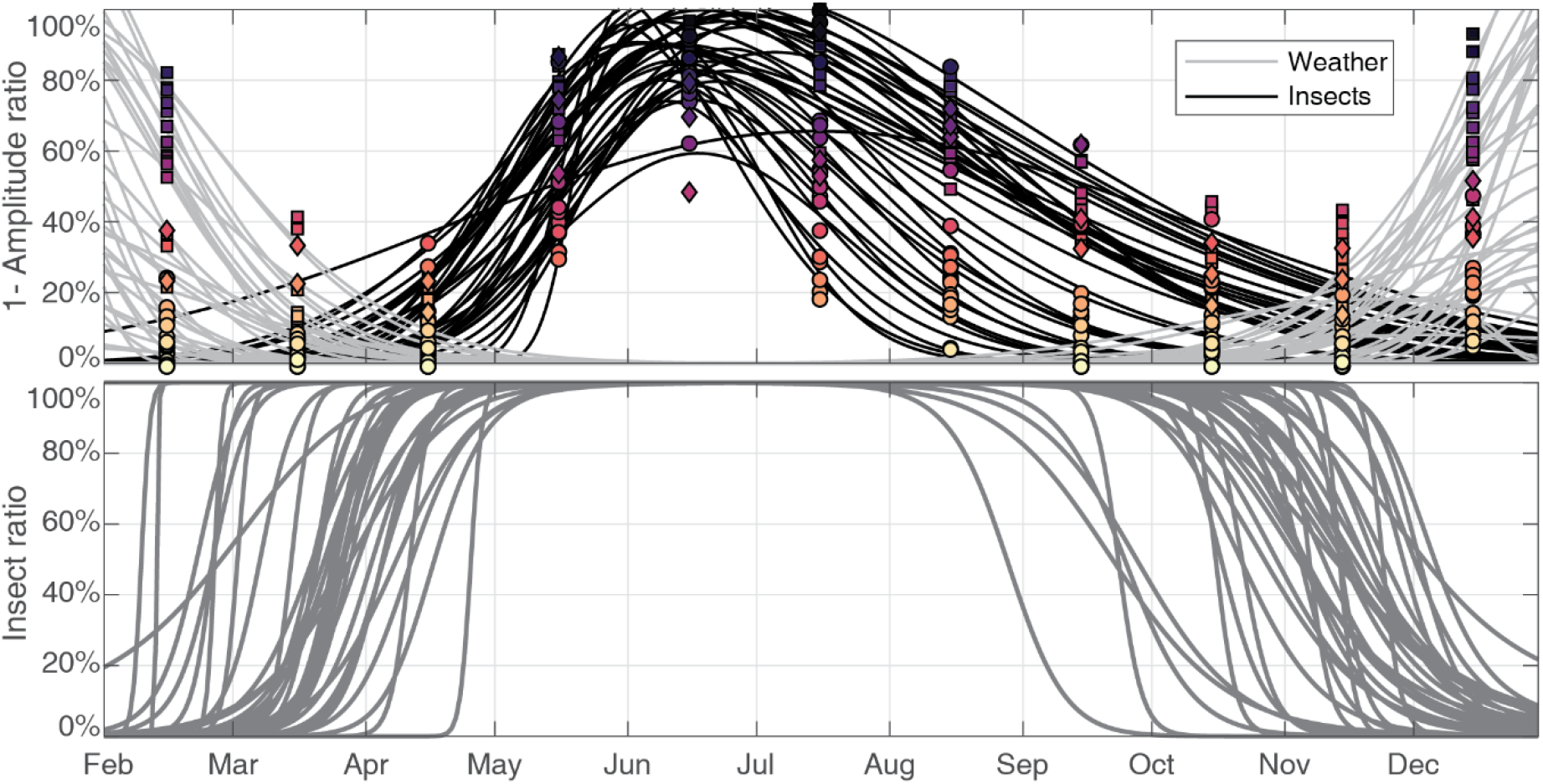
The insect-like signal (1-amplitude ratio) is separated into insect and weather. (a) The sum of two skew normal density functions representing insects (black) and weather (grey) is fitted for each radar (circle for France radar, square for German and diamond for Benelux). (b) The proportion of insects in the original insect signal is estimated from the fitted density distribution functions of **Figure 5a**.

### 3.3 Estimating the proportion of birds and insects

The procedure to estimate the proportion of birds, insects, and remaining weather signal for each datapoint is as follows. For each datapoint, we compute the amplitude ratio from the interpolation in **Figure 4** based on radar and time of year. We then built the two Gaussian distributions using both the amplitude ratio and the means and covariances estimated in **Figure 1**. Next, we determine the probabilities of the bird and non-bird Gaussian probability density functions using the known value of airspeed and radial velocity standard deviation at the datapoint. Finally, the proportion of birds for that particular datapoint is estimated by normalizing the probability of birds with the sums of probabilities of birds and non-birds.

The proportion of insects is determined by multiplying the proportion of non-birds (i.e., 1-proportion of birds) with the insect ratio determined in **Figure 5**b.

### 3.4 Bird density correction

After quantifying the proportion of birds, insects and weather, the bird and insect reflectivity are estimated by dividing the original values of bird reflectivity according to their respective proportion. Bird reflectivity is then converted to bird density assuming a radar cross section of 11 cm^2^ (Dokter et al., 2011).

### 3.5 Ground and air speed correction

We propose here a method to correct the ground speed calculated by vol2bird from the contamination from insect movements according to the proportion of bird estimated in section 3.3.

We formalized this problem with Bayes’ Theorem, where the prior distributions of bird and insect airspeeds is taken from the Gaussian fits of **Figure 1** as normal distribution,

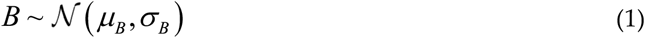

and

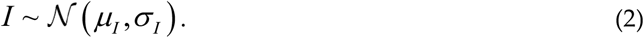

We consider that the airspeed obtained from weather radar is a mixture of birds airspeed and insects airspeed weighted according to the proportion of birds *α* and insects 1 − *α*, as determined in section 3.3. Its distribution is therefore also normal with parameters which can be determined as

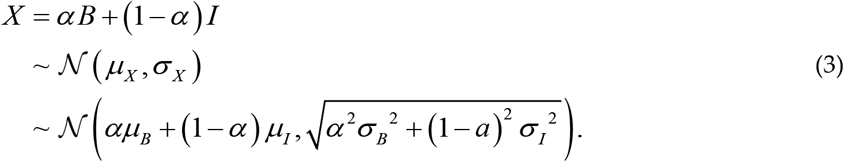

In the Bayes framework, the corrected airspeed for birds is the probability of bird airspeed conditional to the measured airspeed as shown as

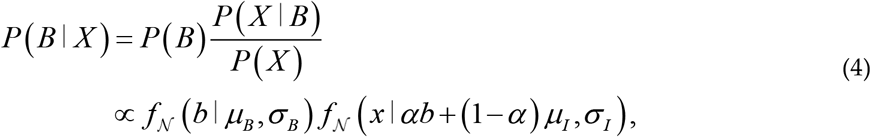

where *f*_𝒩_ is the normal probability distribution function, *x* is the known measured airspeed and *b* is the unknown bird airspeed.

The correction of each airspeed datapoint is performed as follows. For each datapoint, we build the probability distribution of the birds’ airspeed *b* of Equation (4) based on the known proportion of birds *α*, and the original airspeed *x*. The corrected airspeed of birds is then sampled randomly according to the probability density function. Finally, the bird ground speed is determined by re-adding the wind speed to the corrected air speed. The insect ground speed is computed using the sampled bird airspeed and Equation (3).

Figure 6. compares the original airspeed data with the corrected airspeeds for insects and birds. The histograms of the corrected airspeed perfectly reproduce the fitted distribution of insect and bird airspeed by construction. Based on the airspeed distribution, a threshold of 4.8m/s would lead to the best classification of insects and birds. The traditional 5m/s threshold would misclassify 19% of bird samples and 4% of insect samples.

**Figure 6.**
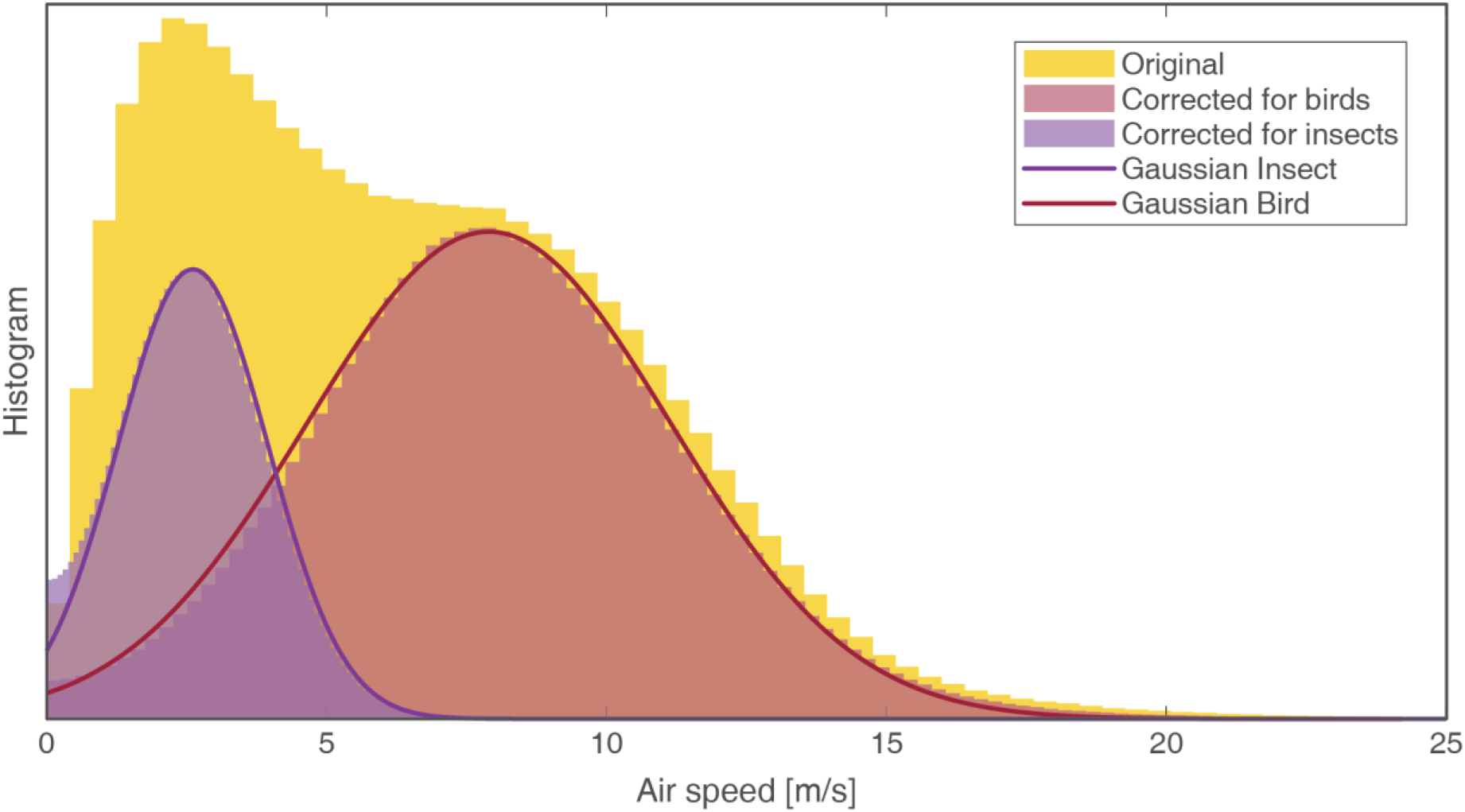
Histogram of airspeed for all datapoints before correction (yellow), corrected for insect (purple) and corrected for bird (red). The Gaussian fit of bird and non-bird from **Figure 1** are shown as line. The histogram before correction shows two modes corresponding to insects and birds. After correction, the airspeed of birds and insects reproduces the Gaussian fit.

## 4 Result

The resulting corrected dataset is available on https://doi.org/10.5281/zenodo.3610184 (Nussbaumer, 2020). **Figure 7** illustrates the weekly mean reflectivity and reflectivity traffic rate (RTR) averaged over all radars for birds, insects, and weather-related signals. Weather related signals account for a small yet significant proportion of the reflectivity in the winter months (Dec-Jan). In contrast, insects contribute significantly to the reflectivity from May to mid-September, with, e.g., more than 90% of the reflectivity attributed to insects in June alone.

**Figure 7.**
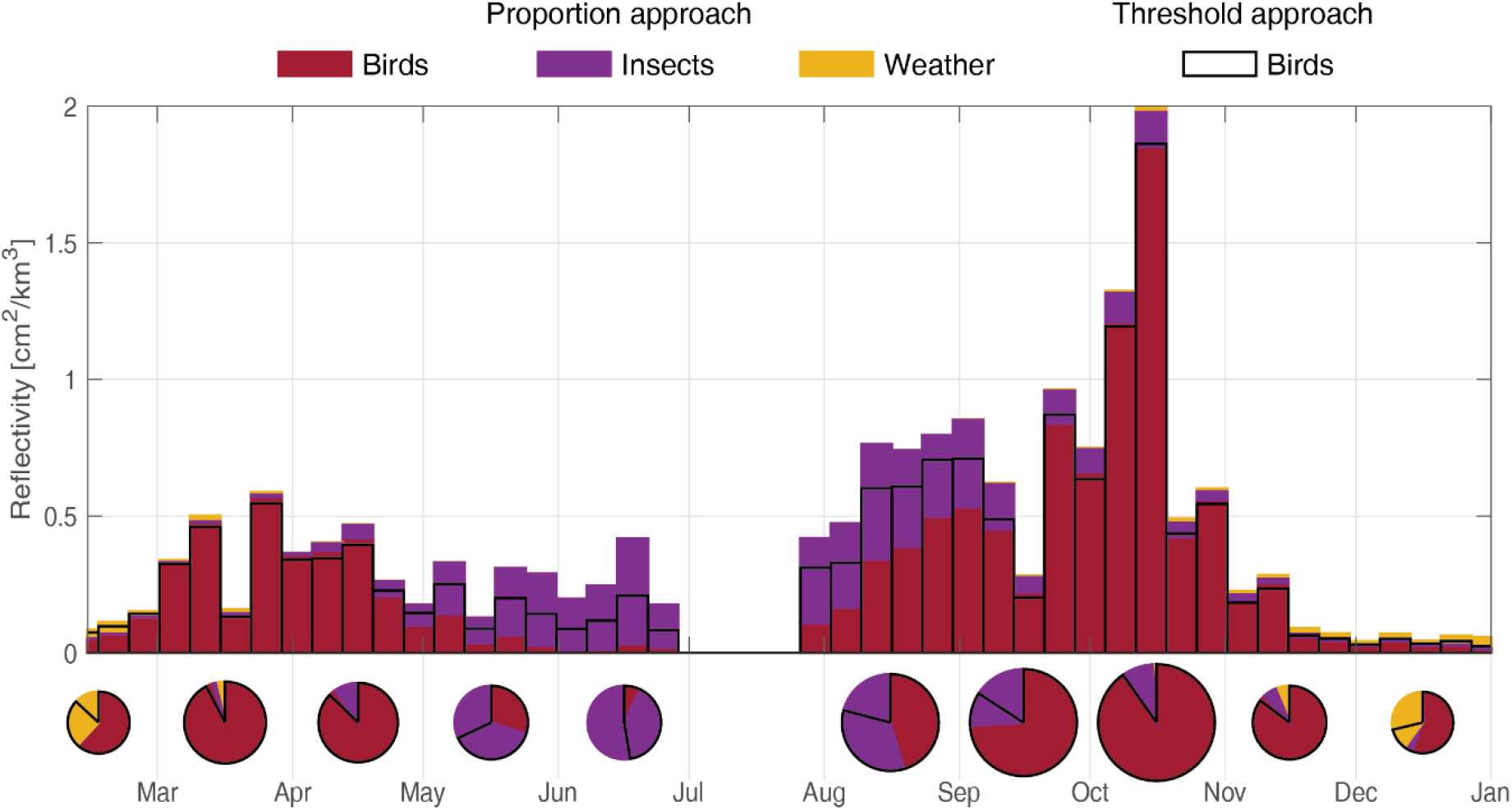
Weekly mean of reflectivity attributed to birds, insects, and weather signal averaged over the 37 radars. The traditional threshold method (‘bird threshold’) is shown as a black line for comparison. The pie charts illustrate the monthly distribution of reflectivity.

In comparison, the traditional threshold approach eliminates datapoints with an airspeed smaller than 5m/s and a radial velocity standard deviation lower than 2m/s. The bird reflectivity resulting from this threshold approach (black line in **Figure 7**) is similar to the proportion approach during the bird migration period (March-May and mid-Sept-mid-Oct), showing that the known threshold from the literature are well calibrated to the airspeed data. Yet, during summer (mid-May to September), the reflectivity derived from the threshold approach is higher than the one derived from the proportion approach (**Figure 1**), as datapoints with airspeed just below 5m/s but with high radial velocity standard deviation (∼5m/s) are discarded by the threshold method while the new approach considers it likely that they are birds.

Reflectivity Traffic Rate (RTR) [reflectivity x speed] (see Supplementary Material 2) assesses the biomass movement (i.e., dynamic instead of static). Because of their lower flight speeds, insects have a smaller proportion in RTR than in reflectivity. However, due to the speed correction, the RTR of birds increases slightly compared to the threshold approach when insects are present (e.g., Aug-sept).

## 5 Discussion

In this study, we presented a new and simple method to separate birds and insects in weather radar signals. The method can easily be applied to existing single-polarization weather radar data archive (e.g., ENRAM, 2020) without the need to recalculate the huge amount of original polar volume data. The separation is based on the differences between birds and insects in airspeed and radial velocity standard deviation (Figure 1). In contrast to simple threshold methods, our approach considers that the velocities derived from the radial doppler-speeds are an average resulting from the movements of all objects (incl. birds and insects) within scanning range of the radar. Consequently, we suggest estimating the proportion of birds rather than whether insects are present or not.

By combining both the average airspeed and the variation of airspeed within the sampled volume we can differentiate between birds and insects. Indeed, birds flying with a low mean airspeed (e.g., 3-5 m/s) typically have a high standard deviation in radial velocity (see skewed orientation of the bird Gaussian contours in **Figure 1**). This is mainly caused by birds flying in different directions, which results in speeds with low average but high variability. Thus, only if we jointly consider both variables can we achieve a better separation. Furthermore, a major strength of the method is its capacity to account for the spatio-temporal variation of birds, insects, and low-reflectivity weather events. This is particularly useful when accounting for the presence for snow which mainly occur in the North (e.g., Germany) during winter (**Figure 3**). Finally, the method can also correctly estimate the airspeed of birds by accounting for the contribution of insects to the average airspeed.

There are, however, simplifying assumptions which were made. Firstly, we calculated the location (mean) and shape (covariance) of the empirical probability density function for the entire dataset, assuming that birds and insect are flying with the same distribution of airspeed and standard deviation of radial velocity throughout the year. However, birds typically fly slower and with a more uniform radial velocity (i.e. lower standard deviation) in warmer months (e.g., April, September) than in colder months (i.e. early spring and late autumn) (**Figure 2**). Indeed, these observations are consistent with the fact that smaller birds (e.g., warblers) with slower optimal airspeeds dominate the long-distance migrations while larger birds (e.g., thrushes) dominate the short-distance migrations (Bruderer & Boldt, 2001). Variable bird airspeed could be accounted in the future with a Gaussian mean function over time. Secondly, airspeed and radial velocity standard deviation are assumed to follow a Gaussian distribution. Despite the imperfect match (**Figure 1**), Gaussian probability density functions were deemed sufficient for the purpose of the separation here, although other distributions could be used in the future to improve the distribution fit (with caution, given the increased complexity and likely minimal benefit).

Lastly, we did not consider potential changes in the airspeed distribution (and amplitude) with altitude as the quality of weather radar at low heights is still questionable (e.g., Nilsson et al., 2018). Nevertheless, future improvement could incorporate the information of altitude for the separation (see Supplementary Material 3).

## Supplementary Materials

SM1: Figures for each radar of the monthly empirical and fitted joint probability distribution of airspeed and radial velocity standard deviation. **SM2**: Altitudinal empirical and fitted joint probability distribution of airspeed and radial velocity standard deviation. **SM3**: Weekly mean of reflectivity traffic rate attributed to birds, insects, and weather signal averaged over the 37 radars.

## Data Availability Statements

The raw data are download from ENRAM repository https://enram.github.io/data-repository/ (ENRAM, 2020). The MATLAB code used for this study can be find on Github (https://rafnuss-postdoc.github.io/BMM/2018/LiveScript/Insect_removal.html). The processed data resulting from the methodology is available at https://doi.org/10.5281/zenodo.3610184 (Nussbaumer, 2020).

## Supporting information

Monthly PDF per radar

Altitudinal PDF

Weekly Reflectivity Traffic Rate

MATLAB code

## Author Contributions

Conceptualization, R.N., F.L. and B.S.; methodology, R.N.; writing—original draft preparation, R.N. and F.L.; writing—review and editing, B.S. and S.B.; supervision, F.L.

## Funding

We acknowledge the European Operational Program for Exchange of Weather Radar Information (EUMETNET/OPERA) for providing access to European radar data, facilitated through a research-only license agreement between EUMETNET/OPERA members and ENRAM (European Network for Radar surveillance of Animal Movements).

We acknowledge the financial support from the Globam project funded by BioDIVERSA, including the Swiss National Science Foundation (31BD30\_184120), Netherlands Organisation for Scientific Research (NWO E10008), Academy of Finland (aka 326315), Belgian Federal Science Policy Office (BelSPO BR/185/A1/GloBAM-BE) and National Science Foundation (NSF 1927743).

## Acknowledgments

We thanks Mathieu Gravey, Lionel Benoit and Grégoire Mariéthoz for useful discussions on the methodology and help with the numerical implementation.

## Conflicts of Interest

The authors declare no conflict of interest.

## Notes

https://doi.org/10.5281/zenodo.3610184

https://enram.github.io/data-repository/

### Competing Interest Statement

The authors have declared no competing interest.

## References

Bachmann, S., & Zrnic, D. (2005). Spectral polarimetry for identifying and separating mixed biological scatterers. 11th Conference on Mesoscale Processes and the 32nd Conference on Radar Meteorology, (V), 295–300.

Bauer, S., Chapman, J. W., Reynolds, D. R., Alves, J. A., Dokter, A. M., Menz, M. M. H., … Shamoun-Baranes, J. (2017). From Agricultural Benefits to Aviation Safety: Realizing the Potential of Continent-Wide Radar Networks. BioScience, 67(10), 912–918. https://doi.org/10.1093/biosci/bix074

Bruderer, B., & Boldt, A. (2001). Flight characteristics of birds: I. Radar measurements of speeds. Ibis, 143(2), 178–204. https://doi.org/10.1111/j.1474-919x.2001.tb04475.x

Chilson, P. B., Stepanian, P. M., & Kelly, J. F. (2017). Radar Aeroecology. In Aeroecology (pp. 277–309). https://doi.org/10.1007/978-3-319-68576-2_12

Cohen, E. B., Horton, K. G., Marra, P. P., Clipp, H. L., Farnsworth, A., Smolinsky, J. A., … Buler, J. J. (2020). A place to land: spatiotemporal drivers of stopover habitat use by migrating birds. Ecology Letters, ele.13618. https://doi.org/10.1111/ele.13618

Copernicus Climate Change Service (C3S). (017). ERA5: Fifth generation of ECMWF atmospheric reanalyses of the global climate. Retrieved February 1, 2019, from Copernicus Climate Change Service Climate Data Store (CDS) website: https://cds.climate.copernicus.eu/cdsapp#/home

Dokter, A. M., Desmet, P., Spaaks, J. H., van Hoey, S., Veen, L., Verlinden, L., … Shamoun-Baranes, J. (2019). bioRad: biological analysis and visualization of weather radar data. Ecography, 42(5), 852–860. https://doi.org/10.1111/ecog.04028

Dokter, A. M., Farnsworth, A., Fink, D., Ruiz-Gutierrez, V., Hochachka, W. M., La Sorte, F. A., … Kelling, S. (2018). Seasonal abundance and survival of North America’s migratory avifauna determined by weather radar. Nature Ecology & Evolution, 2(10), 1603–1609. https://doi.org/10.1038/s41559-018-0666-4

Dokter, A. M., Liechti, F., Stark, H., Delobbe, L., Tabary, P., & Holleman, I. (2011). Bird migration flight altitudes studied by a network of operational weather radars. Journal of The Royal Society Interface, 8(54), 30–43. https://doi.org/10.1098/rsif.2010.0116

ENRAM. (2020). ENRAM data repository for vertical profiles of Birds. Retrieved April 23, 2019, from European Network for the Radar surveillance of Animal Movement website: https://enram.github.io/data-repository/

Farnsworth, A., Van Doren, B. M., Hochachka, W. M., Sheldon, D., Winner, K., Irvine, J., … Kelling, A. S. (2016). A characterization of autumn nocturnal migration detected by weather surveillance radars in the northeastern USA. Ecological Applications, 26(3), 752–770. https://doi.org/10.1890/15-0023/suppinfo

Gauthreaux, S. A. (1970). Weather Radar Quantification of Bird Migration. BioScience, 20(1), 17–20. https://doi.org/10.2307/1294752

Gauthreaux, S. A., & Belser, C. G. (1998). Displays of Bird Movements on the WSR-88D: Patterns and Quantification*. Weather and Forecasting, 13(2), 453–464. https://doi.org/10.1175/1520-0434(1998)013<0453:DOBMOT>2.0.CO;2

Horton, K. G., La Sorte, F. A., Sheldon, D., Lin, T., Winner, K., Bernstein, G., … Farnsworth, A. (2020). Phenology of nocturnal avian migration has shifted at the continental scale. Nature Climate Change, 10(1), 63–68. https://doi.org/10.1038/s41558-019-0648-9

Horton, K. G., Nilsson, C., Van Doren, B. M., La Sorte, F. A., Dokter, A. M., & Farnsworth, A. (2019). Bright lights in the big cities: migratory birds’ exposure to artificial light. Frontiers in Ecology and the Environment, 17(4), 209–214. https://doi.org/10.1002/fee.2029

Horton, K. G., Shriver, W. G., & Buler, J. J. (2015). A comparison of traffic estimates of nocturnal flying animals using radar, thermal imaging, and acoustic recording. Ecological Applications, 25(2), 390–401. https://doi.org/10.1890/14-0279.1.sm

Horton, K. G., Van Doren, B. M., Stepanian, P. M., Farnsworth, A., & Kelly, J. F. (2016). Seasonal differences in landbird migration strategies. The Auk, 133(4), 761–769. https://doi.org/10.1642/AUK-16-105.1

Istok, M. J., Fresch, M., Jing, Z., Smith, S., Murnan, R., Ryzhkov, A., … others. (2009). WSR-88D dual polarization initial operational capabilities. 25th Conf. on Int. Interactive Information and Processing Systems (IIPS) for Meteorology, Oceanography, and Hydrology.

Jacobsen, E., & Lakshmanan, V. (2017). Inferring the State of the Aerosphere from Weather Radar. In Aeroecology (pp. 311–343). https://doi.org/10.1007/978-3-319-68576-2_13

La Sorte, F. A., Hochachka, W. M., Farnsworth, A., Sheldon, D., Van Doren, B. M., Fink, D., & Kelling, S. (2015). Seasonal changes in the altitudinal distribution of nocturnally migrating birds during autumn migration. Royal Society Open Science, 2(12), 150347. https://doi.org/10.1098/rsos.150347

Leskinen, M., Markkula, I., Koistinen, J., Pylkkö, P., Ooperi, S., Siljamo, P., … Tiilikkala, K. (2011). Pest insect immigration warning by an atmospheric dispersion model, weather radars and traps. Journal of Applied Entomology, 135(1–2), 55–67. https://doi.org/10.1111/j.1439-0418.2009.01480.x

Lin, T., Winner, K., Bernstein, G., Mittal, A., Dokter, A. M., Horton, K. G., … Sheldon, D. (2019). M <scp>ist</scp> N <scp>et</scp>?: Measuring historical bird migration in the US using archived weather radar data and convolutional neural networks. Methods in Ecology and Evolution, 10(11), 1908–1922. https://doi.org/10.1111/2041-210X.13280

Nilsson, C., Dokter, A. M., Schmid, B., Scacco, M., Verlinden, L., Bäckman, J., … Liechti, F. (2018). Field validation of radar systems for monitoring bird migration. Journal of Applied Ecology, (October 2017), 1–13. https://doi.org/10.1111/1365-2664.13174

Nilsson, C., Dokter, A. M., Verlinden, L., Shamoun-Baranes, J., Schmid, B., Desmet, P., … Liechti, F. (2019). Revealing patterns of nocturnal migration using the European weather radar network. Ecography, 42(5), 876–886. https://doi.org/10.1111/ecog.04003

Nussbaumer, R. (2020). Vertical profiles time series of bird density and flight speed vector (01.01.2018-01.01.2019). https://doi.org/10.5281/zenodo.3610184

Nussbaumer, R., Benoit, L., Mariethoz, G., Liechti, F., Bauer, S., & Schmid, B. (2019). A Geostatistical Approach to Estimate High Resolution Nocturnal Bird Migration Densities from a Weather Radar Network. Remote Sensing, 11(19), 2233. https://doi.org/10.3390/rs11192233

Sheldon, D., Jeffrey, Winner K., Bhambhani, P., & Bernstein, G. (2019). darkecology/wsrlib: Version 0.2.0. https://doi.org/10.5281/ZENODO.3352264

Stepanian, P. M., Horton, K. G., Melnikov, V. M., Zrnić, D. S., & Gauthreaux, S. A. (2016). Dual-polarization radar products for biological applications. Ecosphere, 7(11). https://doi.org/10.1002/ecs2.1539

Van Doren, B. M., & Horton, K. G. (2018). A continental system for forecasting bird migration. Science, 361(6407), 1115–1118. https://doi.org/10.1126/science.aat7526

Westbrook, J. K., & Eyster, R. S. (2017). Doppler weather radar detects emigratory flights of noctuids during a major pest outbreak. Remote Sensing Applications: Society and Environment, 8, 64–70. https://doi.org/10.1016/j.rsase.2017.07.009

