## Supplementary figures and images for "A simple method to separate birds and insects in single-pol weather radar data"

### Altitudinal PDF

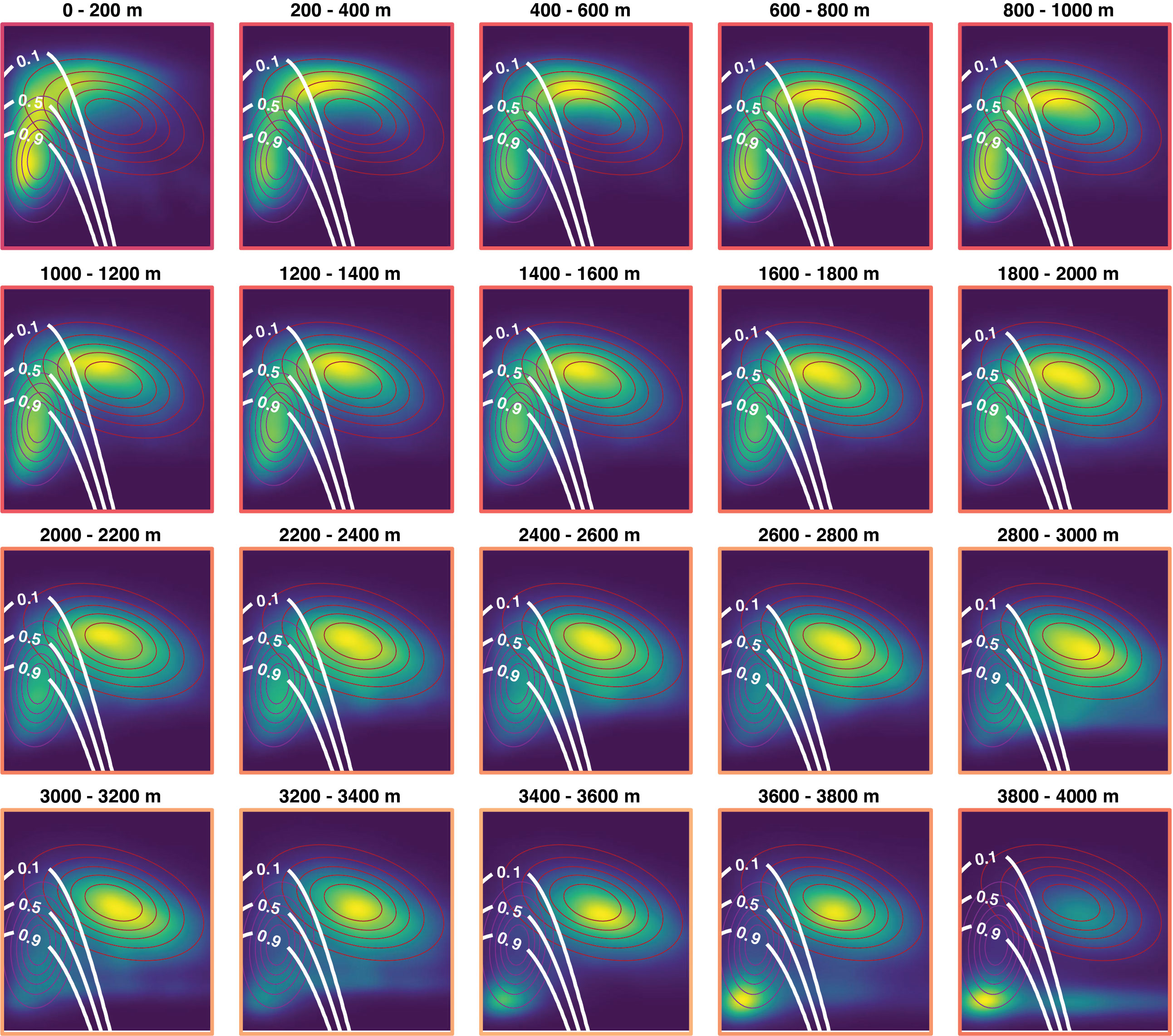

### bewid.png

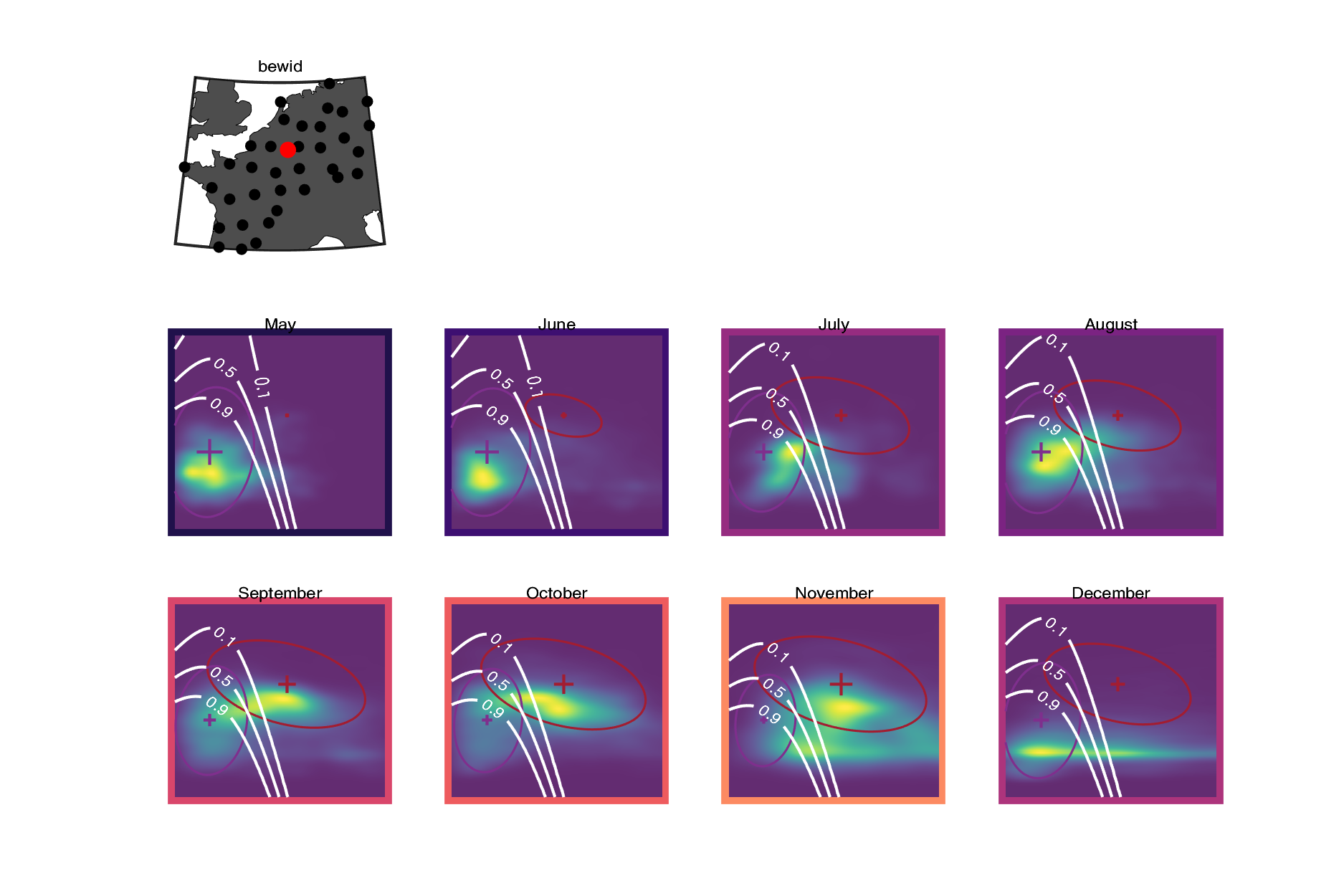

### deboo.png

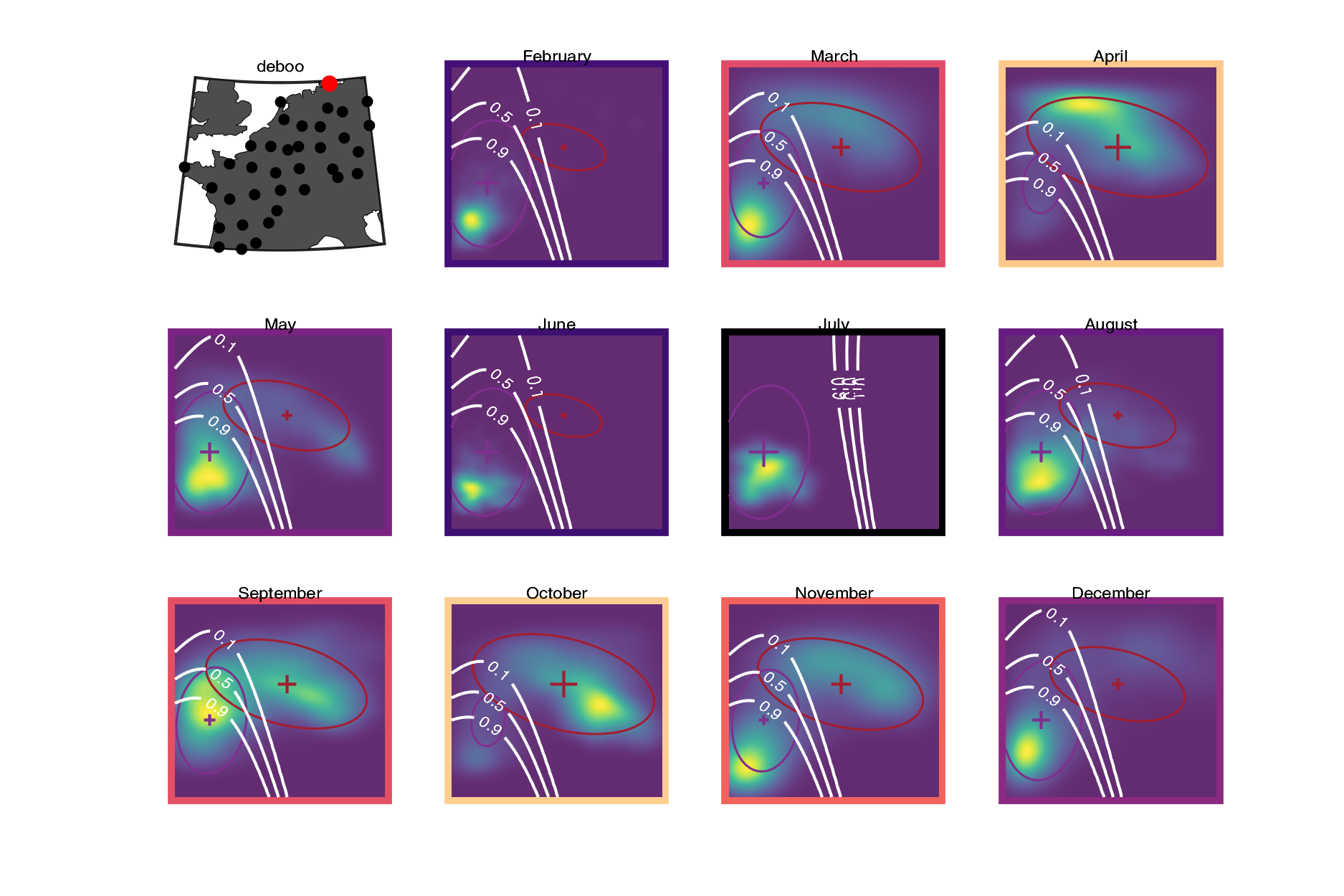

### dedrs.png

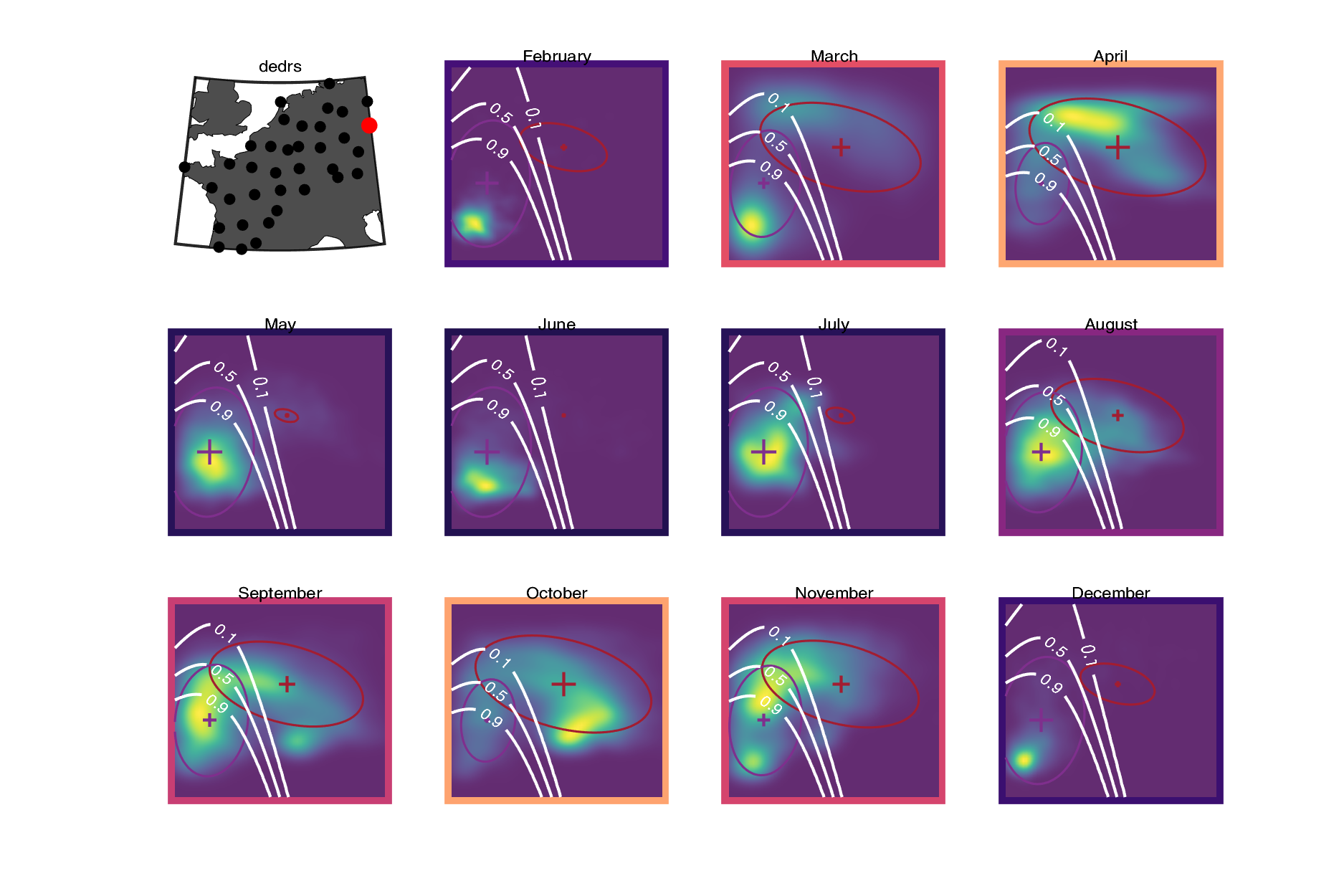

### deeis.png

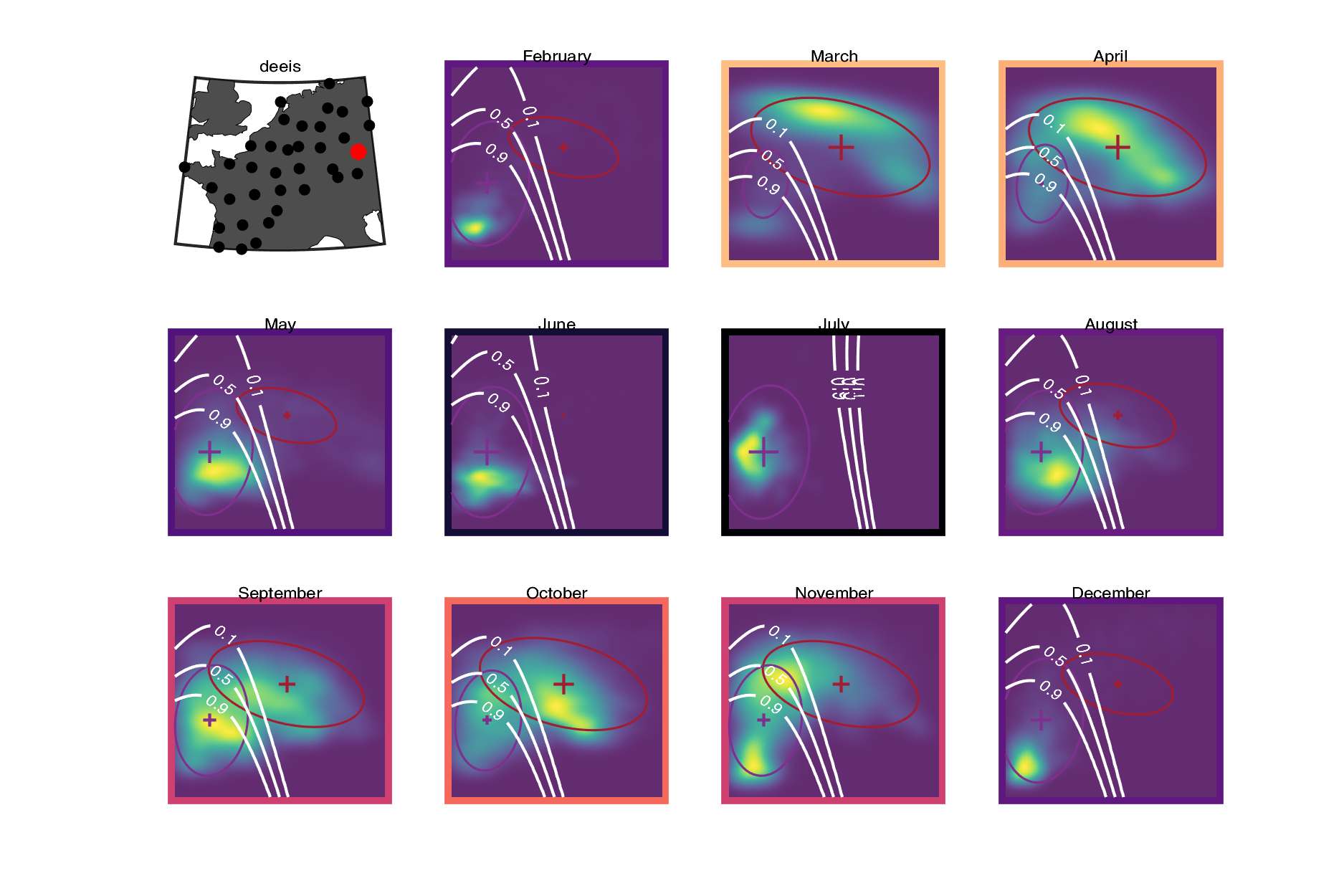

### deess.png

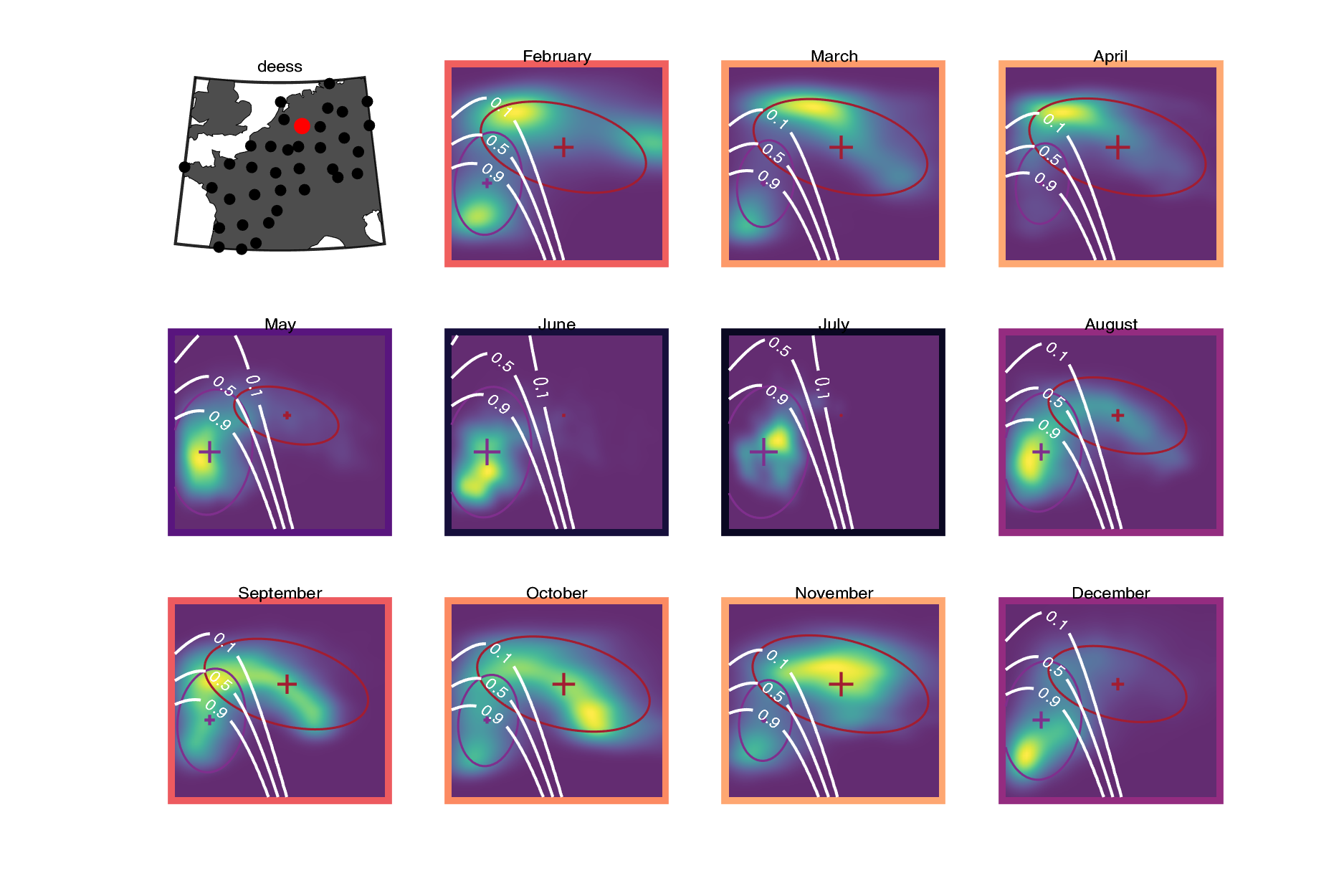

### defld.png

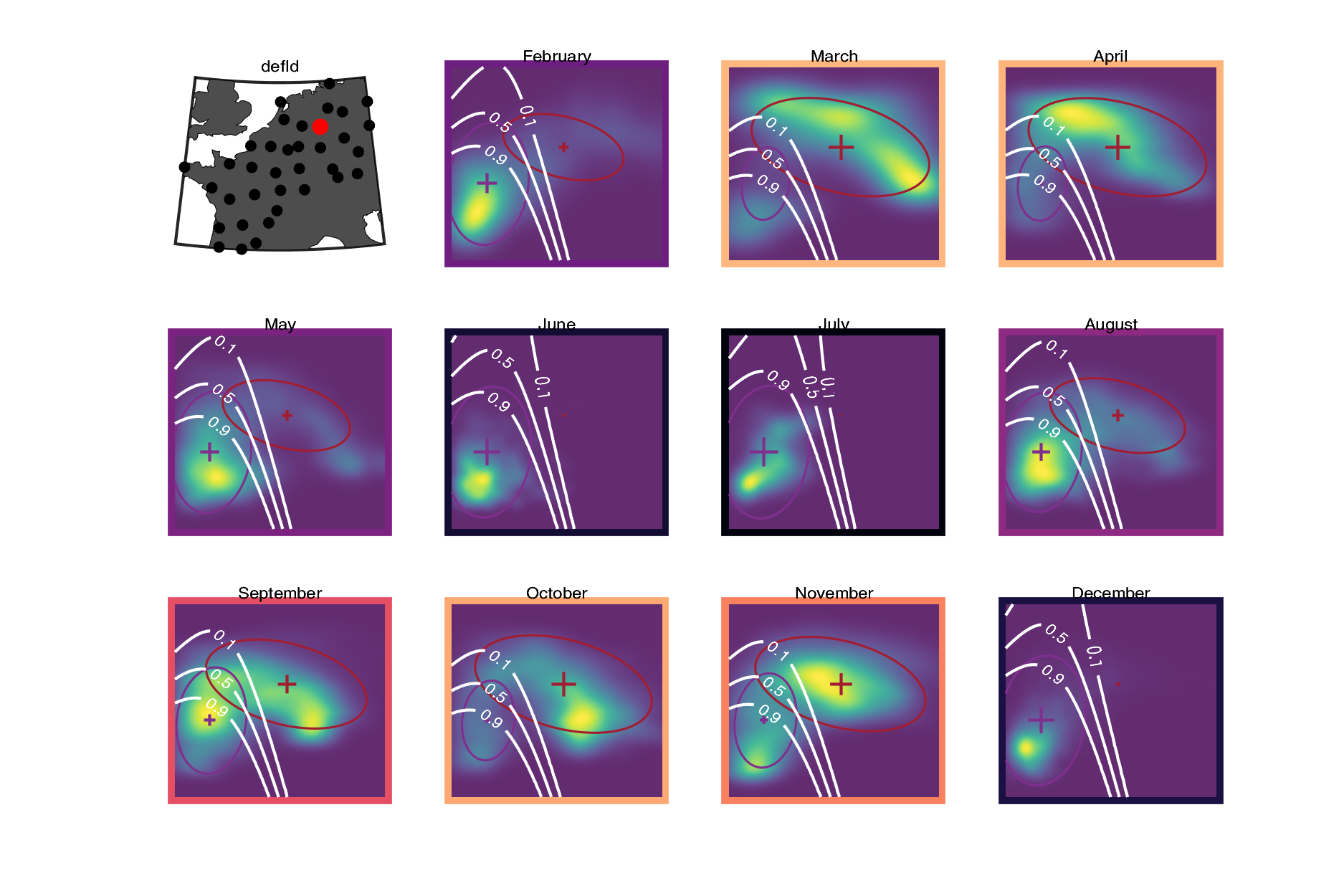

### dehnr.png

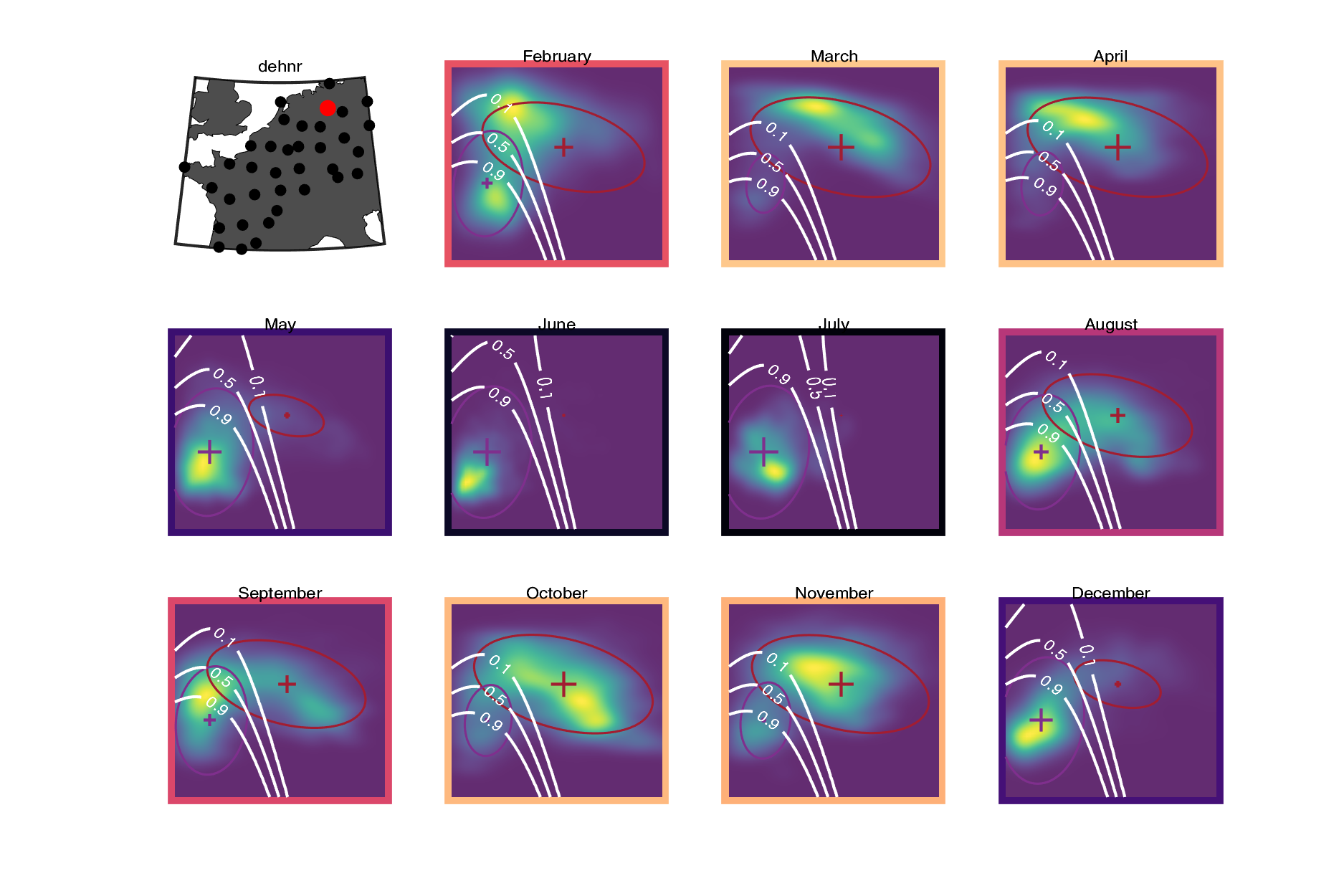

### demem.png

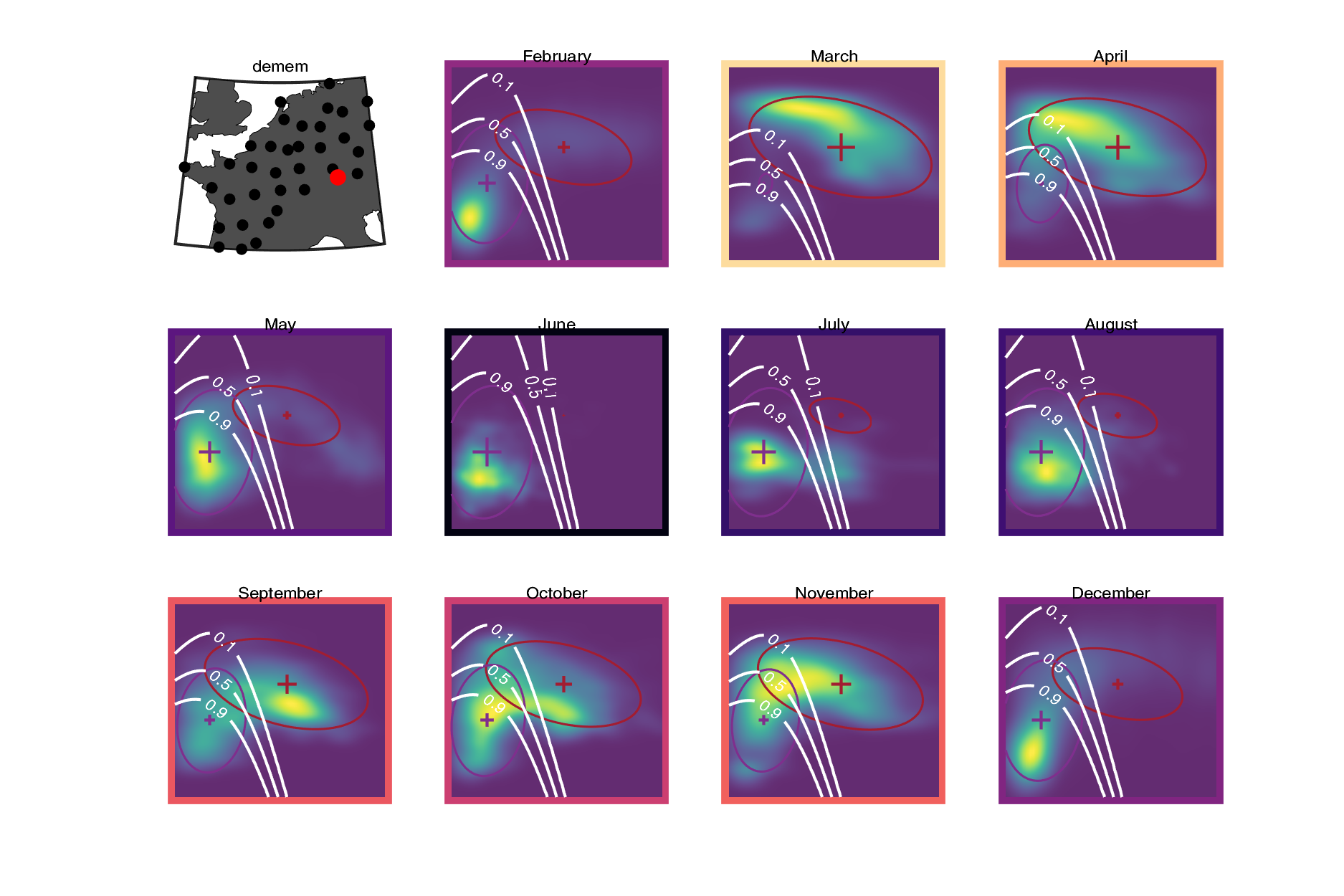

### deneu.png

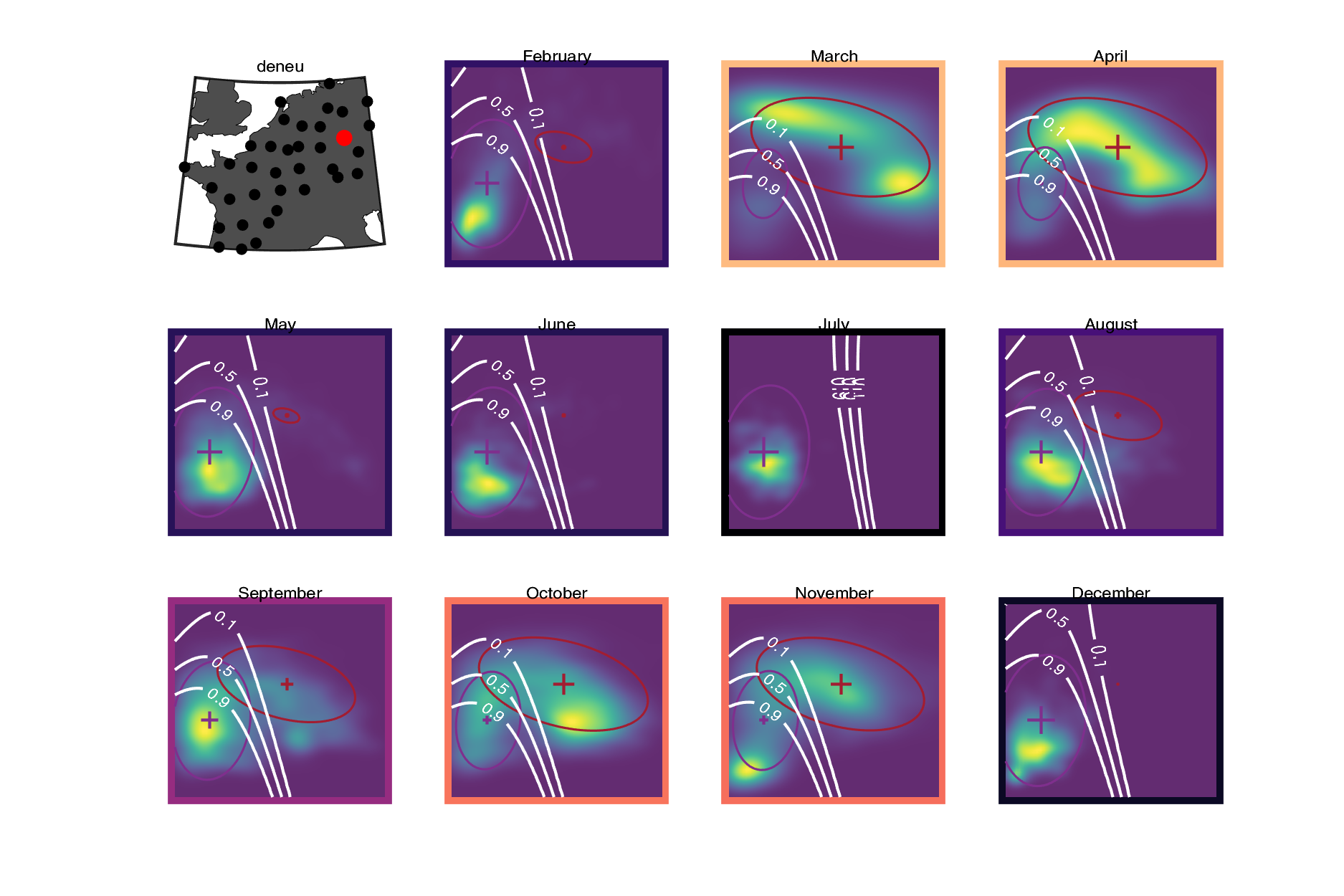

### denhb.png

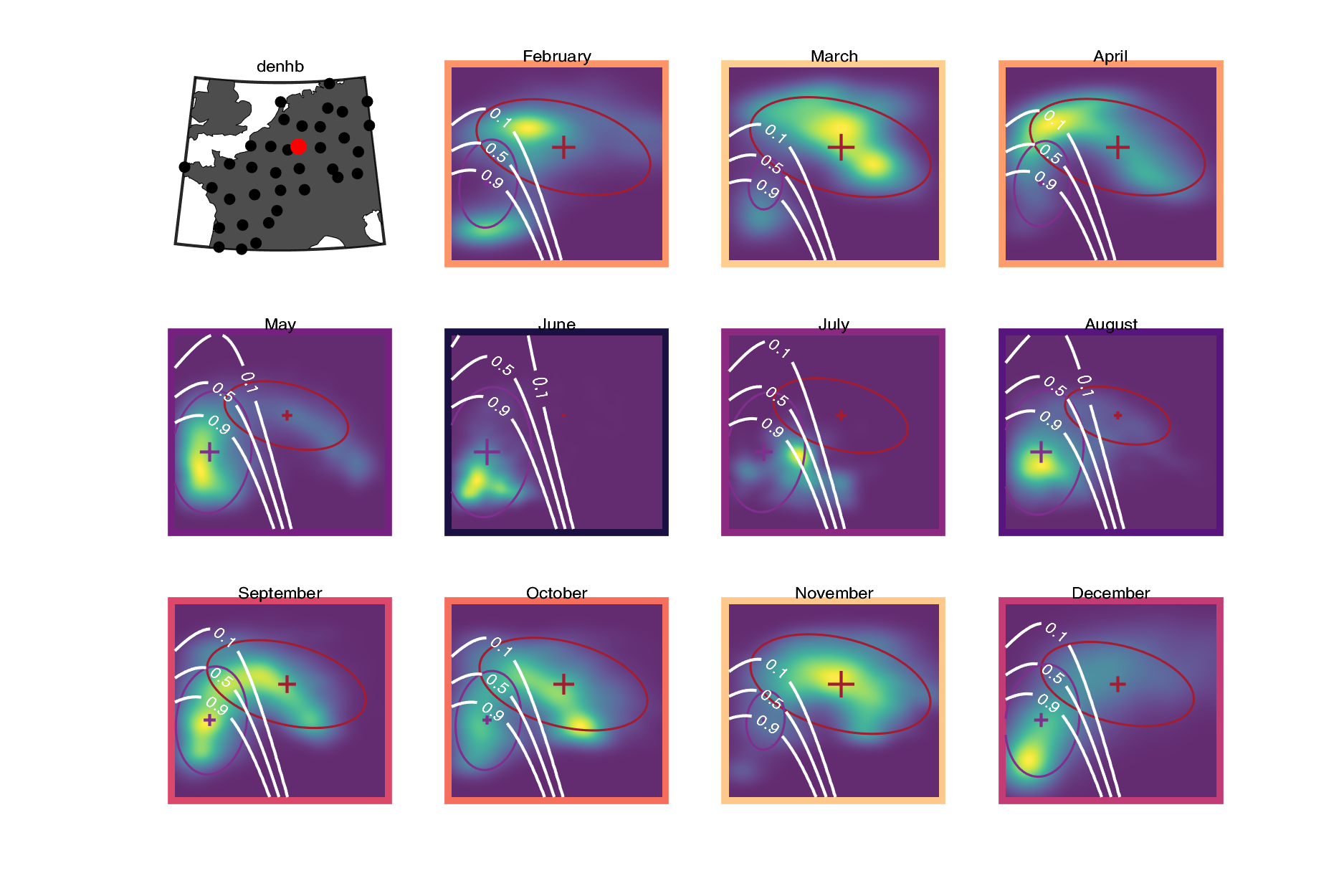

### deoft.png

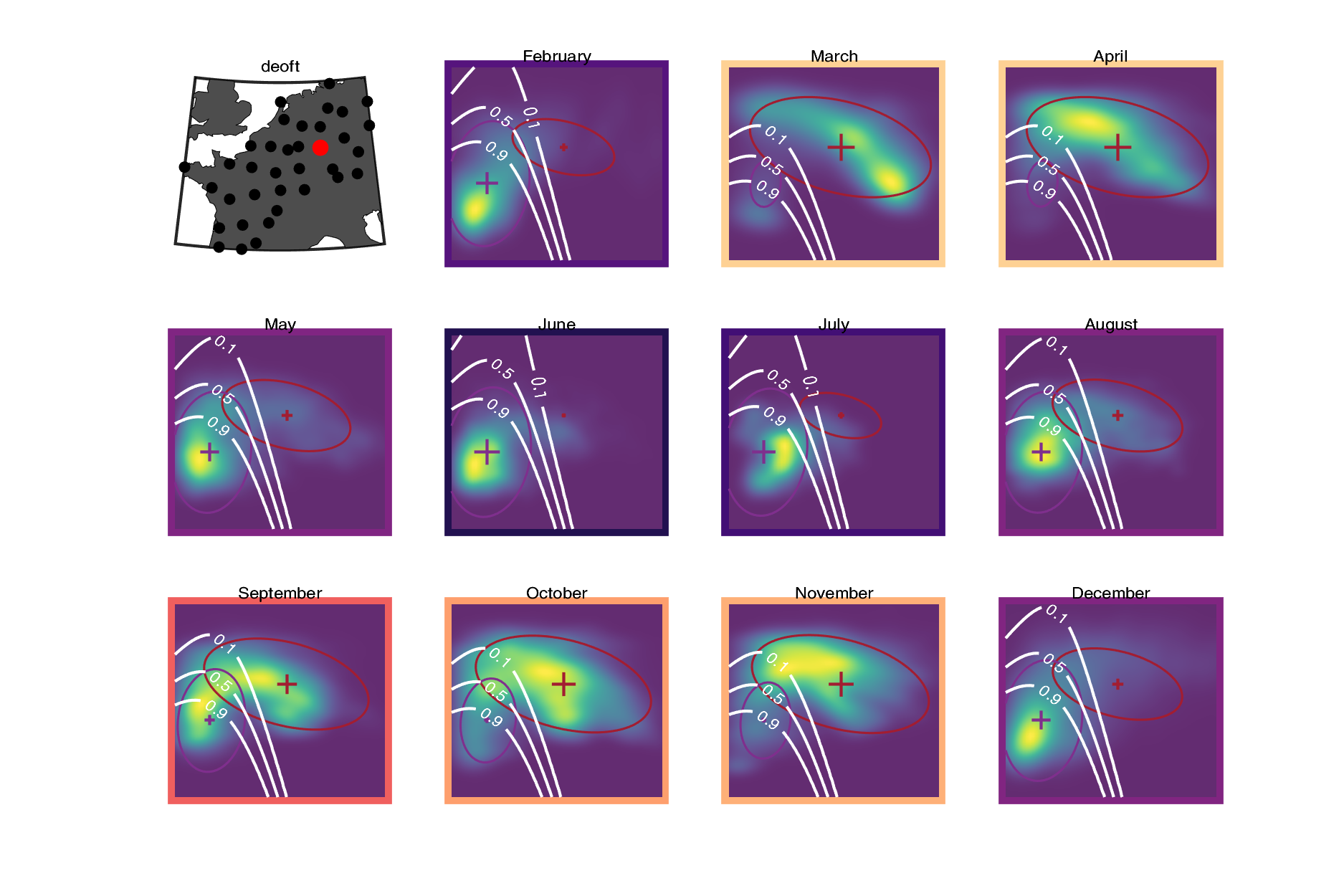

### depro.png

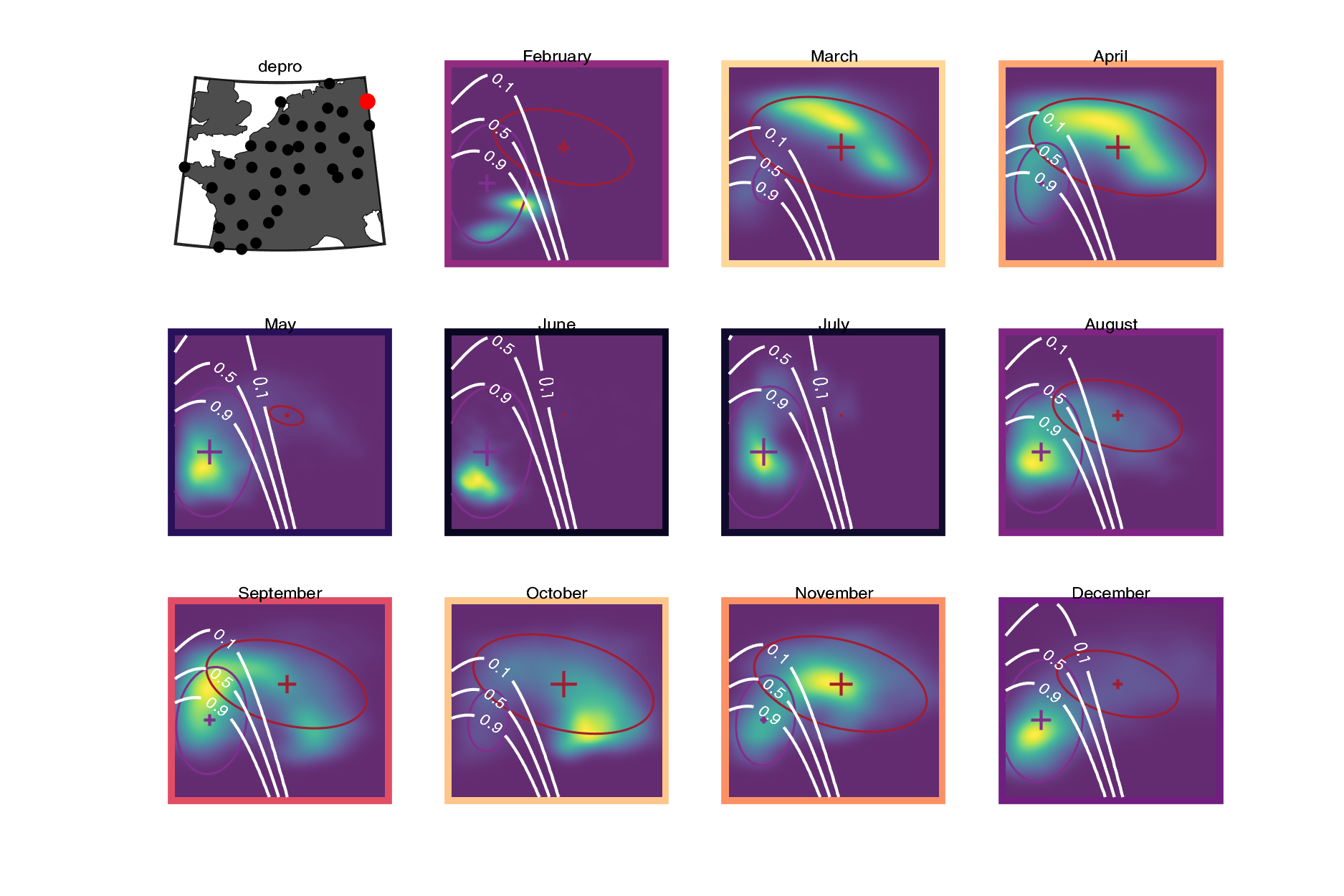

### deros.png

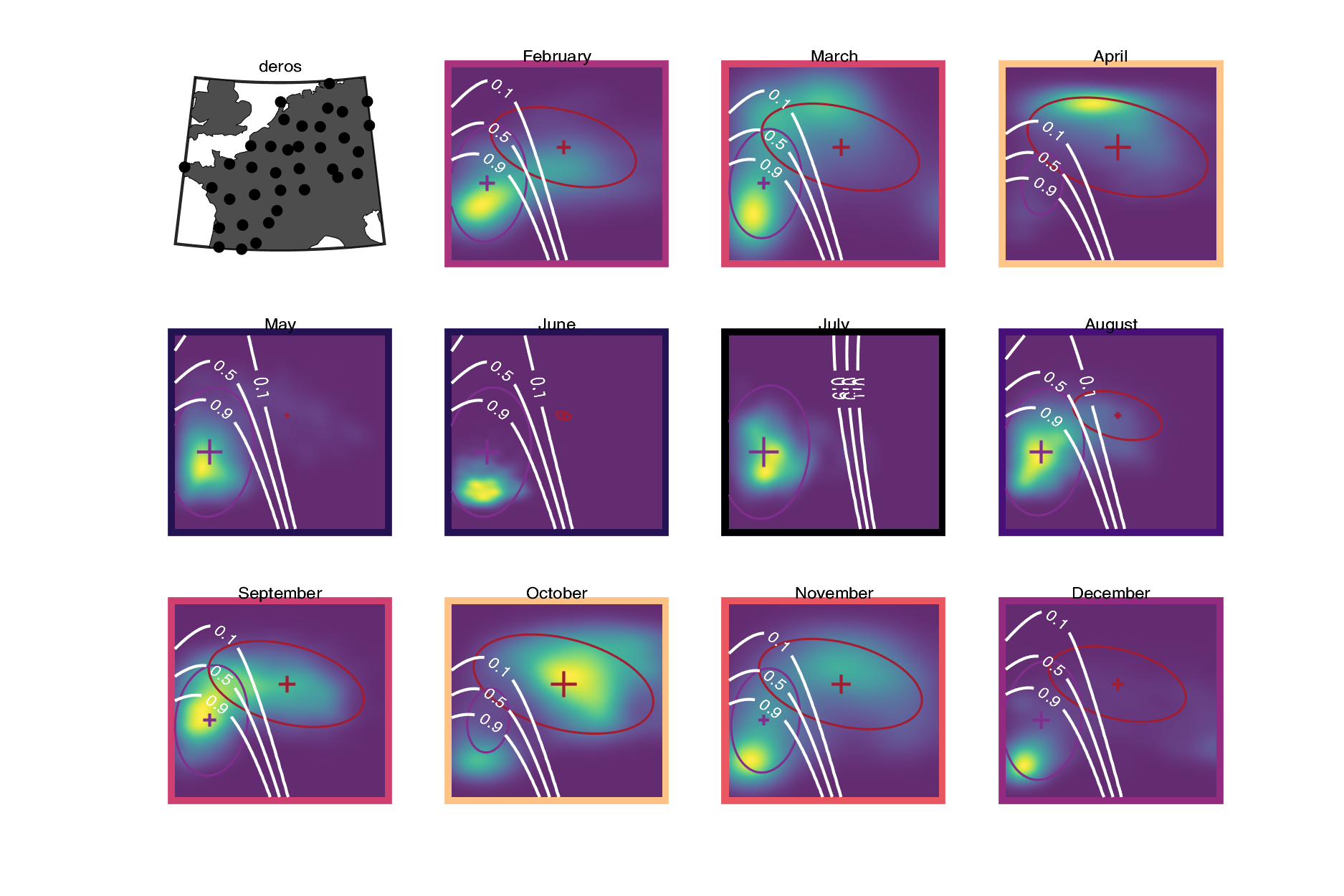

### desna.png

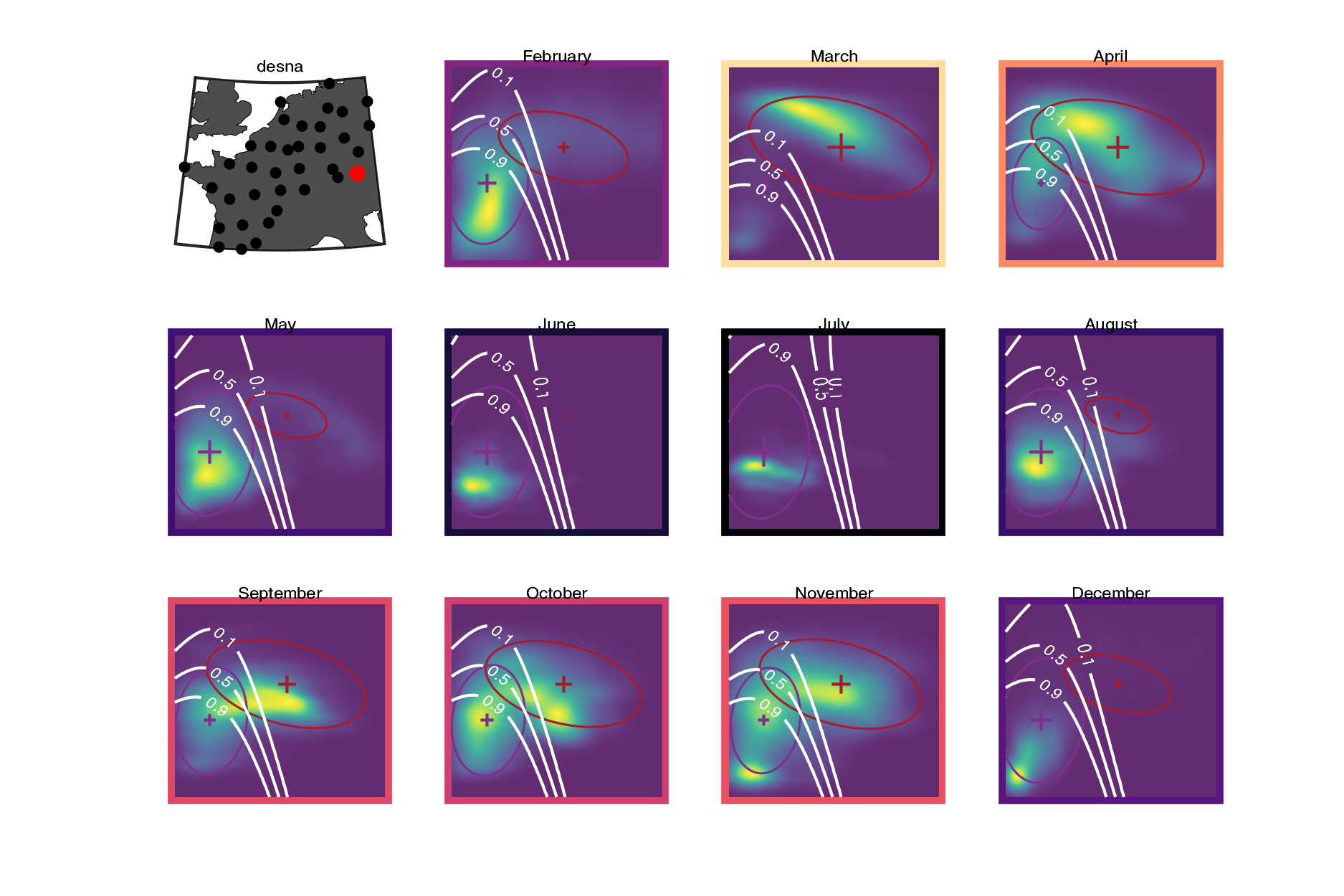

### detur.png

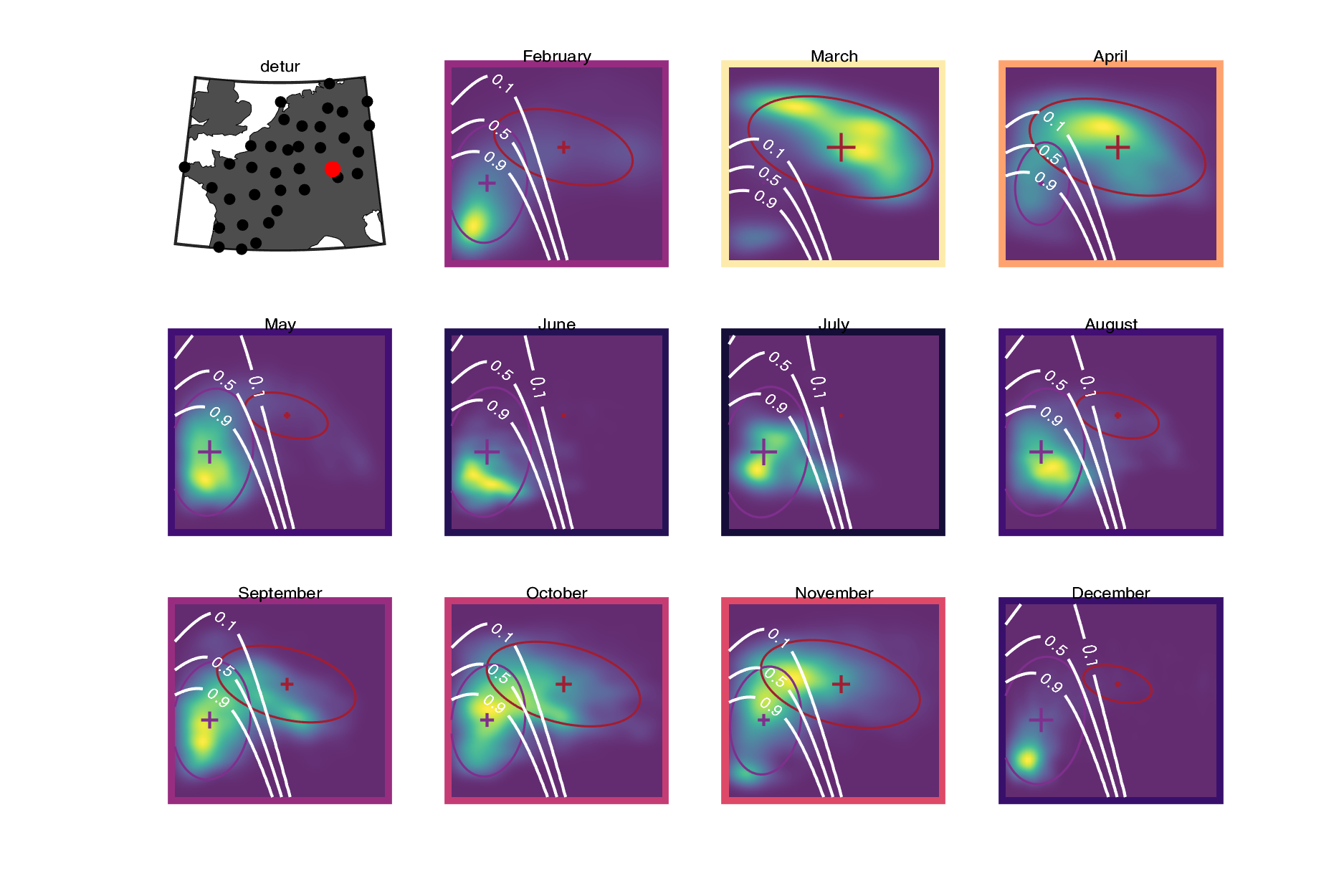

### deumd.png

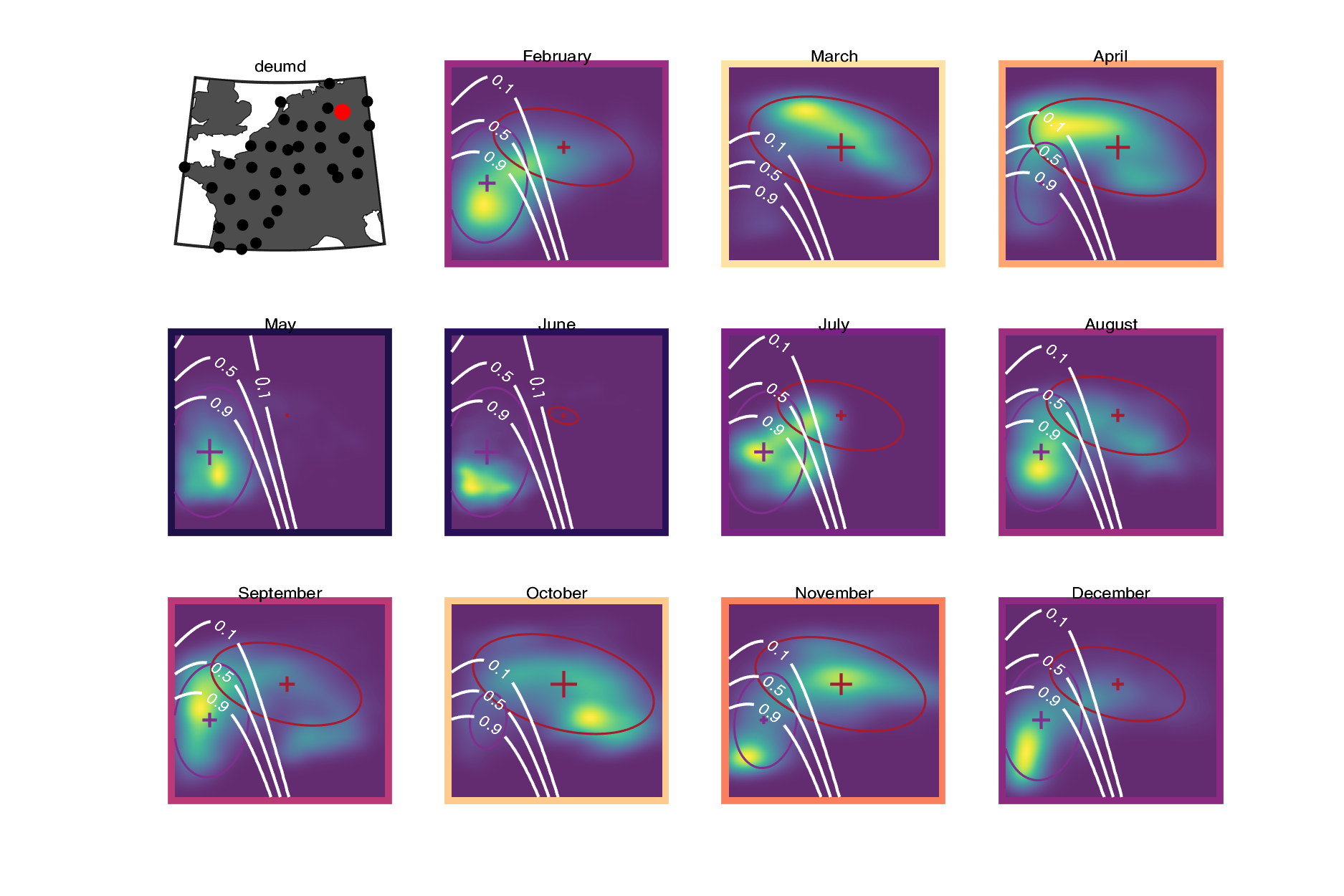

### frabb.png

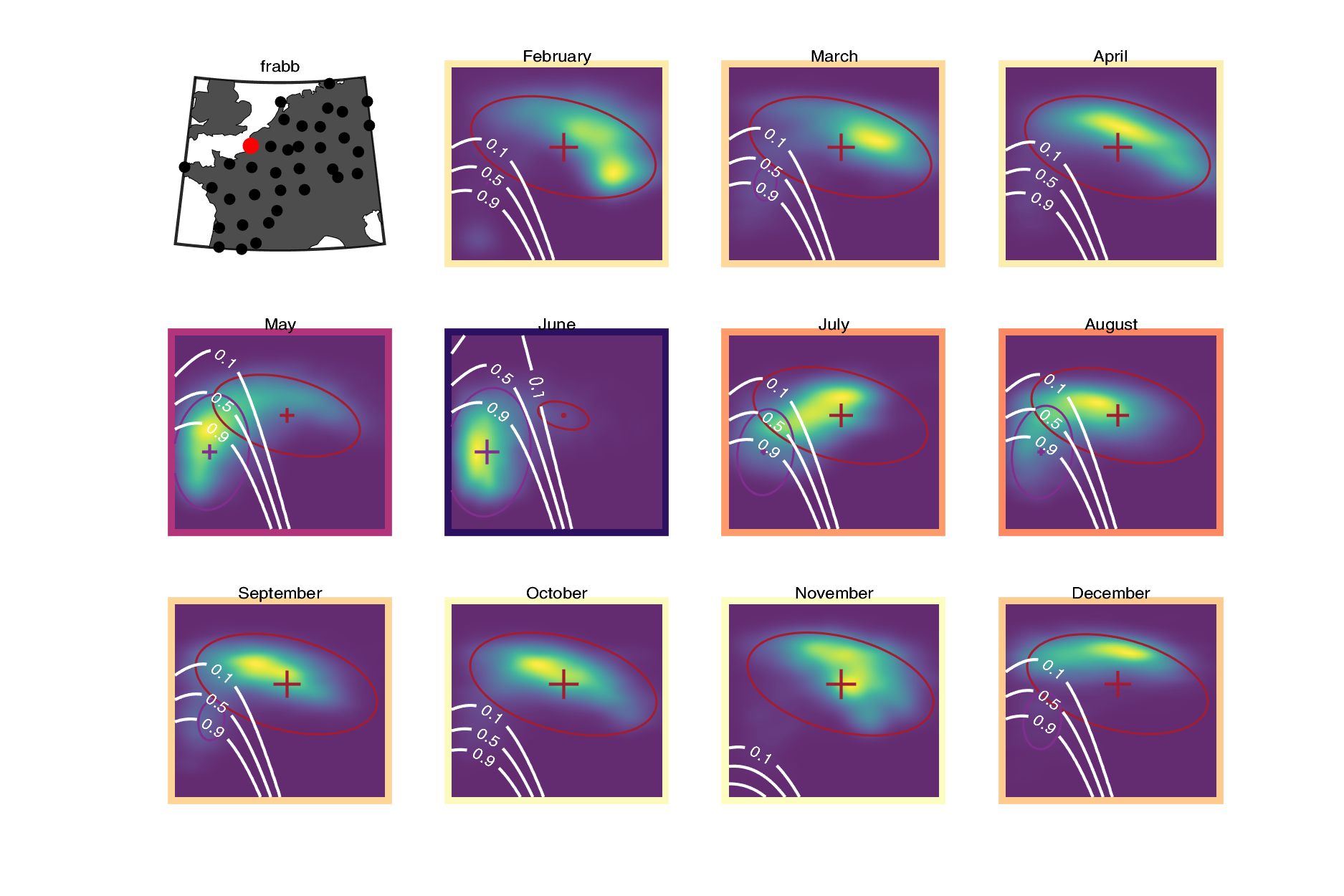

### frave.png

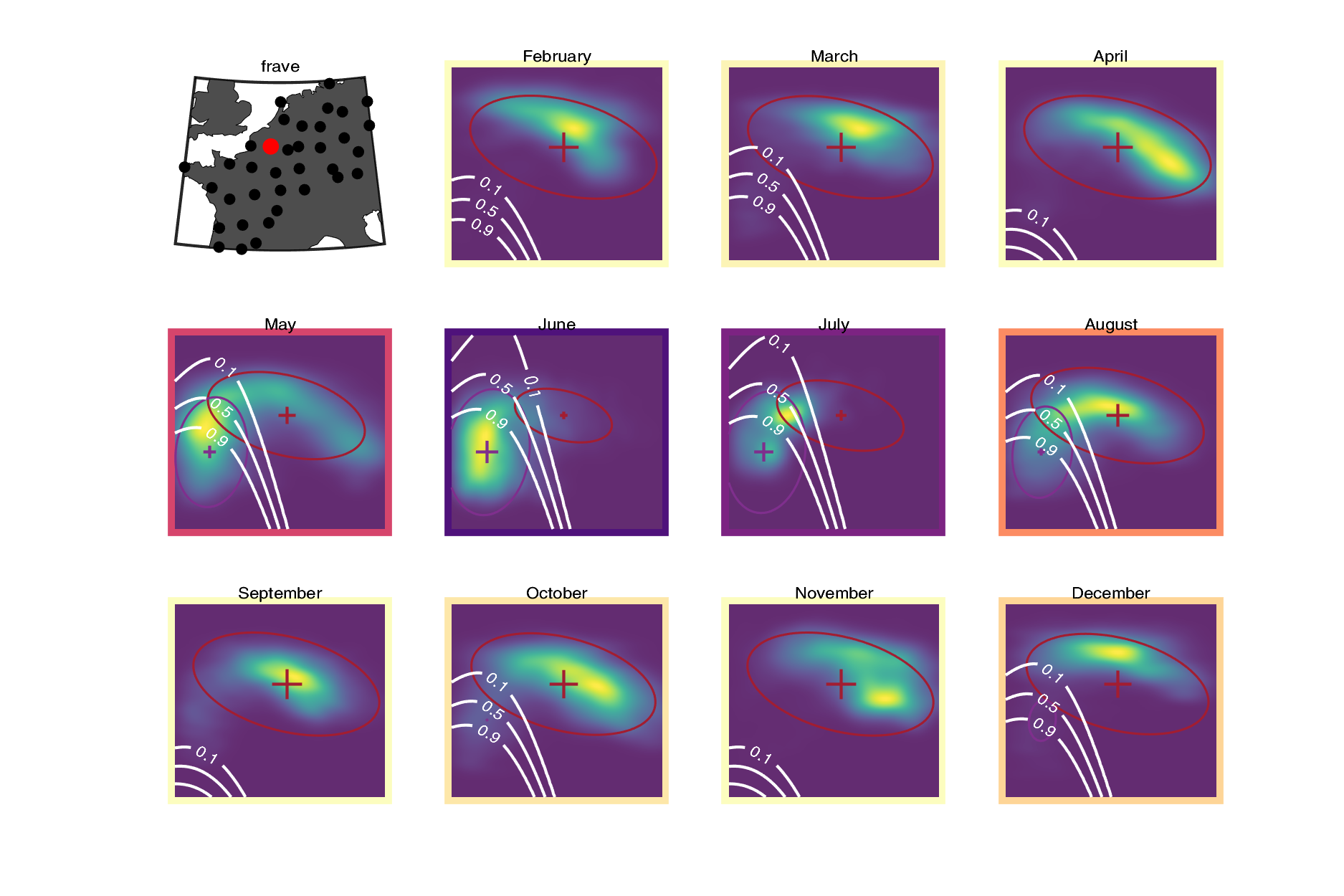

### frbla.png

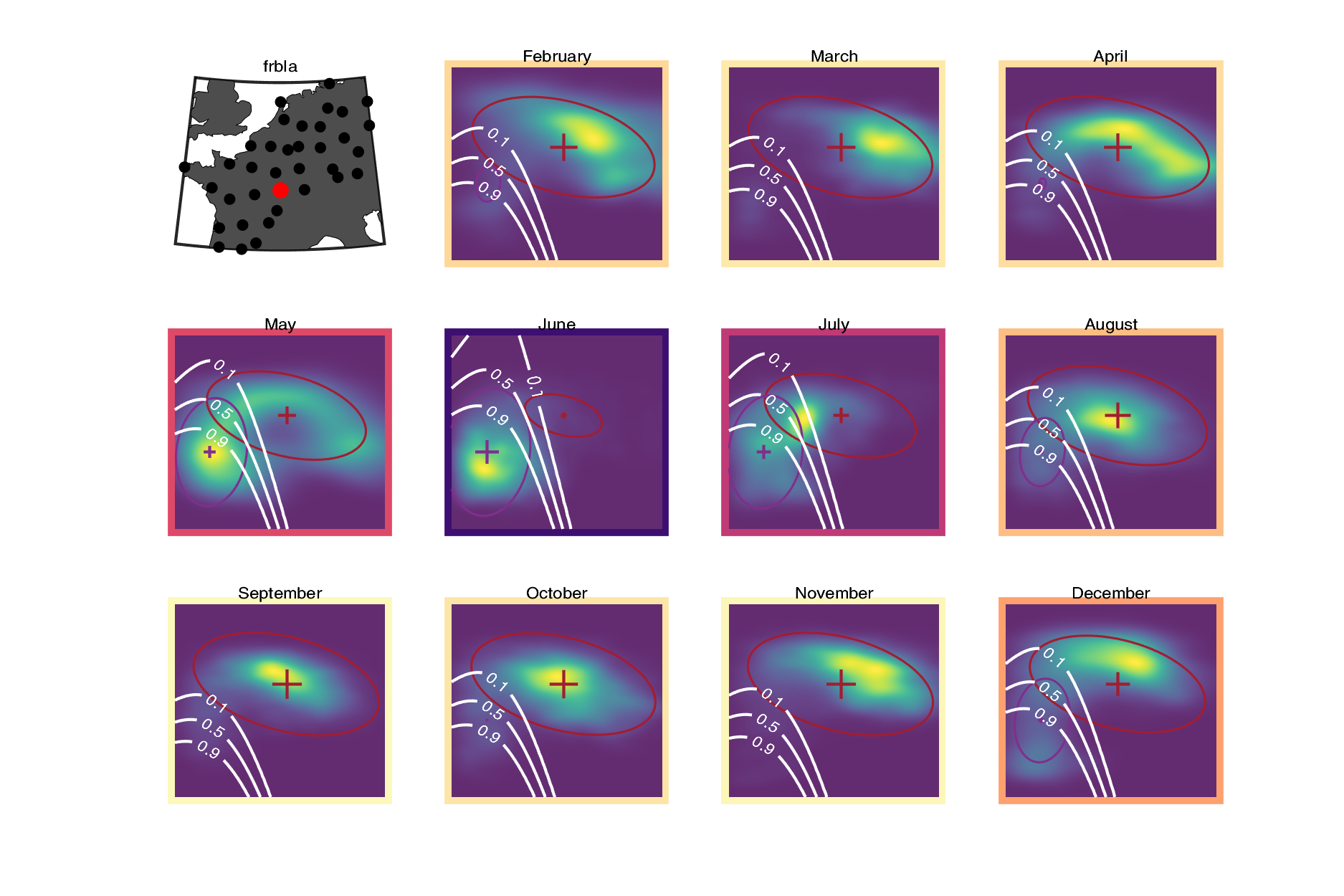

### frbor.png

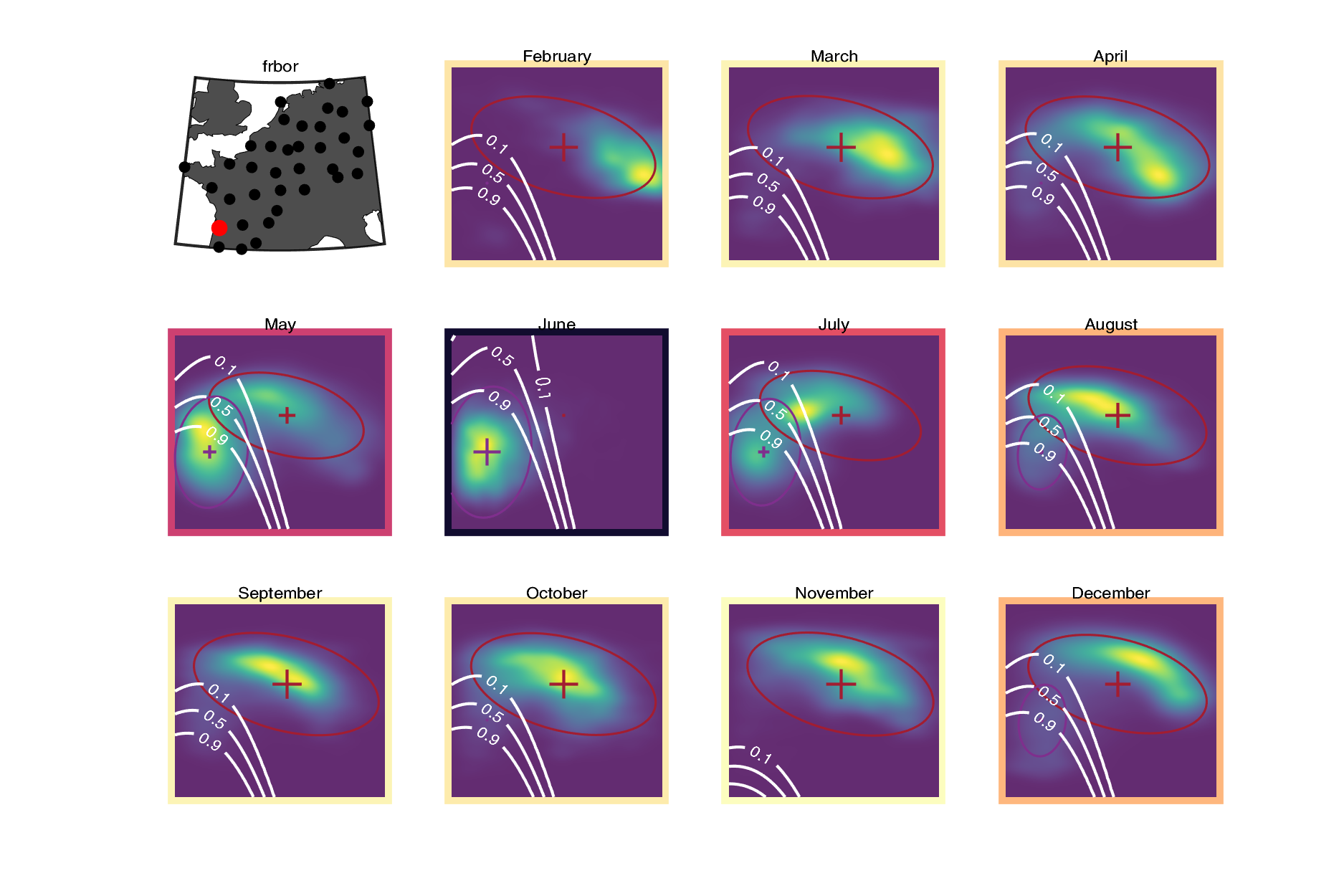

### frbou.png

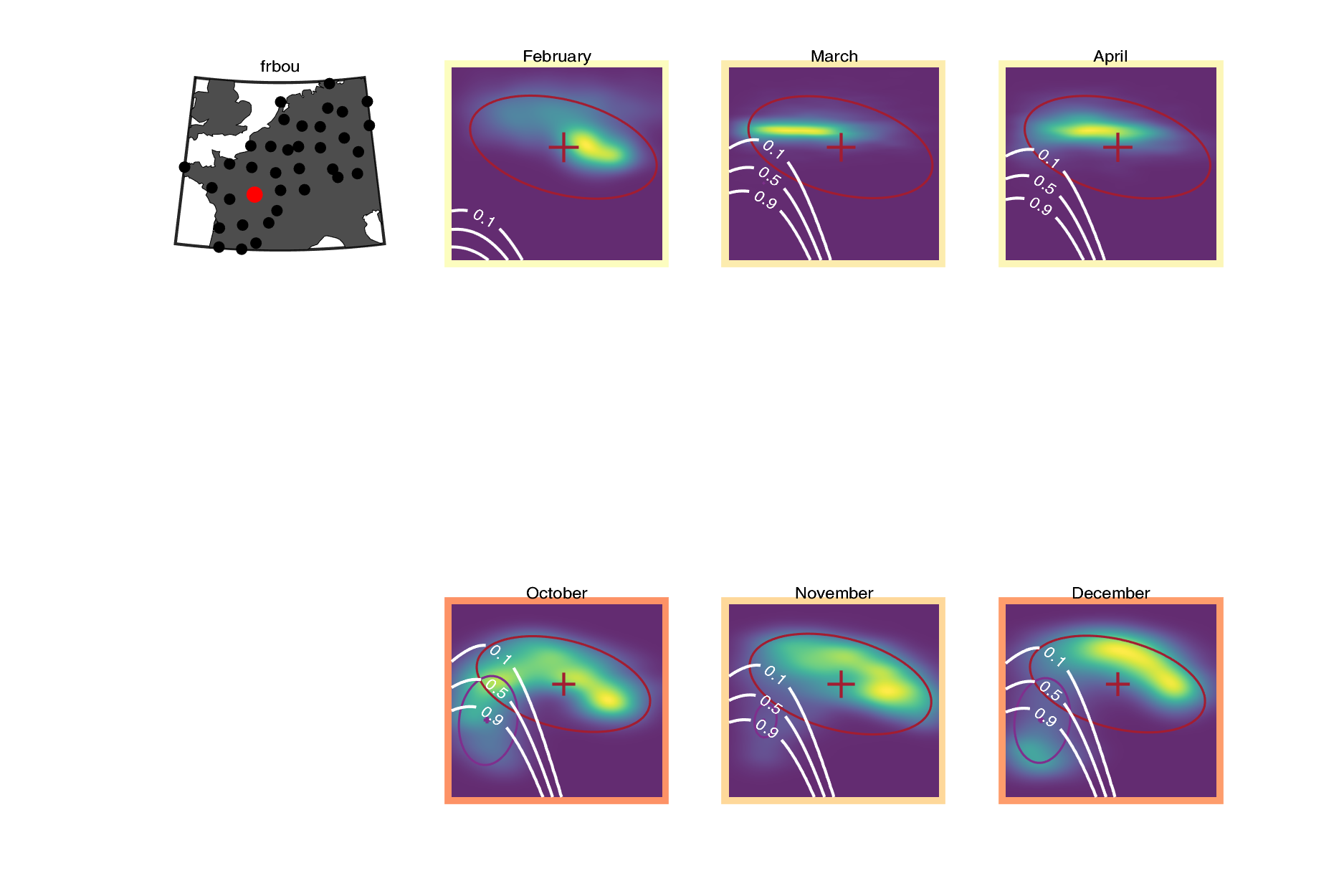

### frcae.png

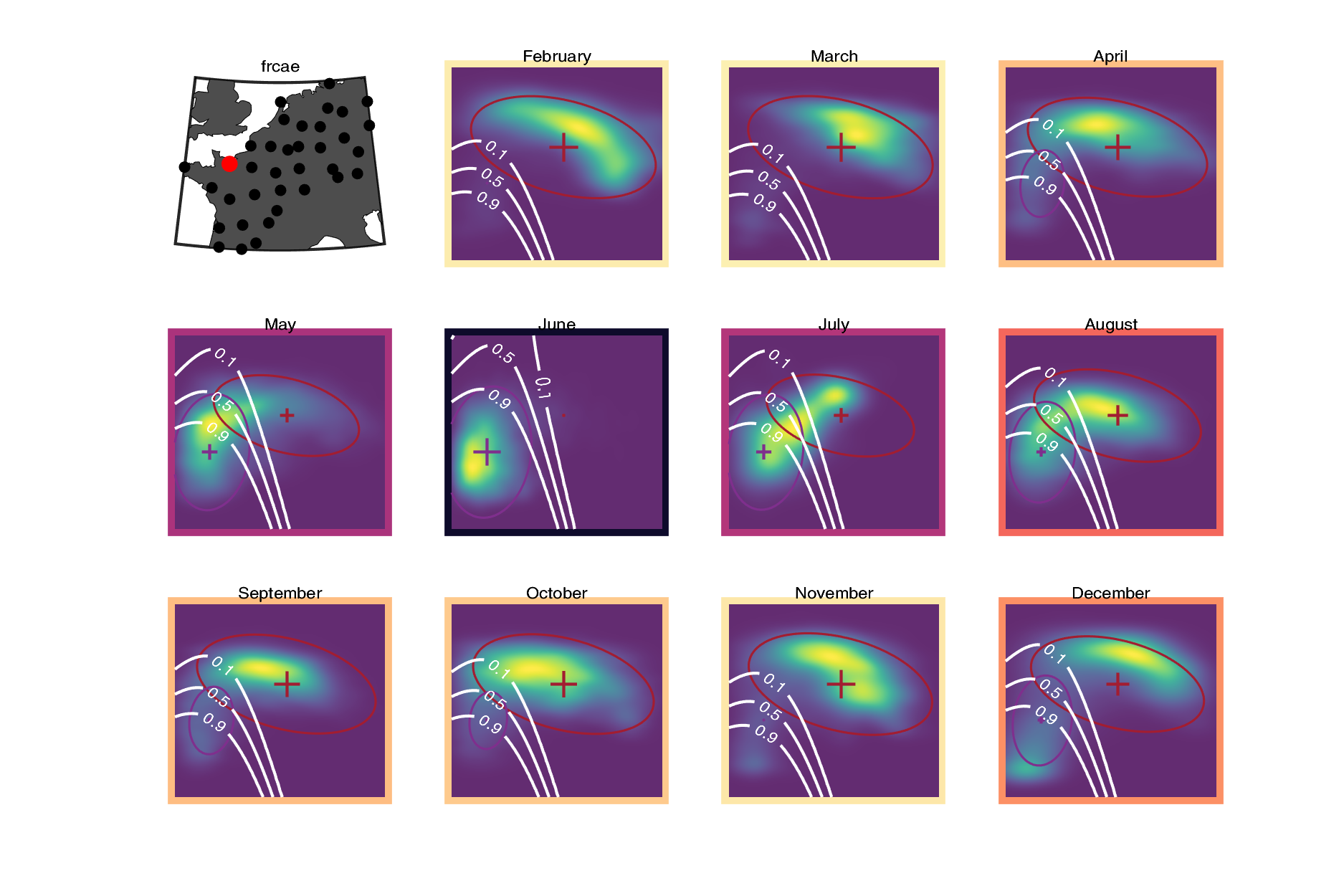

### frche.png

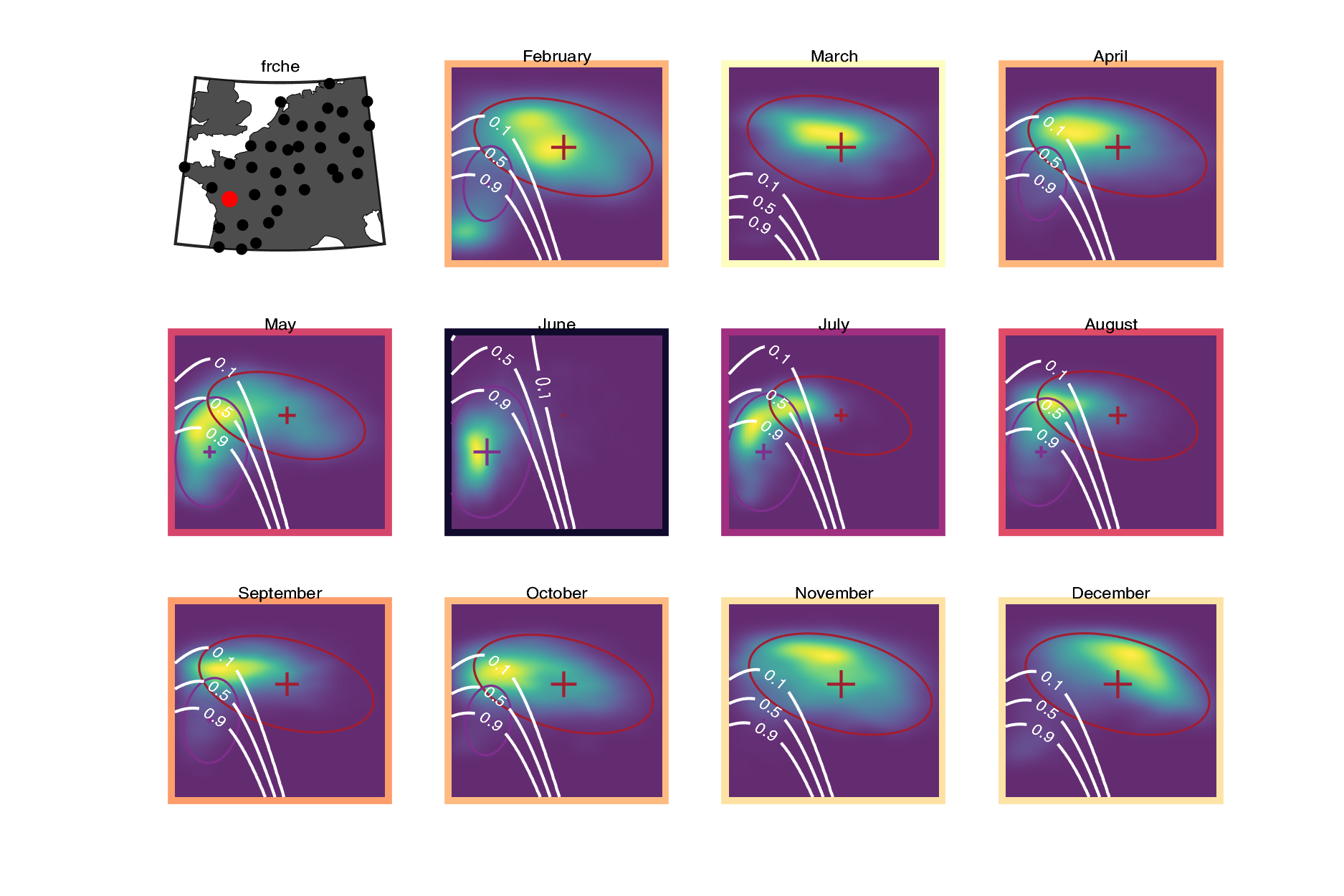

### frgre.png

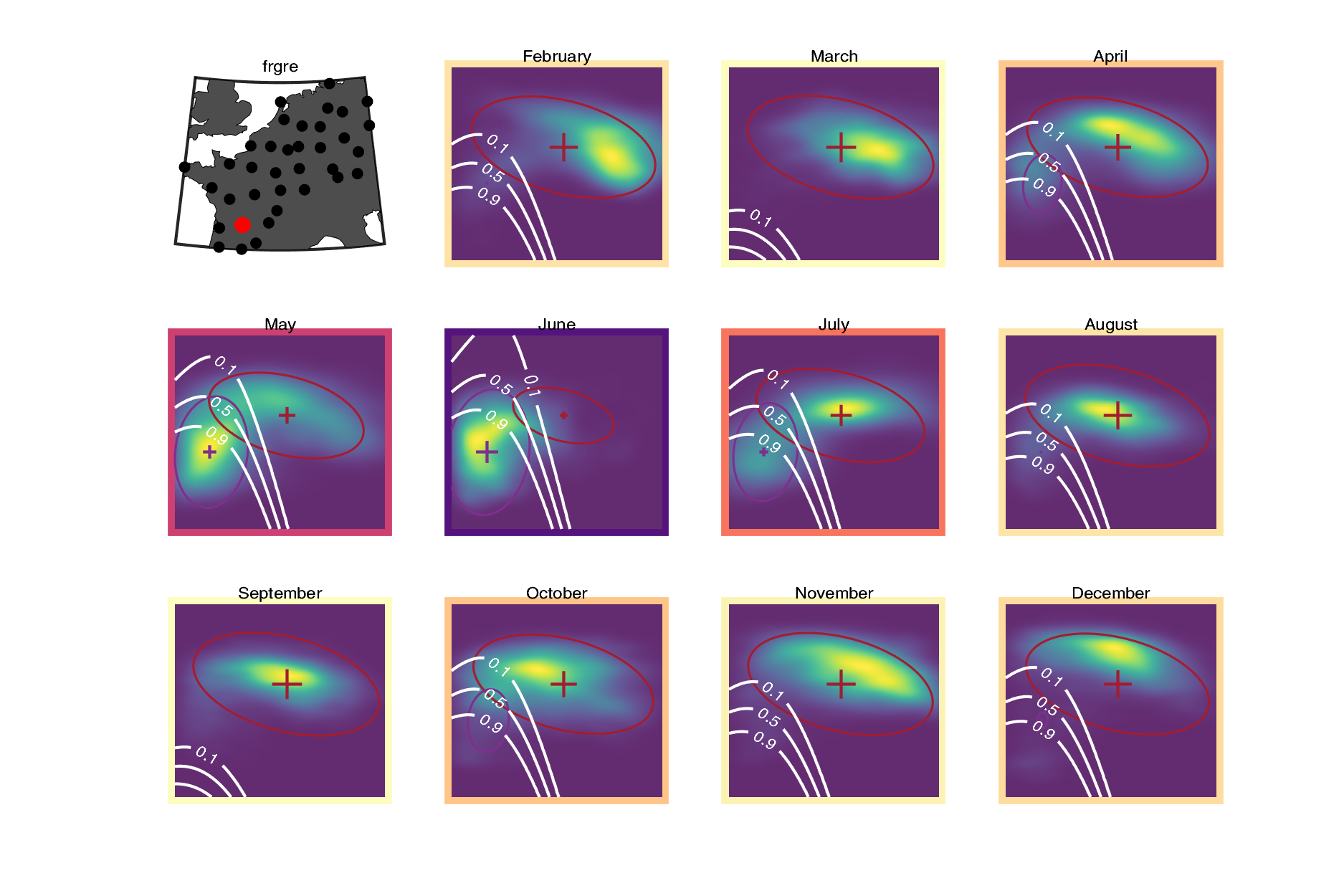

### frlep.png

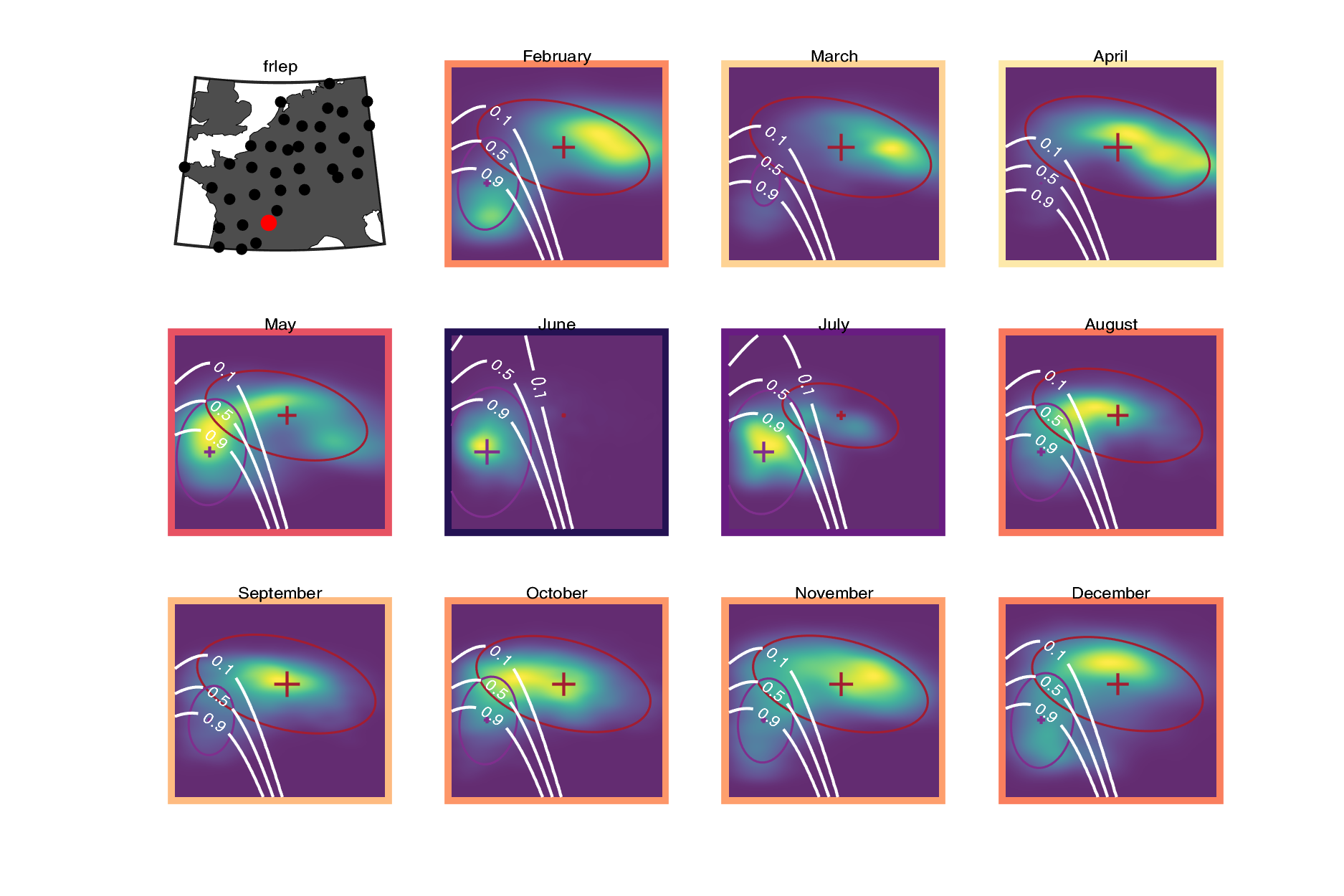

### frmcl.png

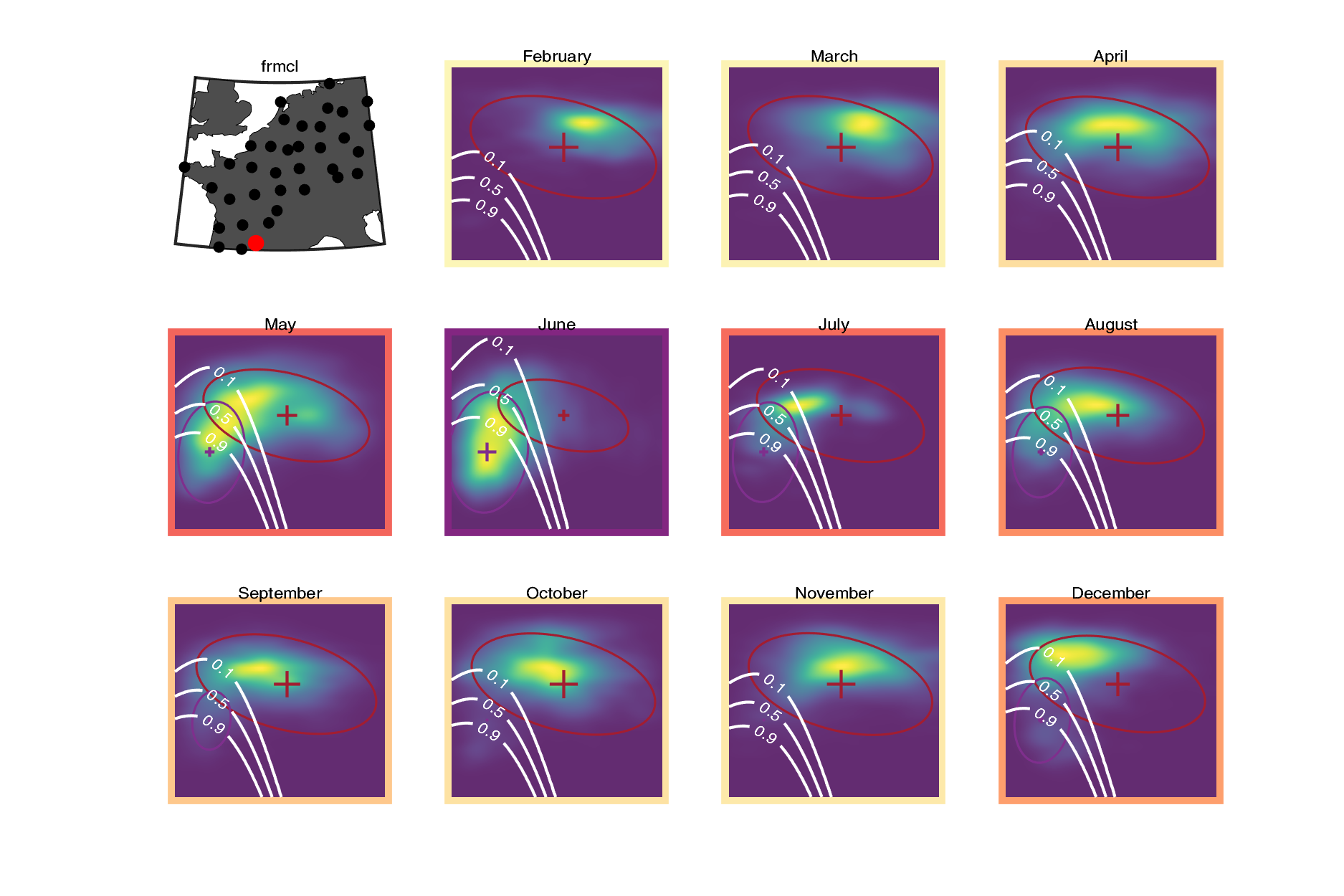

### frmom.png

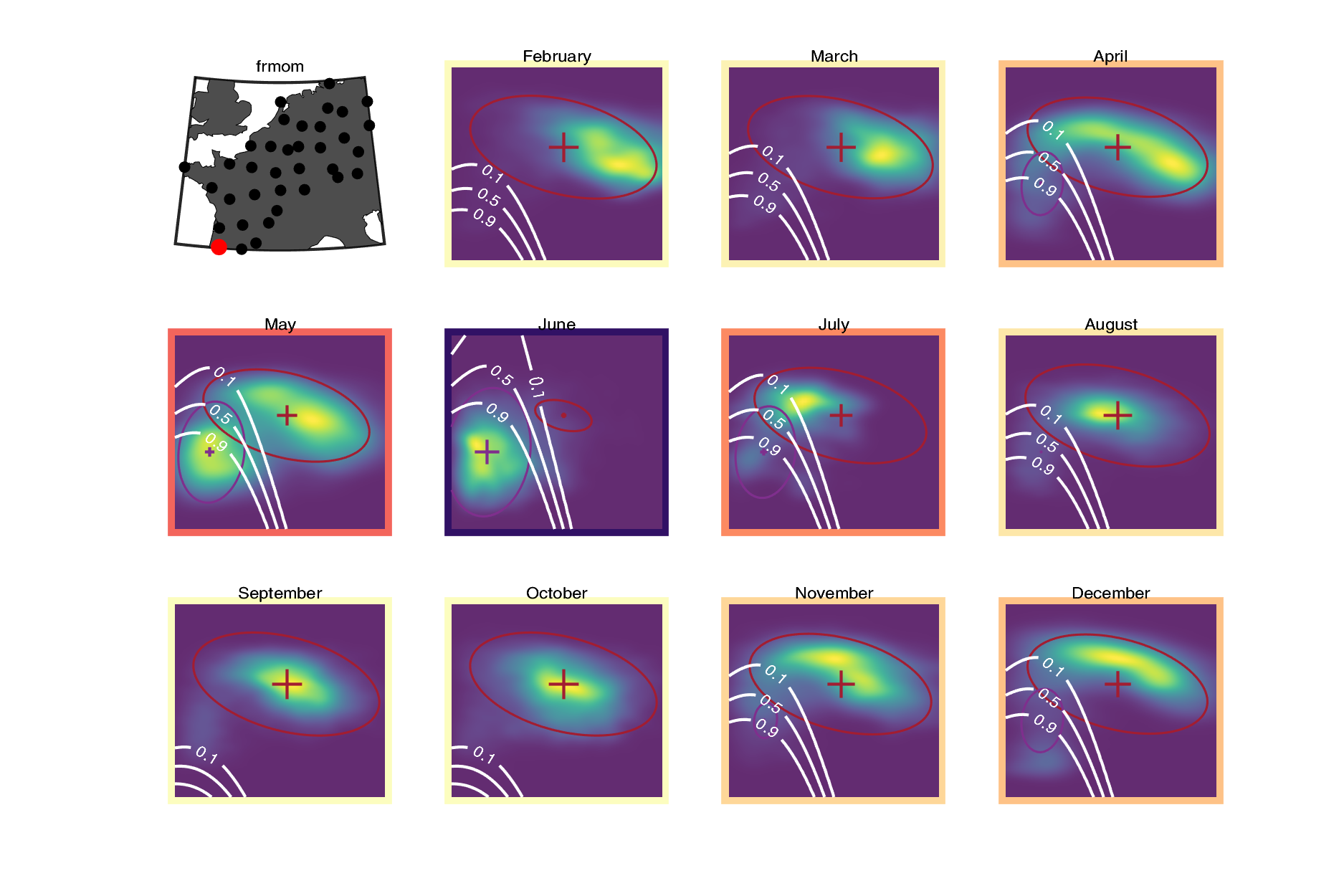

### frmtc.png

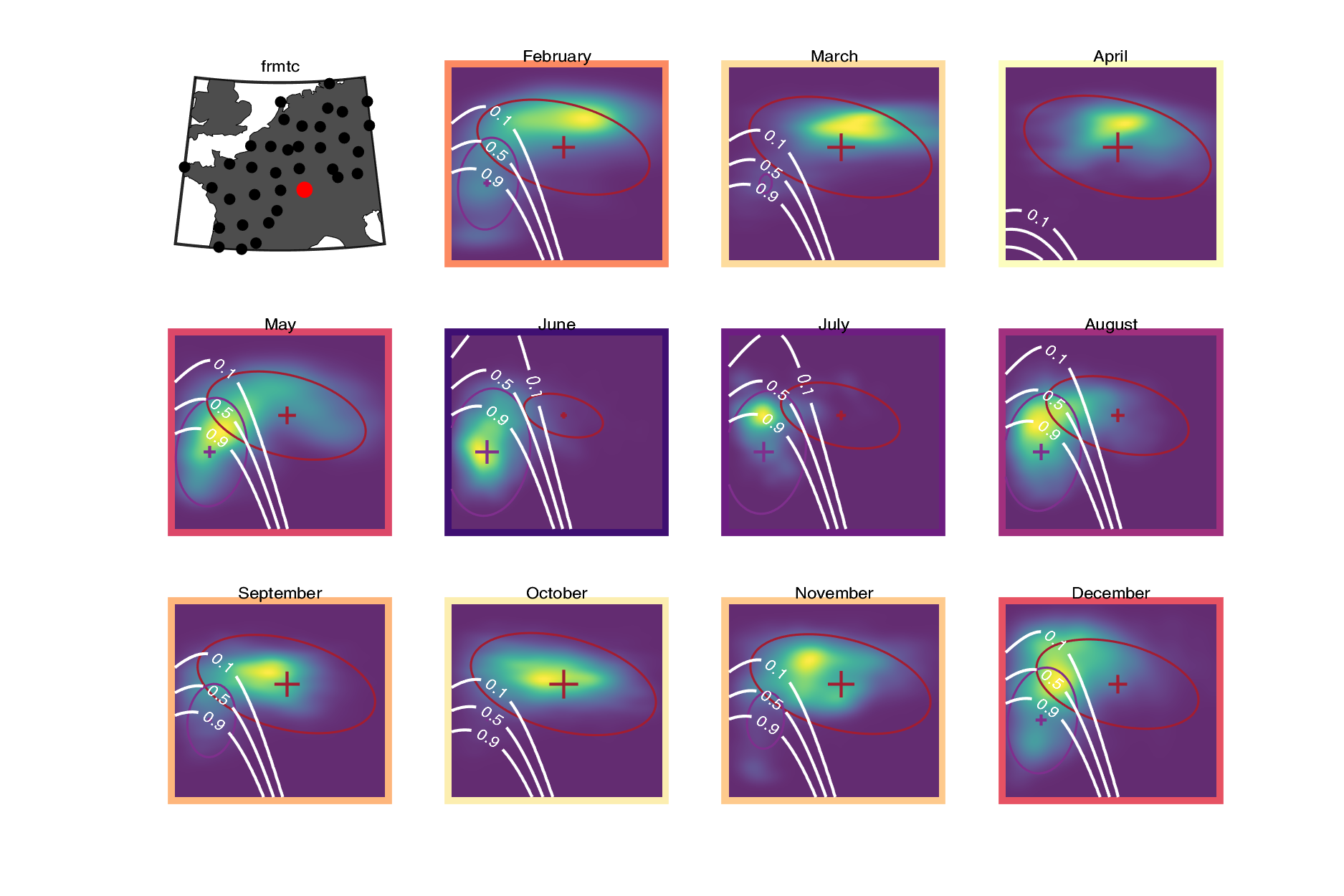

### frnan.png

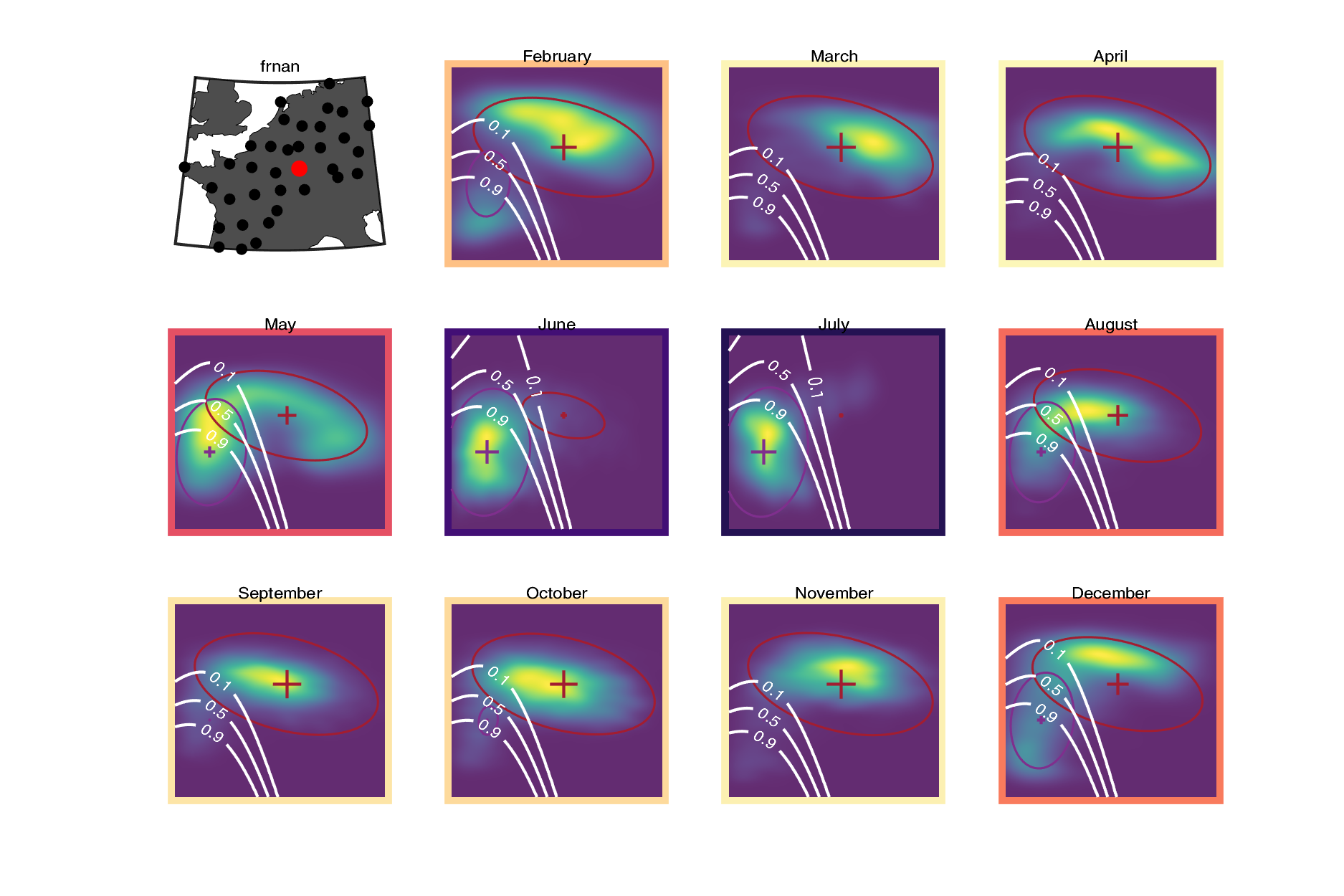

### frniz.png

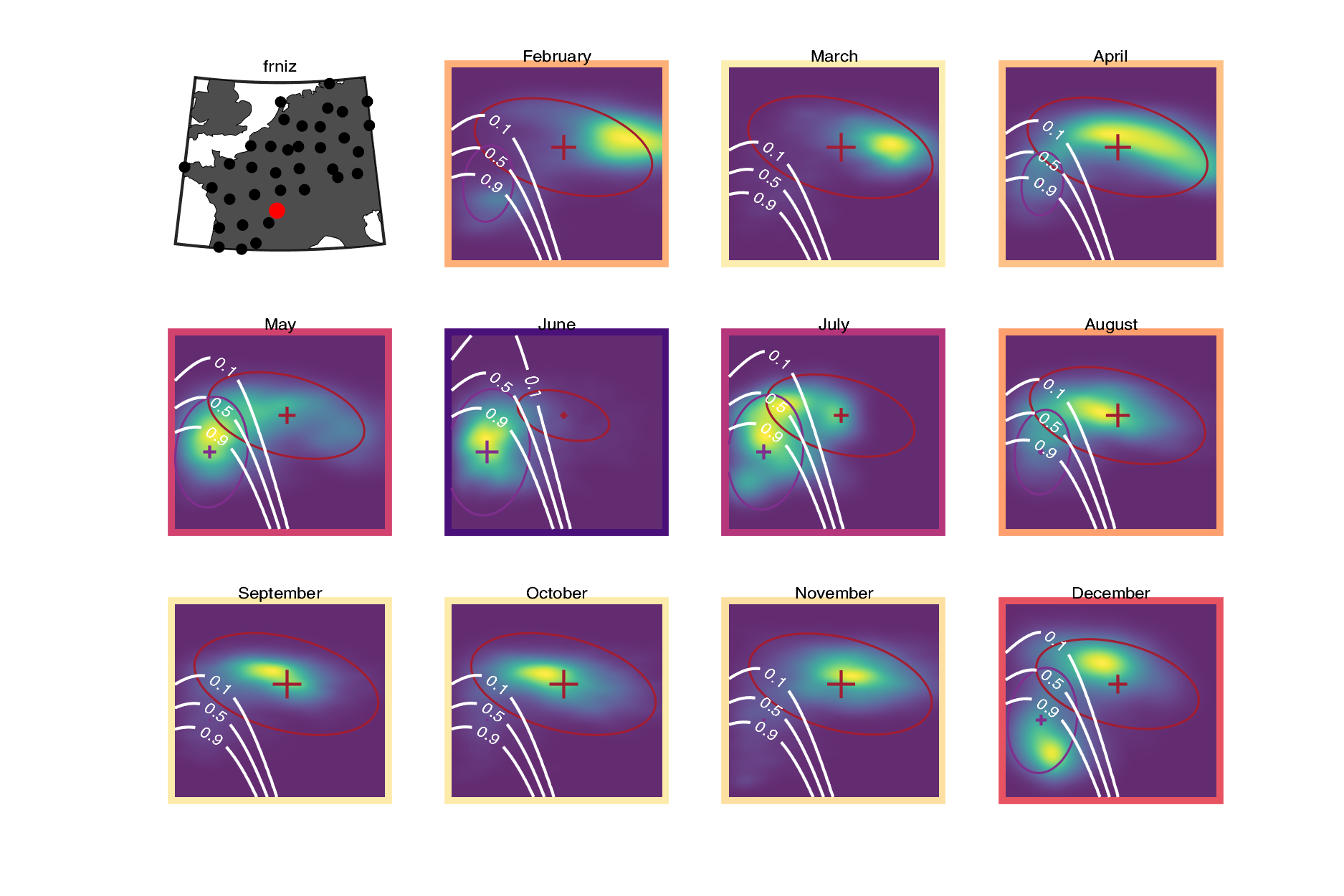

### frpla.png

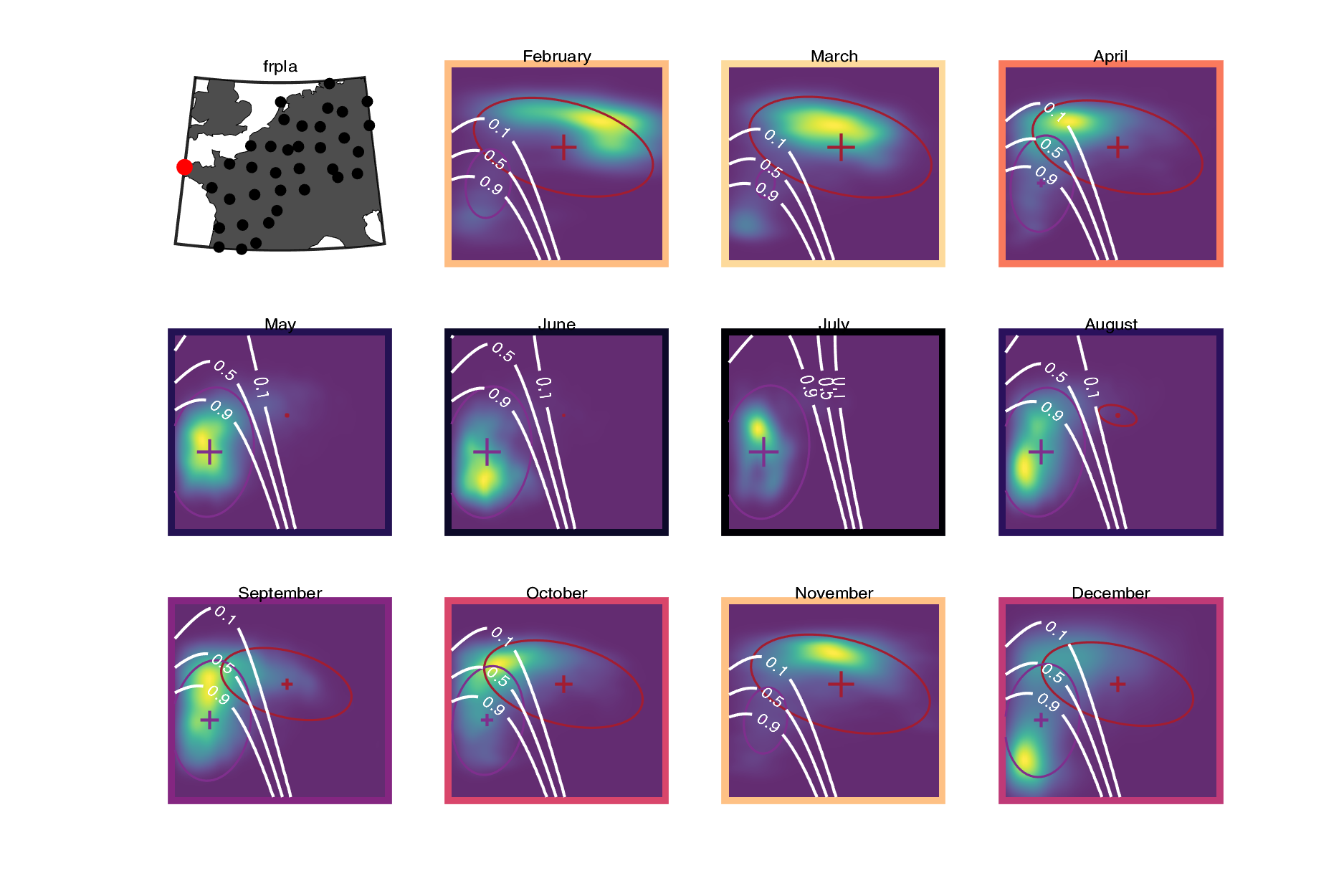

### frtou.png

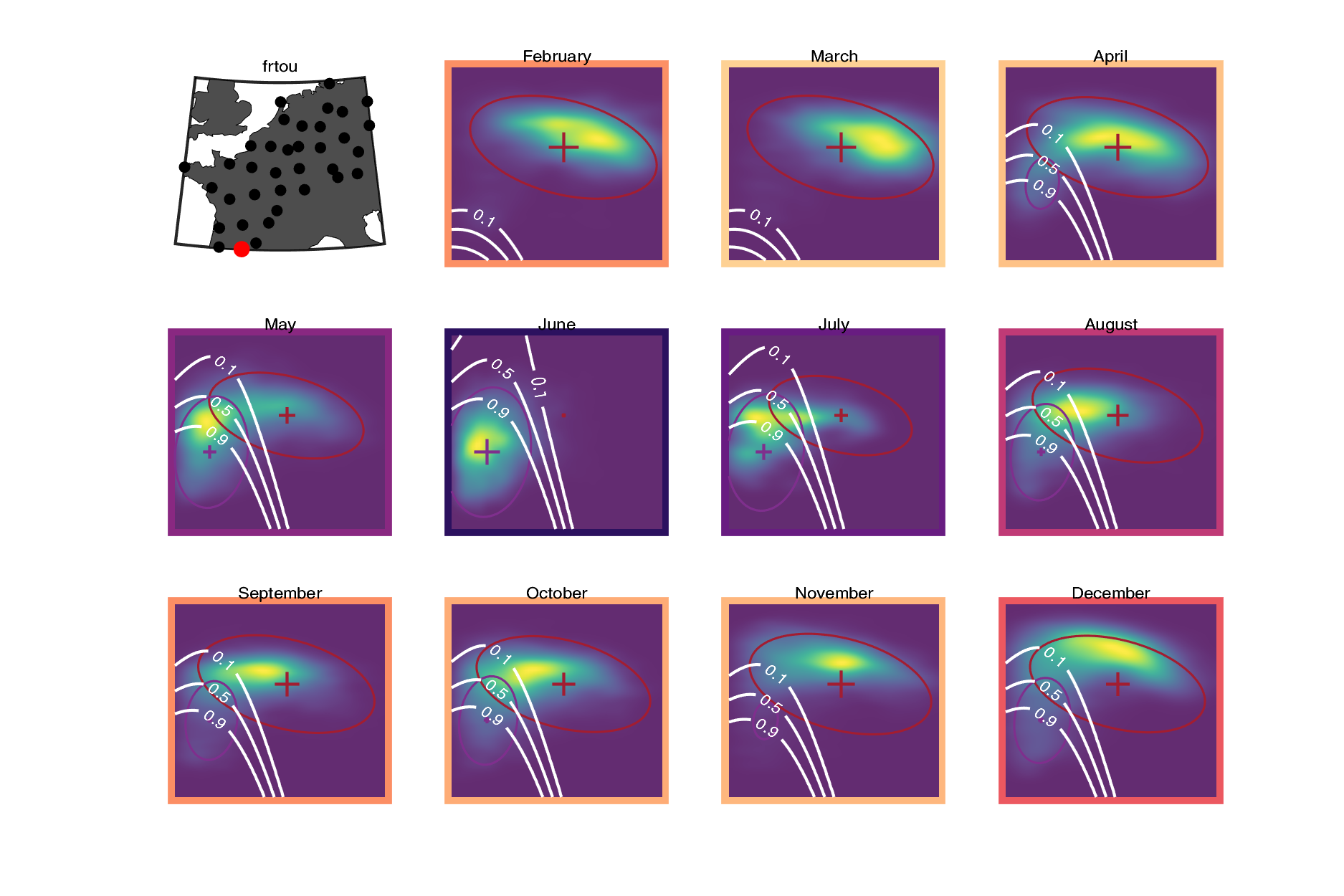

### frtra.png

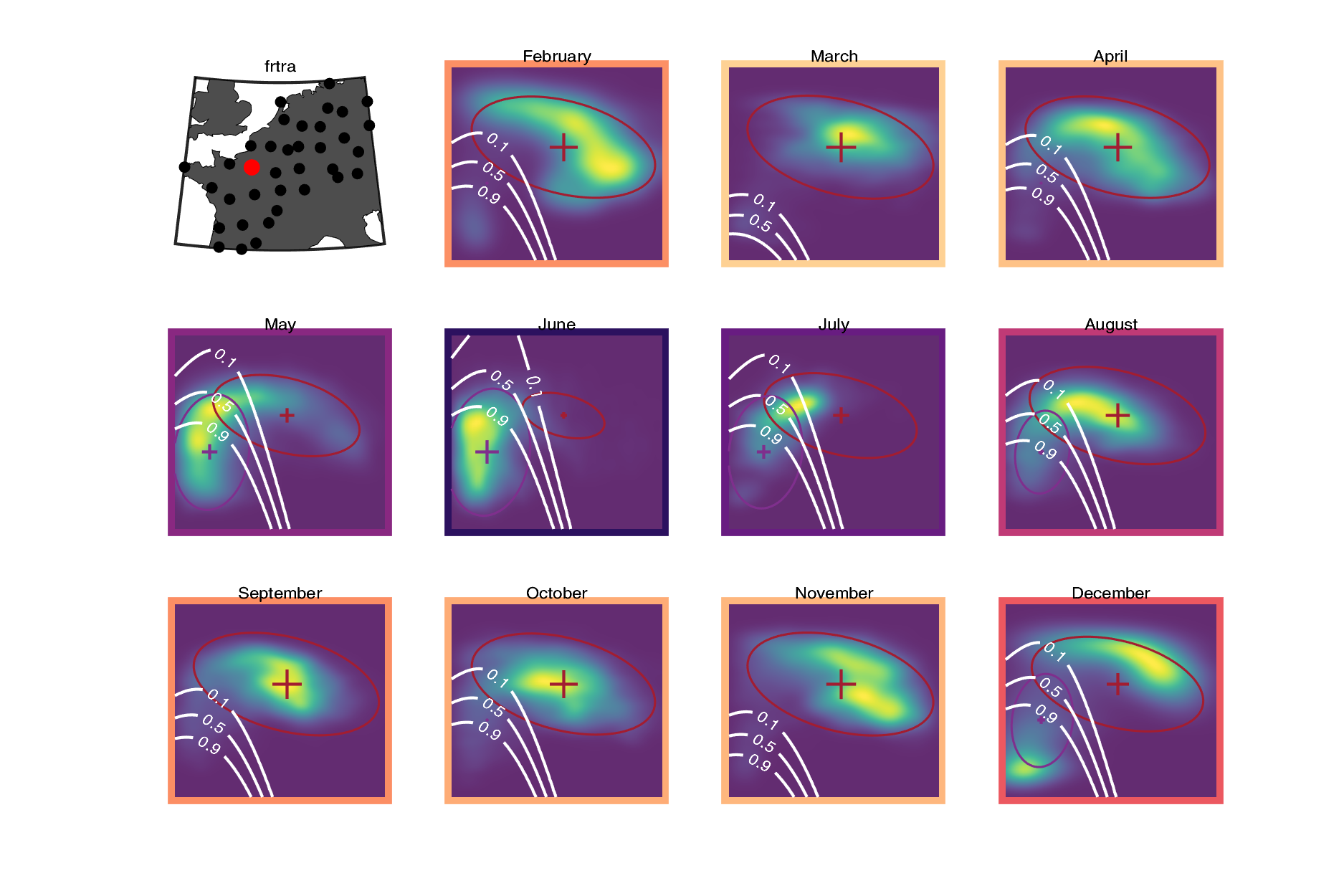

### frtre.png

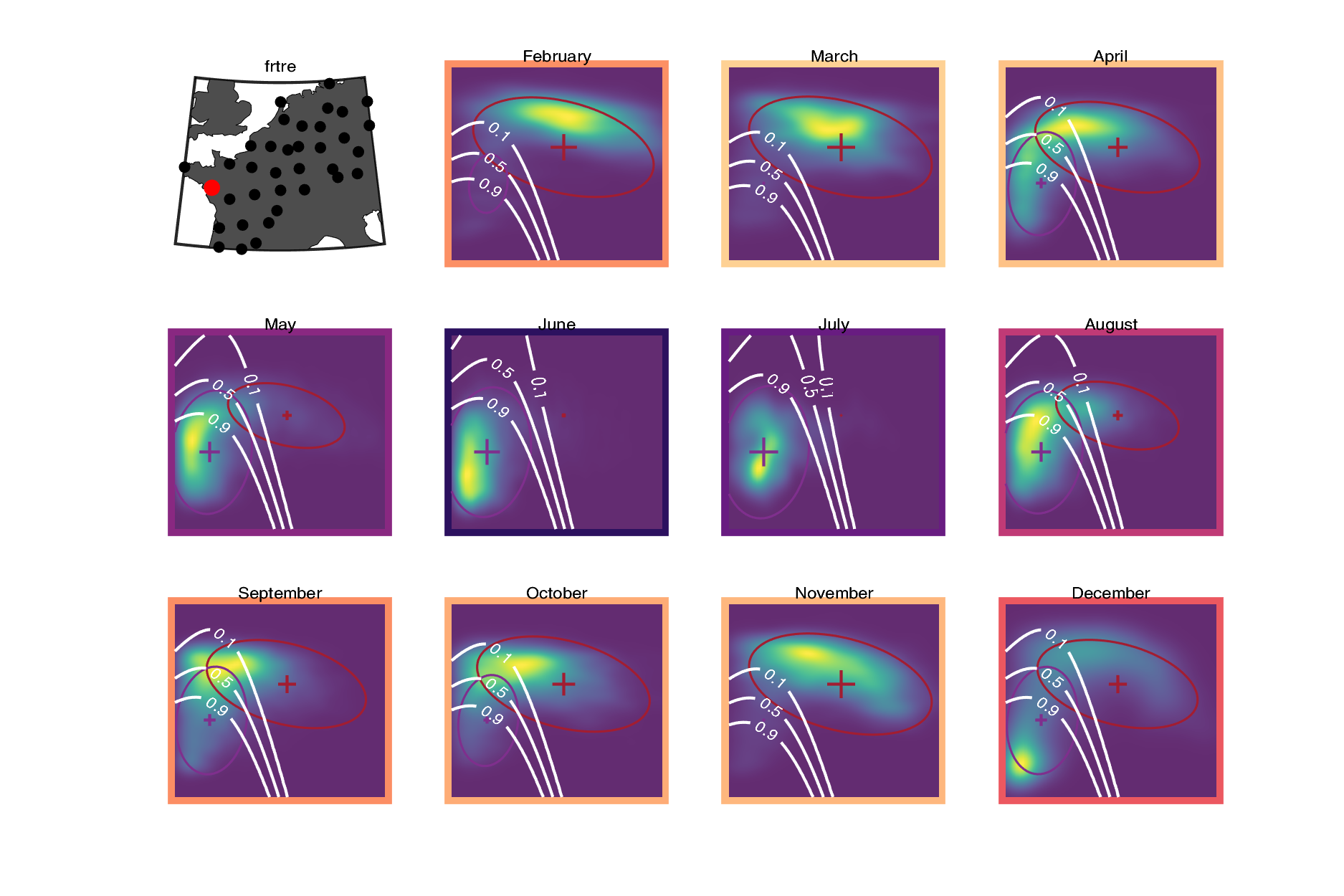

### frtro.png

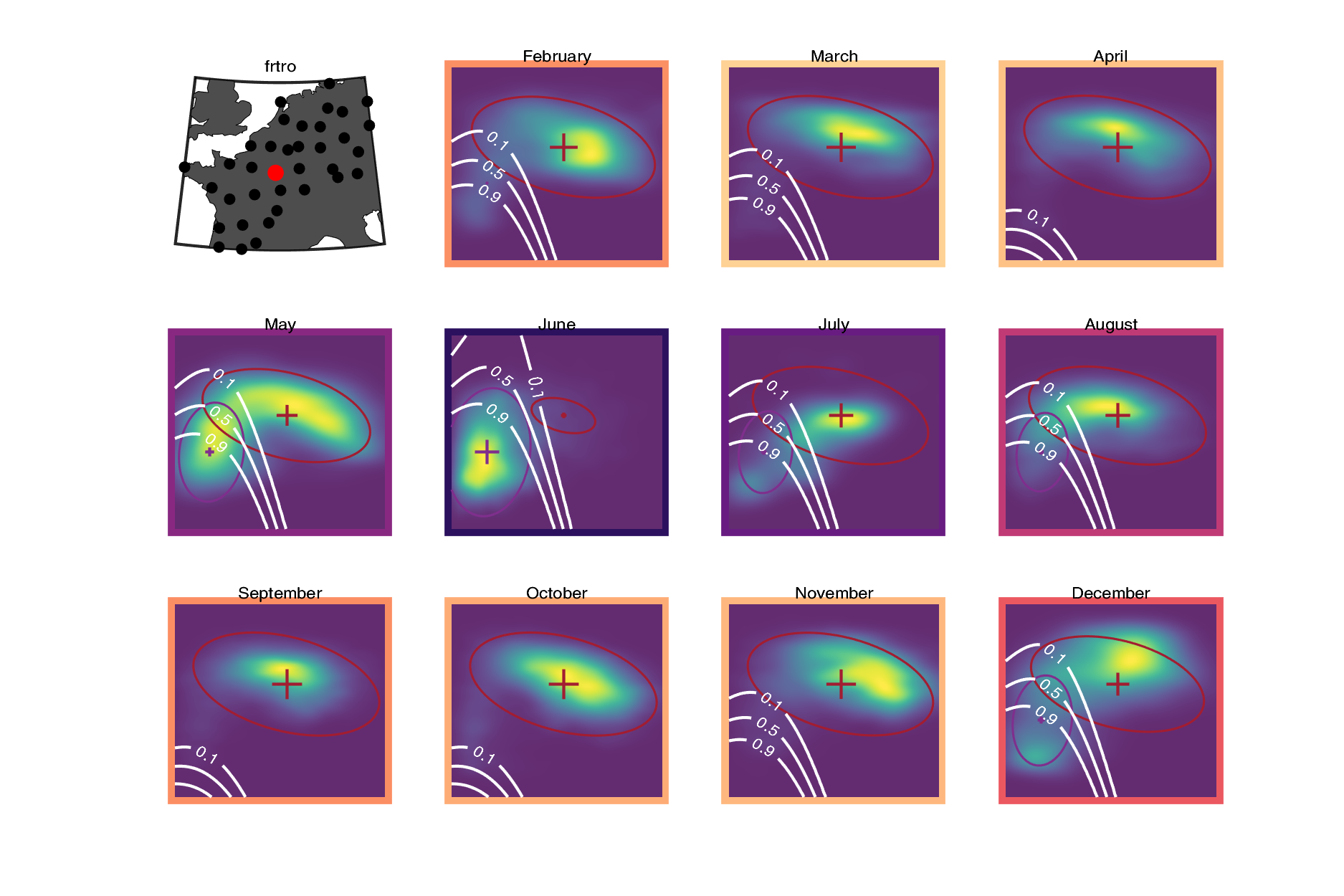

### nldhl.png

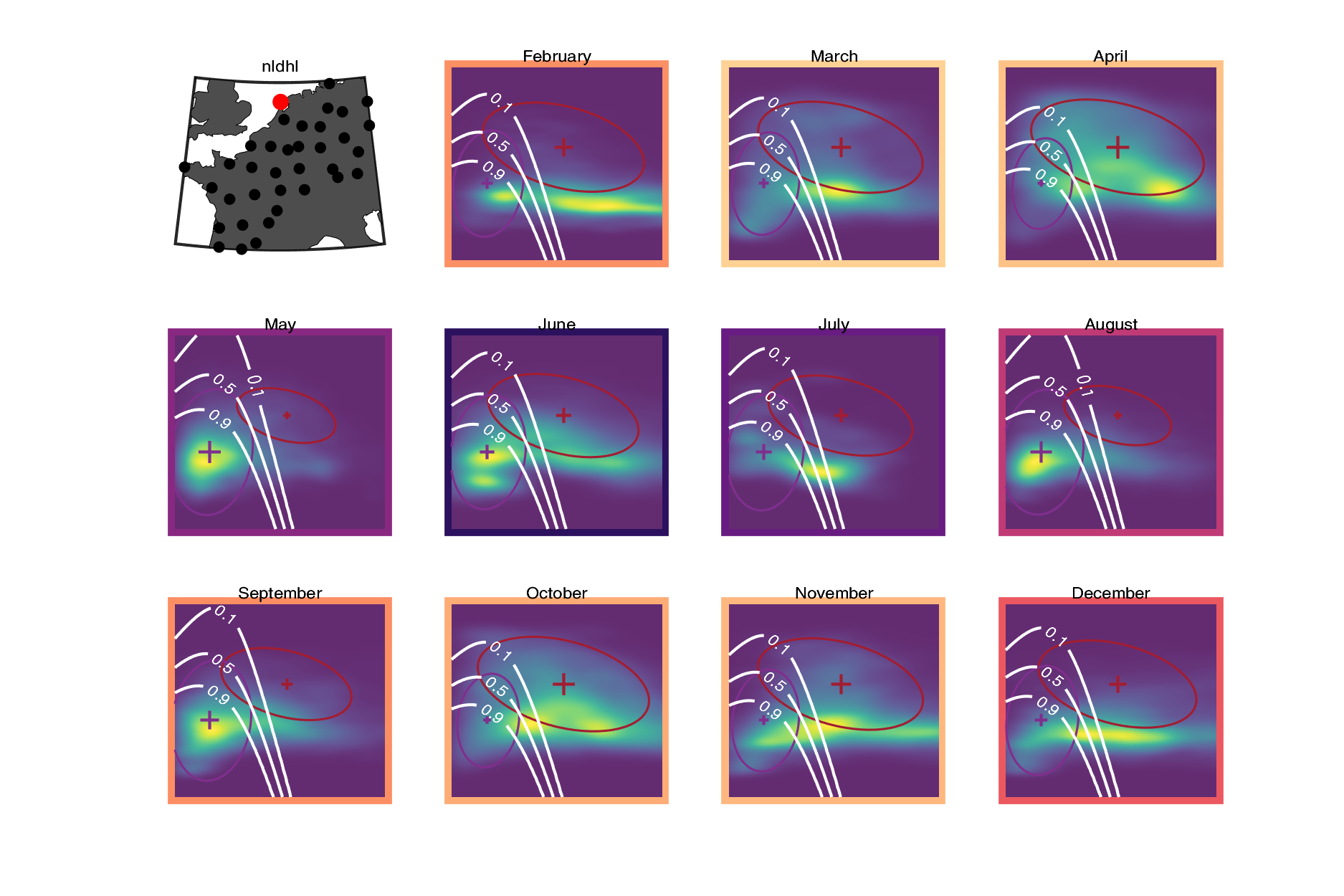

### nlhrw.png

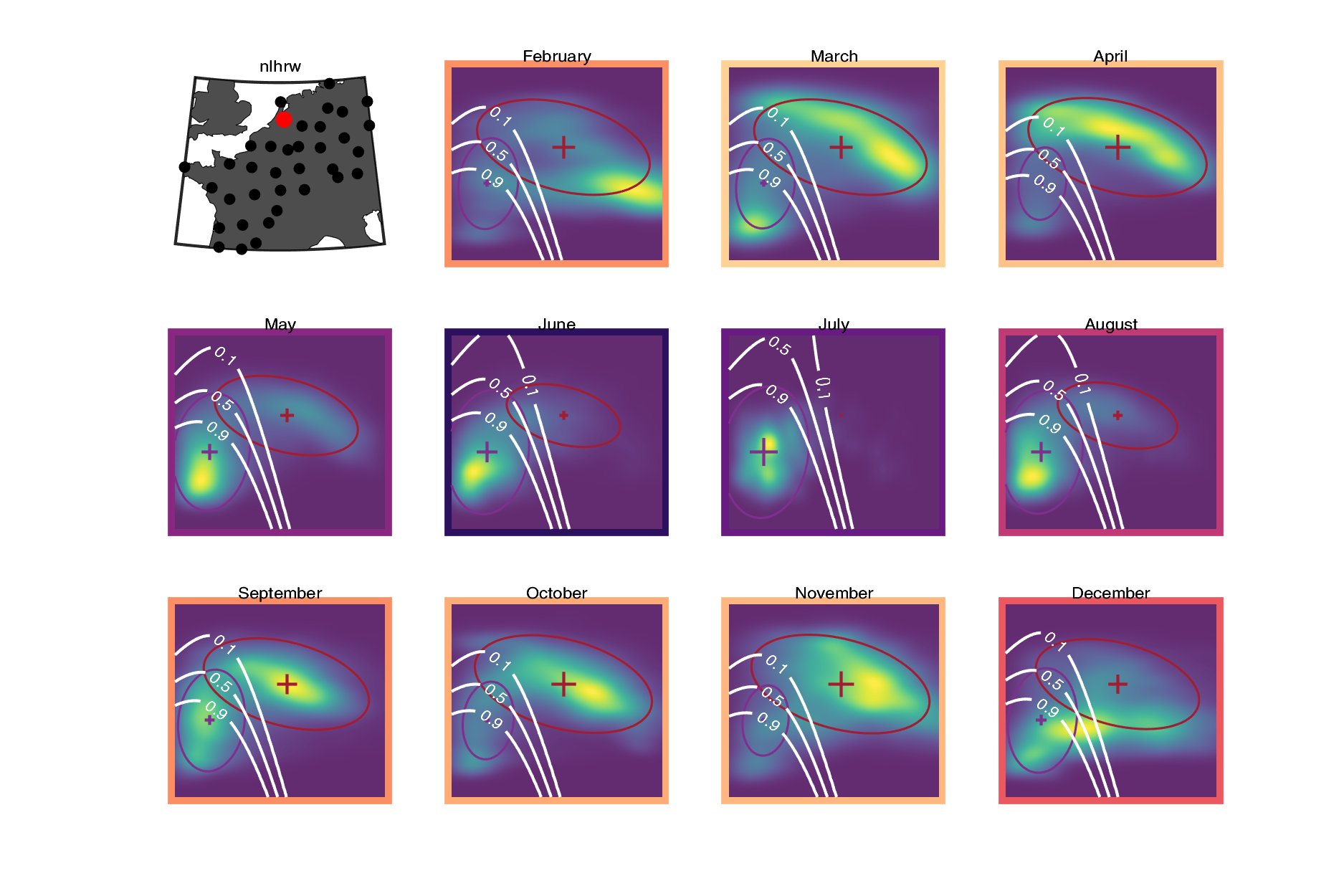

### Weekly Reflectivity Traffic Rate

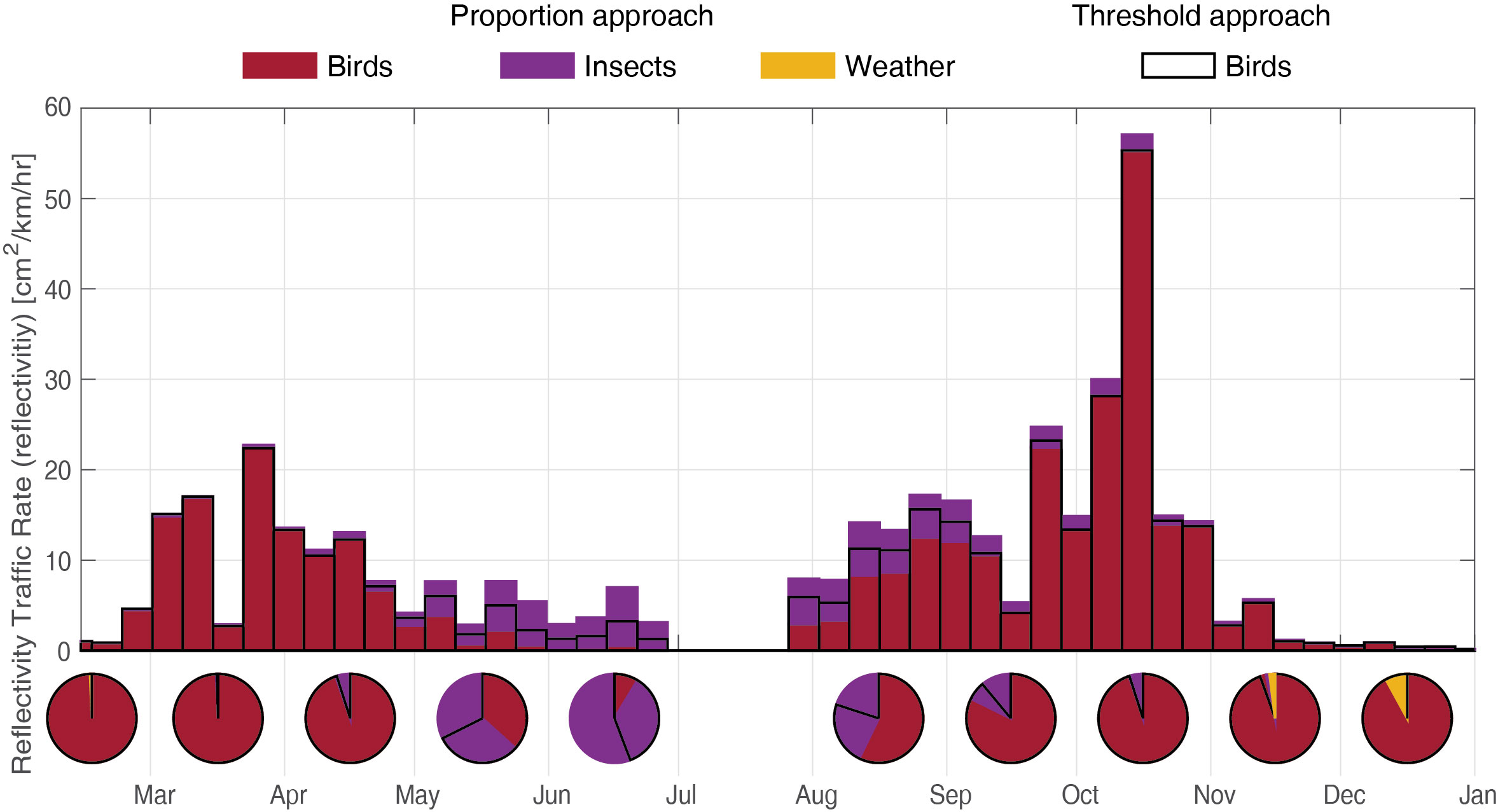
