## Supplementary material for "A simple method to separate birds and insects in single-pol weather radar data": MATLAB code

### Introduction

This code accompagne the publication [...]. It explains how the methodology is implemented, how the figure are produced and how the resulting dateset is exported (available on zenodo: https://doi.org/10.5281/zenodo.3610184)

Table of Contents

Introduction
Setup
 Load the data
 Determine windspeed at radar location
 Compute airspeed and standar deviation v\_a radial velocity sdvvp
Model fitting
 Compute empirical PDF and fit with a mixture of Gaussian
 Compute amplitude ratio
 Compute amplitude ratio over time
 Compute amplitude ratio over space
 Compute amplitude ratio over altitude
Compute amplitude ratio per radar and month
 Temporal interpolation of the amplitude ratio
 Separation of insect and weather
 Correction for bird and insect speed
Apply model to the dataset
 Export the data
Result

### Setup

#### Load the data

load('data/dc\_corr')

load('coastlines.mat')

Define default color as viridis. Code can be downloadeded from https://www.mathworks.com/matlabcentral/fileexchange/51986-perceptually-uniform-colormaps

if (exist('viridis'))

set(0,'DefaultFigureColormap',viridis);

else

web('https://www.mathworks.com/matlabcentral/fileexchange/51986-perceptually-uniform-colormaps')

end

#### Determine windspeed at radar location

The U (parmID=131) and V (parmID=132) component of wind are downloaed from the ERA5 reanalysis (https://doi.org/10.24381/cds.bd0915c6) at pressure level (from 1000hPa to 550hPa), downloaded at the maximal resoluation (hourly and 0.25°x0.25°). See python file ECMWF/script\_ECMWF.py for more details on the download.

%ncdisp(file{1});

file={'./ECMWF/2018\_pressure\_1.nc','./ECMWF/2018\_pressure\_2.nc','./ECMWF/2018\_pressure\_3.nc','./ECMWF/2018\_pressure\_4.nc'};

wind.time = datenum('01-janv-2018'):1/24:datenum('31-dec-2018 23:00');

wind.latitude=flip(double(ncread(file{1},'latitude')));

wind.longitude=double(ncread(file{1},'longitude'));

wind.pressure=double([ncread(file{1},'level') ; ncread(file{2},'level') ; ncread(file{3},'level') ; ncread(file{4},'level') ]);

wind.alt = (1-(wind.pressure\*100/101325).^(1/5.25588))/2.25577/10^(-5);

tmp\_1 = permute(flip(ncread(file{1},'u'),2) , [2 1 4 3]);

tmp\_2 = permute(flip(ncread(file{2},'u'),2) , [2 1 4 3]);

tmp\_3 = permute(flip(ncread(file{3},'u'),2) , [2 1 4 3]);

tmp\_4 = permute(flip(ncread(file{4},'u'),2) , [2 1 4 3]);

wind.u = cat(4,tmp\_1,tmp\_2,tmp\_3,tmp\_4); % m/s

tmp\_1 = permute(flip(ncread(file{1},'v'),2) , [2 1 4 3]);

tmp\_2 = permute(flip(ncread(file{2},'v'),2) , [2 1 4 3]);

tmp\_3 = permute(flip(ncread(file{3},'v'),2) , [2 1 4 3]);

tmp\_4 = permute(flip(ncread(file{4},'v'),2) , [2 1 4 3]);

wind.v = cat(4,tmp\_1,tmp\_2,tmp\_3,tmp\_4); % m/s

tmp\_1 = permute(flip(ncread(file{1},'t'),2) , [2 1 4 3]);

tmp\_2 = permute(flip(ncread(file{2},'t'),2) , [2 1 4 3]);

tmp\_3 = permute(flip(ncread(file{3},'t'),2) , [2 1 4 3]);

tmp\_4 = permute(flip(ncread(file{4},'t'),2) , [2 1 4 3]);

wind.t = cat(4,tmp\_1,tmp\_2,tmp\_3,tmp\_4); % K

clear tmp\_\* file

All three variables are linearly interpolated (time-space 4D) at each datapoint of the weather radar data.

Fu = griddedInterpolant({wind.latitude,wind.longitude,datenum(wind.time),wind.alt},wind.u,'linear','linear');

Fv = griddedInterpolant({wind.latitude,wind.longitude,datenum(wind.time),wind.alt},wind.v,'linear','linear');

Ft = griddedInterpolant({wind.latitude,wind.longitude,datenum(wind.time),wind.alt},wind.t,'linear','linear');

for i\_d=1:numel(dc)

windu = permute(Fu({dc(i\_d).lat,dc(i\_d).lon,datenum(dc(i\_d).time),dc(1).alt}),[3,4,1,2]);

windv = permute(Fv({dc(i\_d).lat,dc(i\_d).lon,datenum(dc(i\_d).time),dc(1).alt}),[3,4,1,2]);

dc(i\_d).wt = permute(Ft({dc(i\_d).lat,dc(i\_d).lon,datenum(dc(i\_d).time),dc(1).alt}),[3,4,1,2]);

dc(i\_d).ws = windu + 1i\*windv;

end

Export dataset and clear

clear wind Fu Fv

% save('data/dc\_corr','dc','start\_date','end\_date','quantity','-v7.3')

#### Compute airspeed and standar deviation v\_a radial velocity sdvvp

v\_a = nan(numel(dc(1).time), numel(dc(1).alt), numel(dc));

sdvvp = v\_a;

for i\_d=1:numel(dc)

% Compute the n/s and e/w componenent of the flight speed

% [-u,v] = pol2cart(deg2rad(dc(i\_d).dd+90), dc(i\_d).ff);

% u = dc(i\_d).ff .\* sind(dc(i\_d).dd); % m/s | 0° is north and 90° is east. -> u is east (+) - west (-)

% v = dc(i\_d).ff .\* cosd(dc(i\_d).dd); % m/s -> v is north (+) - south (-)

% gs = u + 1i\*v;

gs = dc(i\_d).ff .\* exp(deg2rad(mod(-dc(i\_d).dd+90,360))\*1i);

% Remove data not cleaned for density (e.g. rain).

id = isnan(dc(i\_d).dens3) | dc(i\_d).dens3==0;

gs(id) = nan;

sd\_vvp = dc(i\_d).sd\_vvp;

sd\_vvp(id) = nan;

v\_a(:,:,i\_d) = gs - dc(i\_d).ws;

sdvvp(:,:,i\_d) = sd\_vvp;

end

Plot airpseed and sdvvp histogram

figure('position',[0 0 800 400]); hold on;

histogram(abs(v\_a),'EdgeColor','none','DisplayName','Airspeed')

histogram(sdvvp,'EdgeColor','none','DisplayName','SD radial velocity')

legend; xlabel('speed [m/s]'); xlim([0 25])

### Model fitting

#### Compute empirical PDF and fit with a mixture of Gaussian

Avoid to recompute the gmfit and the ampli\* and f\*

load('data/insect\_removal')

Define variable used later

t = datenum(dc(1).time)-datenum(dc(1).time(1));

nmonth = 12;

tmonth = linspace(t(1), t(end), nmonth+1);

tweek = t(1):7:t(end);

nweek = numel(tweek);

tmonth\_mid = tmonth(1:end-1)+diff(tmonth)/2;

tmonth\_mid\_rep = [tmonth\_mid-365 tmonth\_mid tmonth\_mid+365]

France and German radar

id\_de = [false; true(15,1); false(21,1)];

id\_fr = [false(16,1); true(19,1); false(2,1)];

Build the vector of data removing nan value

X = [abs(v\_a(:)) sdvvp(:)];

X = X(~any(isnan(X),2),:);

Fit a kernel density function

[xi1, xi2] = meshgrid(0:.25:15,0:.1:7);

ft = reshape(ksdensity(X, [xi1(:), xi2(:)]),size(xi1));

ft = ft./sum(ft(:));

Fit Gaussian mixture model with two components

S.mu = [8 4 ; 2.5 3];

S.Sigma = cat(3,[11 -1 ; -1 1], [2 0.1 ; 0.1 1]);

S.ComponentProportion = [0.7 0.3];

rng('default')

gmfit = fitgmdist(X,2,'Replicates',1,'Start',S);

Compute the normalized pdf of each gaussian

gm1 = gmdistribution(gmfit.mu(1,:), gmfit.Sigma(:,:,1));

f\_1 = reshape(pdf(gm1,[xi1(:) xi2(:)]),size(xi1));

f\_1 = f\_1./sum(f\_1(:));

gm2 = gmdistribution(gmfit.mu(2,:), gmfit.Sigma(:,:,2));

f\_2 = reshape(pdf(gm2,[xi1(:) xi2(:)]),size(xi1));

f\_2 = f\_2./sum(f\_2(:));

Plot the fitted pdf

figure('position',[0 0 300 300]); hold on; hold on

% plot(xi1(islocalmax(ft)&islocalmax(ft,2)), xi2(islocalmax(ft)&islocalmax(ft,2)),'.r')

surf(xi1, xi2, ft,'FaceAlpha',0.5)

[~,p1]=contour3(xi1, xi2, gmfit.ComponentProportion(1)\*f\_1,10,'Color',[162 29 49]/255,'linewidth',1.5);

[~,id\_max] = max(f\_1(:));

plot3(xi1(id\_max),xi2(id\_max),gmfit.ComponentProportion(1)\*f\_1(id\_max),'+','Color',[162 29 49]/255,'LineWidth',2,'MarkerSize',6)

plot3([xi1(id\_max) xi1(id\_max)],[xi2(id\_max) xi2(id\_max)], [0 gmfit.ComponentProportion(1)\*f\_1(id\_max)],'Color',[162 29 49]/255,'LineWidth',1.5 )

[~,p2]=contour3(xi1, xi2, gmfit.ComponentProportion(2)\*f\_2,10,'Color',[127 47 141]/255,'linewidth',1.5);

[~,id\_max] = max(f\_2(:));

plot3(xi1(id\_max),xi2(id\_max),gmfit.ComponentProportion(2)\*f\_2(id\_max),'+','Color',[127 47 141]/255,'LineWidth',2,'MarkerSize',6)

plot3([xi1(id\_max) xi1(id\_max)],[xi2(id\_max) xi2(id\_max)], [0 gmfit.ComponentProportion(2)\*f\_2(id\_max)],'Color',[127 47 141]/255,'LineWidth',1.5 )

axis tight;shading interp; view(37,31)

xlabel('airspeed [m/s]'); ylabel('\sigma\_{vvp} [m/s]'); zlabel('PDF')

%legend([p1 p2], 'Multi-normal corresponding to bird','Multi-normal corresponding to insect')

Ax = gca; Ax.ZAxis.TickValues=[];

colormap(gca,viridis) %colormap(gca,1-(1-viridis)/1.2) %

Display parameters of the gaussian

disp(gmfit.mu)

7.9599 4.1032
2.6119 2.7799

disp(gmfit.Sigma)

(:,:,1) =
11.5652 -1.1574
-1.1574 0.9159
(:,:,2) =
1.8133 0.1578
0.1578 1.0829

Plot figure as a comparison with threashold

figure('position',[0 0 800 600]); hold on; hold on

h(1)=pcolor(xi1(1,:), xi2(:,1), ft./max(ft(:)));

h(2)=plot([0 15],[2 2],'r','linewidth',2);

plot([5 5],[0 7],'r','linewidth',2);

[~,h(3)]=contour(xi1, xi2, f\_1,6,'Color',[162 29 49]/255,'LineWidth',1.5);

[~,id\_max] = max(f\_1(:)); plot(xi1(id\_max),xi2(id\_max),'+','Color',[162 29 49]/255,'LineWidth',2,'MarkerSize',10)

[~,h(4)]=contour(xi1, xi2, f\_2,6,'Color',[127 47 141]/255,'LineWidth',1.5);

[~,id\_max] = max(f\_2(:)); plot(xi1(id\_max),xi2(id\_max),'+','Color',[127 47 141]/255,'LineWidth',2,'MarkerSize',10)

[C,h(5)]=contour(xi1, xi2, f\_2./(f\_1+f\_2),.1:.2:.9,'w','ShowText','on','LineWidth',2);

clabel(C,h(5),'FontSize',15,'Color','w','LabelSpacing',500,'FontWeight','bold')

axis tight;shading interp;

xlabel('airspeed [m/s]'); ylabel('\sigma\_{vvp} [m/s]');

legend(h,{'Kernel density estimation','Traditional Threashold approach','Bird Gaussian Fit','Insect Gaussian Fit','New Proportion approach'})

colormap(gca,1-(1-viridis)/1.2) %colormap(gca,viridis)

#### Compute amplitude ratio

##### Compute amplitude ratio over time

amplit = nan(1,nmonth);

ftime = nan(size(xi1,1), size(xi1,2),nmonth);

for i=1:nmonth

id = tmonth(i)<t & tmonth(i+1)>t;

X = [reshape(abs(v\_a(id,:,:)),[],1),reshape(sdvvp(id,:,:),[],1)];

X = X(~any(isnan(X),2),:);

if ~isempty(X)

ftime(:,:,i) = reshape(ksdensity(X, [xi1(:), xi2(:)]),size(xi1));

ftime(:,:,i) = ftime(:,:,i)./sum(sum(ftime(:,:,i)));

mismatch = @(alpha) ( alpha.\*f\_1 + (1-alpha).\*f\_2);

amplit(i) = fminsearch(@(alpha) sum(sum((ftime(:,:,i)-mismatch(alpha)).^2)) , 0.5 );

end

end

amplit(amplit==0)=nan;

figure('position',[0 0 1200 800]); hold on;

c\_map = magma;

for i=1:nmonth

ax=subplot(3,4,i); hold on

if i==1

colorbar

colormap(gca,magma)

else

id = tmonth(i)<t & tmonth(i+1)>t;

pcolor(xi1(1,:), xi2(:,1), ftime(:,:,i)./max(max(ftime(:,:,i))))

contour(xi1, xi2, amplit(i)\*f\_1,[0.0002 0.01],'Color',[162 29 49]/255,'LineWidth',1.5);

[~,id\_max] = max(f\_1(:));

plot(xi1(id\_max),xi2(id\_max),'+','Color',[162 29 49]/255,'LineWidth',2,'MarkerSize',amplit(i)\*20)

contour(xi1, xi2, (1-amplit(i))\*f\_2,[0.0002 0.01],'Color',[127 47 141]/255,'LineWidth',1.5);

[~,id\_max] = max(f\_2(:));

plot(xi1(id\_max),xi2(id\_max),'+','Color',[127 47 141]/255,'LineWidth',2,'MarkerSize',(1-amplit(i))\*20)

[C,h]=contour(xi1, xi2, (1-amplit(i))\*f\_2./(amplit(i)\*f\_1+(1-amplit(i))\*f\_2),[.1 .5 .9],'w','ShowText','on','LineWidth',2);

clabel(C,h,'Color','w','LabelSpacing',500,'FontWeight','bold')

axis tight;shading interp;

title(month(median(dc(1).time(id)),'name'))

%xlabel('airspeed [m/s]'); ylabel('\sigma\_{vvp} [m/s]');

colormap(gca,1-(1-viridis)/1.2)

box on

i\_c = ceil(amplit(i)\*size(c\_map,1));

ax.XColor = c\_map(i\_c,:);

ax.YColor = c\_map(i\_c,:);

ax.LineWidth=10;

ax.XTick=[]; ax.YTick=[];

end

end

##### Compute amplitude ratio over space

amplir = nan(1,numel(dc));

fradar = nan(size(xi1,1), size(xi1,2),numel(dc));

for i\_d=1:numel(dc)-1

X = [reshape(abs(v\_a(:,:,i\_d)),[],1),reshape(sdvvp(:,:,i\_d),[],1)];

X = X(~any(isnan(X),2),:);

if ~isempty(X)

fradar(:,:,i\_d) = reshape(ksdensity(X, [xi1(:), xi2(:)]),size(xi1));

fradar(:,:,i\_d) = fradar(:,:,i\_d)./sum(sum(fradar(:,:,i\_d)));

mismatch = @(alpha) ( alpha.\*f\_1 + (1-alpha).\*f\_2);

amplir(i\_d) = fminsearch(@(alpha) sum(sum((fradar(:,:,i\_d)-mismatch(alpha)).^2)) , 0.5 );

end

end

amplir(amplir==0)=nan;

figure('position',[0 0 1000 1000]); hold on;

for i\_d=1:numel(dc)-1

ax=subplot(6,6,i\_d); hold on

pcolor(xi1(1,:), xi2(:,1), fradar(:,:,i\_d)./max(max(fradar(:,:,i\_d))))

contour(xi1, xi2, amplir(i\_d)\*f\_1,5,'Color',[162 29 49]/255);

contour(xi1, xi2, (1-amplir(i\_d))\*f\_2,5,'Color',[127 47 141]/255);

[C,h]=contour(xi1, xi2, (1-amplir(i\_d))\*f\_2./(amplir(i\_d)\*f\_1+(1-amplir(i\_d))\*f\_2),[.1 .5 .9],'w','ShowText','on','LineWidth',2);

clabel(C,h,'Color','w','LabelSpacing',500,'FontWeight','bold')

axis tight;shading interp; title(dc(i\_d).name)

ax.XTick=[]; ax.YTick=[];

%xlabel('airspeed [m/s]'); ylabel('\sigma\_{vvp} [m/s]');

end

##### Compute amplitude ratio over altitude

amplia = nan(1,25);

falt = nan(size(xi1,1), size(xi1,2), 25);

for i=1:25

X = [reshape(abs(v\_a(:,i,:)),[],1),reshape(sdvvp(:,i,:),[],1)];

X = X(~any(isnan(X),2),:);

if ~isempty(X)

falt(:,:,i) = reshape(ksdensity(X, [xi1(:), xi2(:)]),size(xi1));

falt(:,:,i) = falt(:,:,i)./sum(sum(falt(:,:,i)));

mismatch = @(alpha) ( alpha.\*f\_1 + (1-alpha).\*f\_2);

amplia(i) = fminsearch(@(alpha) sum(sum((falt(:,:,i)-mismatch(alpha)).^2)) , 0.5 );

end

end

amplia(amplia==0)=nan;

figure('position',[0 0 1200 1000]); hold on; c\_map = magma;

for i=1:20

ax=subplot(4,5,i); hold on

pcolor(xi1(1,:), xi2(:,1), falt(:,:,i)./max(max(falt(:,:,i))))

contour(xi1, xi2, amplia(i)\*f\_1,5,'Color',[162 29 49]/255);

contour(xi1, xi2, (1-amplia(i))\*f\_2,5,'Color',[127 47 141]/255);

contour(xi1, xi2, (1-amplia(i))\*f\_2./(amplia(i)\*f\_1+(1-amplia(i))\*f\_2),[.1 .5 .9],'w','ShowText','on','LineWidth',2)

axis tight;shading interp; xticks([]); yticks([])

%xlabel('airspeed [m/s]'); ylabel('\sigma\_{vvp} [m/s]');

title([num2str(dc(1).alt(i)-100) ' - ' num2str(dc(1).alt(i)+100) ' m'])

box on

i\_c = ceil(amplia(i)\*size(c\_map,1));

ax.XColor = c\_map(i\_c,:);

ax.YColor = c\_map(i\_c,:);

ax.LineWidth=8;

ax.XTick=[]; ax.YTick=[];

end

### Compute amplitude ratio per radar and month

ampliall = nan(nmonth,numel(dc));

fall = nan(size(xi1,1), size(xi1,2), nweek, numel(dc));

nall = nan(nmonth,numel(dc));

for i=1:nmonth

id = tmonth(max(1,i-1))<t & tmonth(min(nweek,i+2))>t;

for i\_d=1:numel(dc)

X = [reshape(abs(v\_a(id,:,i\_d)),[],1),reshape(sdvvp(id,:,i\_d),[],1)];

X = X(~any(isnan(X),2),:);

nall(i,i\_d) = size(X,1);

if ~isempty(X)

fall(:,:,i,i\_d) = reshape(ksdensity(X, [xi1(:), xi2(:)]),size(xi1));

fall(:,:,i,i\_d) = fall(:,:,i,i\_d)./sum(sum(fall(:,:,i,i\_d)));

mismatch = @(alpha) ( alpha.\*f\_1 + (1-alpha).\*f\_2);

ampliall(i,i\_d) = fminbnd(@(alpha) sum(sum((fall(:,:,i,i\_d)-mismatch(alpha)).^2)) , 0, 1 );

end

end

end

ampliall(ampliall==0)=nan;

i\_d = find(strcmp({dc.name}, 'dedrs'));

% for i\_d=1:numel(dc)

figure('position',[0 0 1200 800]); hold on;

c\_map = magma;

for i=1:nmonth

if i==1

ax=subplot(3,4,i); hold on

title(dc(i\_d).name)

h = worldmap([min([dc.lat]) max([dc.lat])], [min([dc.lon]) max([dc.lon])]);

setm(h,'frame','on','grid','off'); set(findall(h,'Tag','MLabel'),'visible','off'); set(findall(h,'Tag','PLabel'),'visible','off')

geoshow('landareas.shp', 'FaceColor', [77 77 77]./255);

scatterm([dc.lat], [dc.lon], 60,'ok', 'filled');

scatterm(dc(i\_d).lat, dc(i\_d).lon, 120,'or','filled');

elseif all(all(isnan(fall(:,:,i,i\_d))))

%title(month(median(dc(1).time(id)),'name'))

else

ax=subplot(3,4,i); hold on

id = tmonth(i)<t & tmonth(i+1)>t;

pcolor(xi1(1,:), xi2(:,1), fall(:,:,i,i\_d)./max(max(fall(:,:,i,i\_d))))

contour(xi1, xi2, ampliall(i,i\_d)\*f\_1,[0.0002 0.01],'Color',[162 29 49]/255,'LineWidth',1.5);

[~,id\_max] = max(f\_1(:));

plot(xi1(id\_max),xi2(id\_max),'+','Color',[162 29 49]/255,'LineWidth',2,'MarkerSize',ampliall(i,i\_d)\*20)

contour(xi1, xi2, (1-ampliall(i,i\_d))\*f\_2,[0.0002 0.01],'Color',[127 47 141]/255,'LineWidth',1.5);

[~,id\_max] = max(f\_2(:));

plot(xi1(id\_max),xi2(id\_max),'+','Color',[127 47 141]/255,'LineWidth',2,'MarkerSize',(1-ampliall(i,i\_d))\*20)

[C,h]=contour(xi1, xi2, (1-ampliall(i,i\_d))\*f\_2./(ampliall(i,i\_d)\*f\_1+(1-ampliall(i,i\_d))\*f\_2),[.1 .5 .9],'w','ShowText','on','LineWidth',2);

clabel(C,h,'Color','w','LabelSpacing',500,'FontWeight','bold')

axis tight;shading interp;

title(month(median(dc(1).time(id)),'name'))

%xlabel('airspeed [m/s]'); ylabel('\sigma\_{vvp} [m/s]');

%xticks([]); yticks([]);

colormap(gca,1-(1-viridis)/1.2)

box on

i\_c = ceil(ampliall(i,i\_d)\*size(c\_map,1));

ax.XColor = c\_map(i\_c,:);

ax.YColor = c\_map(i\_c,:);

ax.LineWidth=10;

ax.XTick=[]; ax.YTick=[];

end

end

% saveas(gcf,['C:\Users\rnussba1\switchdrive\Paper\InsectWR\figure\PerRadar\' dc(i\_d).name '.png']) % ,'-dpdf -painters epsFig'

% end

Map the amplitude ratio per month and radar

figure('position',[0 0 1200 800]);

for i=1:nmonth

id = tmonth(i)<t & tmonth(i+1)>t;

subplot(3,4,i);title(month(median(dc(1).time(id)),'name'))

h = worldmap([min([dc.lat]) max([dc.lat])], [min([dc.lon]) max([dc.lon])]);

setm(h,'frame','on','grid','off'); set(findall(h,'Tag','MLabel'),'visible','off'); set(findall(h,'Tag','PLabel'),'visible','off')

geoshow('landareas.shp', 'FaceColor', [77 77 77]./255);

scatterm([dc(id\_de).lat], [dc(id\_de).lon], 180,ampliall(i,id\_de),'s', 'filled');

scatterm([dc(id\_fr).lat], [dc(id\_fr).lon], 180, ampliall(i,id\_fr),'o','filled');

scatterm([dc(~id\_fr&~id\_de).lat]', [dc(~id\_fr&~id\_de).lon]', 180, ampliall(i,~id\_fr&~id\_de)','d','filled');

caxis([0 1])

colormap(gca,magma)

if i==1

colorbar

end

end

##### Temporal interpolation of the amplitude ratio

warning('off')

ampliall(ampliall==0)=nan;

ampliall\_mean =repmat(nanmean(ampliall,2),1,37);

ampliall\_fil = ampliall;

ampliall\_fil(isnan(ampliall\_fil))=ampliall\_mean(isnan(ampliall\_fil));

ampliall\_rep = [ampliall\_fil; ampliall\_fil; ampliall\_fil];

for i\_d=1:numel(dc)

% i\_c = ceil((dc(i\_d).lat-43.5)/(54.2-43.5)\*size(c\_map,1));

ti(:,i\_d) = interp1( tmonth\_mid\_rep, ampliall\_rep(:,i\_d), t, 'pchip');

end

Figure

figure('position',[0 0 1000 450]); hold on;

plot( t, ti,'Color',[.7 .7 .7],'LineWidth',2);

scatter(repmat(tmonth\_mid',sum(id\_de),1), reshape(ampliall(:,id\_de),1,[])',[],reshape(ampliall(:,id\_de),1,[])', 's','filled','MarkerEdgeColor','k');

scatter(repmat(tmonth\_mid',sum(id\_fr),1), reshape(ampliall(:,id\_fr),1,[])',[],reshape(ampliall(:,id\_fr),1,[])', 'o','filled','MarkerEdgeColor','k');

scatter(repmat(tmonth\_mid',sum(~id\_fr&~id\_de),1), reshape(ampliall(:,~id\_fr&~id\_de),1,[])',[],reshape(ampliall(:,~id\_fr&~id\_de),1,[])', 'd','filled','MarkerEdgeColor','k');

ylim([0 1]); xlim([31 365]); datetick('x','keeplimits');

c=colorbar;c.Label.String='Ratio of amplitude';

box on; grid on

colormap(gca,magma)

#### Separation of insect and weather

gaussian = @(mu,sigma,x) exp(-((x-mu)/sigma).^2/2);

skewedgaussian = @(mu,sigma,alpha,x) 2\*gaussian(mu,sigma,x).\*normcdf(alpha\*(x-mu)/sigma);

funall= @(A1,m1,s1,a1,A2,m2,s2,a2,x) (A1\*skewedgaussian(m1,s1,a1,x) + A2\*(skewedgaussian(m2,s2,a2,x)+skewedgaussian(m2+365,s2,a2,x)));

funratio = @(A1,m1,s1,a1,A2,m2,s2,a2,x) A1\*skewedgaussian(m1,s1,a1,x)./(A1\*skewedgaussian(m1,s1,a1,x) + A2\*(skewedgaussian(m2,s2,a2,x)+skewedgaussian(m2+365,s2,a2,x)));

parmbird0=[0.9,365/2,365/15,0];

parminsect0=[0.1,0,365/15,0];

parmall0=[parmbird0 parminsect0];

for i\_d=1:numel(dc)

[xData, yData] = prepareCurveData( tmonth\_mid, 1-ampliall\_fil(:,i\_d) );

fitresult{i\_d} = fit( xData, yData, funall,...

'StartPoint', parmall0, ...

'Lower', [.5, 50, 5, -5, .01, -15, 5, -5], ...

'Upper', [ 1, 150,100, 5, 1, 50, 50, 5]);

% A1, m1, s1, a1, A2, m2, s2, a2

end

Figure

figure('position',[0 0 800 450]);

subplot(2,1,1); hold on;

for i\_d=1:numel(dc)

i\_c = ceil((dc(i\_d).lat-43.5)/(54.2-43.5)\*size(parula,1));

plot( 1:365, fitresult{i\_d}.A2\*skewedgaussian(fitresult{i\_d}.m2,fitresult{i\_d}.s2,fitresult{i\_d}.a2,1:365),'Color',[.7 .7 .7]);

plot( 1:365, fitresult{i\_d}.A1\*skewedgaussian(fitresult{i\_d}.m1,fitresult{i\_d}.s1,fitresult{i\_d}.a1,1:365),'Color',[.5 .5 .5]);

plot( 1:365, fitresult{i\_d}.A2\*skewedgaussian(fitresult{i\_d}.m2+365,fitresult{i\_d}.s2,fitresult{i\_d}.a2,1:365),'Color',[.7 .7 .7]);

end

scatter(repmat(tmonth\_mid',sum(id\_de),1), 1-reshape(ampliall(:,id\_de),1,[])',[],reshape(ampliall(:,id\_de),1,[])', 's','filled','MarkerEdgeColor','k');

scatter(repmat(tmonth\_mid',sum(id\_fr),1), 1-reshape(ampliall(:,id\_fr),1,[])',[],reshape(ampliall(:,id\_fr),1,[])', 'o','filled','MarkerEdgeColor','k');

scatter(repmat(tmonth\_mid',sum(~id\_fr&~id\_de),1), 1-reshape(ampliall(:,~id\_fr&~id\_de),1,[])',[],reshape(ampliall(:,~id\_fr&~id\_de),1,[])', 'd','filled','MarkerEdgeColor','k')

datetick('x','keeplimits');

ylim([0 1]); xlim([31 365]); datetick('x','keeplimits');

box on; grid on; legend('Insect','Weather')

ylabel('1 - Amplitude Ratio'); colormap(gca,magma)

subplot(2,1,2); hold on;

for i\_d=1:numel(dc)

i\_c = ceil((dc(i\_d).lat-43.5)/(54.2-43.5)\*size(parula,1));

plot( 1:365, funratio(fitresult{i\_d}.A1,fitresult{i\_d}.m1,fitresult{i\_d}.s1,fitresult{i\_d}.a1,fitresult{i\_d}.A2,fitresult{i\_d}.m2,fitresult{i\_d}.s2,fitresult{i\_d}.a2,1:365),'Color',[.4 .4 .4],'LineWidth',2);

end

datetick('x','keeplimits');ylim([0 1]); xlim([31 365]); grid on; box on

ylabel('Insect Ratio');

#### Correction for bird and insect speed

We can used the value fitted on the gaussians which informs us on the mean and std airspeed of insect and bird

Define airspeed range

as=-5:0.05:25;

Compute the corresponding pdf for instect and bird

pdfB = normpdf(as,gmfit.mu(1,1), sqrt(gmfit.Sigma(1,1,1)));

pdfI = normpdf(as,gmfit.mu(2,1), sqrt(gmfit.Sigma(1,1,2)));

Plot the pdfs

figure('position',[0 0 600 400]); hold on;

ksdensity(abs(v\_a(:)))

plot(as,pdfI,'DisplayName','Fitted Insect');

plot(as,pdfB,'DisplayName','Fitted Bird');

xlabel('airspeed [m/s]'); ylabel('pdf')

box on; xlim([-5 20])

l=legend(); l.String{1} ='All data empirical';

Test for synthetic case

alpha0 = 0.6;

XI = normrnd(gmfit.mu(2,1), sqrt(gmfit.Sigma(1,1,2)),100000,1);

XB = normrnd(gmfit.mu(1,1), sqrt(gmfit.Sigma(1,1,1)),100000,1);

%XI = normrnd(gmfit.mu(1,1), 2.5,1000,1);

%XB = normrnd(gmfit.mu(2,1), 2.5,1000,1);

X = alpha0\*XI + (1-alpha0)\*XB;

% fg = @(mu,sigma,x) exp(-((x-mu)/sigma).^2/2);

% fb = @(b,x,alpha) fg(gmfit.mu(1,1),sqrt(gmfit.Sigma(1,1,1)),b) .\* fg((1-alpha)\*b+alpha\*gmfit.mu(2,1), sqrt(gmfit.Sigma(1,1,2)),x);

% cb = @(x,alpha,r) integral( @(b) fb(b,x,alpha),-Inf,r) ./ integral( @(b) fb(b,x,alpha),-Inf,Inf);

% fsample = @(x,alpha) fsolve( @(r) abs(cb(x,alpha,r)-rand()), 10);

% fsample(X,alpha0)

% m1 = gmfit.mu(1,1);

% s1 = gmfit.Sigma(1,1,1);

% m2 = @(alpha,x) (x-alpha\*gmfit.mu(2,1))/(1-alpha);

% s2 = @(alpha) gmfit.Sigma(1,1,2)/(1-alpha)^2;

% m12 = @(alpha,x) (m1\*s2(alpha)+m2(alpha,x)\*s1)/(s1\*s2(alpha));

% s12 = @(alpha) (s1\*s2(alpha))/(s1+s2(alpha));

% pdf12 = @(alpha,x) normpdf(as,m12(alpha,x),sqrt(s12(alpha)));

% rnd12 = @(alpha,x) normrnd(m12(alpha,x),sqrt(s12(alpha)),1);

pBsX = pdfB .\* normpdf(X,alpha0.\* gmfit.mu(2,1) + (1-alpha0)\*as, sqrt(gmfit.Sigma(1,1,2)));

pBsX = pBsX./sum(pBsX,2);

[~,id] = min(abs(cumsum(pBsX,2) - rand(numel(X),1)),[],2);

% figure; hold on;

% tmp = cumsum(pBsX'./sum(pBsX'));

% plot(as,tmp)

% tmp = fb(as,X,alpha0);

% plot(as,cumsum(tmp)./sum(tmp))

% for i=1:numel(as)

% plot(as(i),cb(X,alpha0,as(i)),'.k')

% end

%plot(normpdf(as,m2(alpha0,X),sqrt(s2(alpha0))))

%plot(normpdf(as,m2(alpha0,X),sqrt(s2(alpha0))))

%plot(normpdf(X,alpha0.\* gmfit.mu(2,1) + (1-alpha0)\*as, sqrt(gmfit.Sigma(1,1,2))))

Xcorr = as(id)+median(diff(as))\*(rand(size(id'))-.5);

% Xcorr = X .\* (1 - alpha0 + sqrt(gmfit.Sigma(1,1,2))/sqrt(gmfit.Sigma(1,1,1)).\*alpha0) + (gmfit.mu(2,1) - gmfit.mu(1,1)\*sqrt(gmfit.Sigma(1,1,2))/sqrt(gmfit.Sigma(1,1,1)) ).\*alpha0;

%Xcorr = X\*(1-alpha0+4.1/2.8\*alpha0) + (8 - 2.6\*4.1/2.8).\*alpha0;

pIsX = pdfI .\* normpdf(X,(1-alpha0).\* gmfit.mu(1,1) + alpha0\*as, sqrt(gmfit.Sigma(1,1,1)));

pIsX = pIsX./sum(pIsX,2);

[~,id] = min(abs(cumsum(pIsX,2) - rand(numel(X),1)),[],2);

XcorrI = as(id)+median(diff(as))\*(rand(size(id'))-.5);

figure('position',[0 0 1000 400]); hold on;

plot(as,pdfB,'DisplayName','Bird','linewidth',2);

plot(as,pdfI,'DisplayName','Insect','linewidth',2);

histogram(X,'Normalization','pdf')

histogram(Xcorr,'Normalization','pdf')

histogram(XcorrI,'Normalization','pdf')

l=legend(); l.String{3}='Raw data (insect+bird)'; l.String{4}='Corrected for bird'; l.String{5}='Corrected for insect';

box on; grid on; title(['Synthetic case for \alpha=' num2str(alpha0)])

Test on v\_a (not the full dataset)

isnanv\_a=isnan(v\_a);

X=abs(v\_a(~isnanv\_a));

alpha = reshape([dc.bird],size(v\_a));

alpha0=alpha(~isnanv\_a);

subs=randi(numel(X),100000,1);

pBsX = pdfB .\* normpdf(X(subs),(1-alpha0(subs)).\* gmfit.mu(2,1) + alpha0(subs)\*as, sqrt(gmfit.Sigma(1,1,2)));

pBsX = pBsX./sum(pBsX,2);

[~,id] = min(abs(cumsum(pBsX,2) - rand(numel(X(subs)),1)),[],2);

Xcorr = as(id)+median(diff(as))\*(rand(size(id'))-.5);

figure('position',[0 0 1000 300]); hold on;

plot(as,pdfB,'DisplayName','Bird','linewidth',2);

plot(as,pdfI,'DisplayName','Insect','linewidth',2);

histogram(X(subs),100,'Normalization','pdf')

histogram(Xcorr,100,'Normalization','pdf')

legend()

### Apply model to the dataset

% r = 200; % ratio of correlation between vertical and time: 1 => 200m = 5minutes

% [X,Y]=meshgrid(linspace(1,r\*25,25),1:numel(dc(i\_d).time));

for i\_d=1:numel(dc)

% Compute insect proportion

tmp1 = repmat(ti(:,i\_d),1,25).\*reshape(pdf(gm1,[reshape(abs(v\_a(:,:,i\_d)),[],1) reshape(dc(i\_d).sd\_vvp2,[],1)]),size(v\_a(:,:,i\_d)));

tmp2 = (1-repmat(ti(:,i\_d),1,25)).\*reshape(pdf(gm2,[reshape(abs(v\_a(:,:,i\_d)),[],1) reshape(dc(i\_d).sd\_vvp2,[],1)]),size(v\_a(:,:,i\_d)));

dc(i\_d).bird = tmp1 ./(tmp1+tmp2);

% Interpolate the insect/bird ratio when dens3 exist but not sd\_vvp/u/v

idp = ~isnan(dc(i\_d).dens3) & isnan(dc(i\_d).bird);

% Fbird = scatteredInterpolant(X(~isnan(dc(i\_d).bird)), Y(~isnan(dc(i\_d).bird)), dc(i\_d).bird(~isnan(dc(i\_d).bird)),'nearest','nearest');

% dc(i\_d).bird(idp) = Fbird(X(idp), Y(idp));

Fbird = fillmissing(dc(i\_d).bird,'nearest',2);

Fbird = fillmissing(Fbird,'nearest',1);

dc(i\_d).bird(idp) = Fbird(idp);

% Separate Insect from weather

dc(i\_d).insect=(1-dc(i\_d).bird).\*funratio(fitresult{i\_d}.A1,fitresult{i\_d}.m1,fitresult{i\_d}.s1,fitresult{i\_d}.a1,fitresult{i\_d}.A2,fitresult{i\_d}.m2,fitresult{i\_d}.s2,fitresult{i\_d}.a2,datenum(dc(i\_d).time-datetime(2018,1,1)))';

% imagesc(dc(i\_d).insect','AlphaData',~isnan(dc(i\_d).insect'))

% Correct density by removing the

dc(i\_d).dens4 = dc(i\_d).dens3 .\* dc(i\_d).bird;

% Compute airspeed component (after interpolation so can't use v\_a)

airspeed\_u = dc(i\_d).u-real(dc(i\_d).ws);

airspeed\_v = dc(i\_d).v-imag(dc(i\_d).ws);

airspeed\_l2 = sqrt(airspeed\_u.^2 + airspeed\_v.^2);

% OLD

% Compute the correction factor

%cc = speed\_corr(airspeed\_l2, dc(i\_d).insect)

% apply to each component

%dc(i\_d).u2 = airspeed\_u.\*cc./airspeed\_l2 + dc(i\_d).windu;

%dc(i\_d).v2 = airspeed\_v.\*cc./airspeed\_l2 + dc(i\_d).windv;

% find nan to reduce time

idnan = ~(isnan(airspeed\_l2(:)) | isnan(dc(i\_d).insect(:)));

% Build the pdf of the posteriori distribution of airspeed

pBsX = pdfB .\* normpdf(airspeed\_l2(idnan),(1-dc(i\_d).bird(idnan)).\* gmfit.mu(2,1) + dc(i\_d).bird(idnan)\*as, sqrt(gmfit.Sigma(1,1,2)));

pBsX = pBsX./sum(pBsX,2);

% Sample randomly in the cdf

[~,id] = min(abs(cumsum(pBsX,2) - rand(sum(idnan),1)),[],2);

% reshape the value

airspeed\_l2\_corr=nan(size(airspeed\_l2));

airspeed\_l2\_corr(idnan) = as(id)+median(diff(as))\*(rand(size(id'))-.5);

% Recompute the groundspeed components

dc(i\_d).ub = airspeed\_u.\*airspeed\_l2\_corr./airspeed\_l2 + real(dc(i\_d).ws);

dc(i\_d).vb = airspeed\_v.\*airspeed\_l2\_corr./airspeed\_l2 + imag(dc(i\_d).ws);

% Same for insect

pIsX = pdfI .\* normpdf(airspeed\_l2(idnan),dc(i\_d).bird(idnan).\* gmfit.mu(1,1) + (1-dc(i\_d).bird(idnan))\*as, sqrt(gmfit.Sigma(1,1,1)));

pIsX = pIsX./sum(pIsX,2);

[~,id] = min(abs(cumsum(pIsX,2) - rand(sum(idnan),1)),[],2);

airspeed\_l2\_corr=nan(size(airspeed\_l2));

airspeed\_l2\_corr(idnan) = as(id)+median(diff(as))\*(rand(size(id'))-.5);

% Recompute the groundspeed components

dc(i\_d).ui = airspeed\_u.\*airspeed\_l2\_corr./airspeed\_l2 + real(dc(i\_d).ws);

dc(i\_d).vi = airspeed\_v.\*airspeed\_l2\_corr./airspeed\_l2 + imag(dc(i\_d).ws);

end

figure('position',[0 0 800 450]); hold on;

histogram(sqrt( ([dc.ub] - real([dc.ws])).^2 + ([dc.vb] - imag([dc.ws])).^2),100,'EdgeColor','none','DisplayName','after correction')

histogram(sqrt( ([dc.ui] - real([dc.ws])).^2 + ([dc.vi] - imag([dc.ws])).^2),100,'EdgeColor','none','DisplayName','after correction')

histogram(sqrt( ([dc.u] - real([dc.ws])).^2 + ([dc.v] - imag([dc.ws])).^2),100,'EdgeColor','none','DisplayName','before correction')

f=ksdensity(abs(v\_a(:)),as);

plot(as,pdfB./max(pdfB)\*271200,'DisplayName','Bird','linewidth',2);

plot(as,pdfI./max(pdfI)\*335700,'DisplayName','Insect','linewidth',2);

xlabel('Air speed [m/s]')

xlim([0 25]); box on;

#### Export the data

% CleaningV(dc)

Save

% save('data/dc\_corr','dc','start\_date','end\_date','quantity','-v7.3')

save('data/insect\_removal','xi1','xi2','ft','gmfit','f\_1','f\_2','amplia','ampliall','amplir','amplit','fall','falt','fradar','ft','ftime','tmonth','ti','gm1','gm2','fitresult','funratio')

### Result

[G,day\_num]=findgroups(datenum(dateshift(dc(1).time,'start','day','nearest')));

G2=repmat(G',1,25);

birdDENSmean=nan(numel(day\_num),numel(dc));

birdDENSmean\_OLD=birdDENSmean;

insectDENSmean = birdDENSmean;

birdMTRmean = birdDENSmean;

insectMTRmean = birdDENSmean;

weatherDENSmean = birdDENSmean;

birdMTRmean\_OLD = birdDENSmean;

birdDIRmean=birdDENSmean;

insectDIRmean=birdDENSmean;

for i\_d=1:numel(dc)

% Figure of proportion

% propdenssum = splitapply(@nansum,dc(i\_d).dens3(:).\*dc(i\_d).bird(:),G2(:));

% denssum = splitapply(@nansum,dc(i\_d).dens3(:),G2(:));

% plot(day\_num, propdenssum./denssum);

% datetick('x','keeplimits')

birdDENSmean(:,i\_d) = splitapply(@nanmean,nansum(dc(i\_d).dens3 .\* dc(i\_d).bird .\* .2,2),G');

insectDENSmean(:,i\_d) = splitapply(@nanmean,nansum(dc(i\_d).dens3 .\* dc(i\_d).insect .\* .2,2),G');

weatherDENSmean(:,i\_d) = splitapply(@nanmean,nansum(dc(i\_d).dens3 .\* (1-dc(i\_d).bird-dc(i\_d).insect) .\* .2, 2),G');

% /1000\*60\*60\*5 :m/s -> km/hr + bird/km^3 -> bird/km^2

tmp = dc(i\_d).dens3 .\* dc(i\_d).bird .\* .2 .\* sqrt(dc(i\_d).ub.^2+dc(i\_d).vb.^2);

birdMTRmean(:,i\_d) = splitapply(@nanmean, nansum(tmp,2) /1000\*60\*60,G');

assert(0==nansum(tmp(isnan(dc(i\_d).vb(:))&isnan(dc(i\_d).ub(:)))))

V = splitapply(@nansum, tmp(:).\*dc(i\_d).vb(:),G2(:)) ./ splitapply(@nansum, tmp(:),G2(:));

U = splitapply(@nansum, tmp(:).\*dc(i\_d).ub(:),G2(:)) ./ splitapply(@nansum, tmp(:),G2(:));

birdDIRmean(:,i\_d) = atan2d(V,U);

tmp = dc(i\_d).dens3 .\* dc(i\_d).insect.\* .2 .\* sqrt(dc(i\_d).ui.^2+dc(i\_d).vi.^2);

insectMTRmean(:,i\_d) = splitapply(@nanmean, nansum(tmp,2) /1000\*60\*60,G');

assert(0==nansum(tmp(isnan(dc(i\_d).vi(:))&isnan(dc(i\_d).ui(:)))))

V = splitapply(@nansum, tmp(:).\*dc(i\_d).vi(:),G2(:)) ./ splitapply(@nansum, tmp(:),G2(:));

U = splitapply(@nansum, tmp(:).\*dc(i\_d).ui(:),G2(:)) ./ splitapply(@nansum, tmp(:),G2(:));

insectDIRmean(:,i\_d) = atan2d(V,U);

tmp = dc(i\_d).dens3 .\* dc(i\_d).insect.\* .2 .\* sqrt(dc(i\_d).ui.^2+dc(i\_d).vi.^2);

weatherMTRmean(:,i\_d) = splitapply(@nanmean, nansum(tmp,2) /1000\*60\*60,G');

tmp = dc(i\_d).dens3;

tmp(sqrt(dc(i\_d).u.^2 + dc(i\_d).v.^2)<5)=0;

tmp(dc(i\_d).sd\_vvp<2)=0;

birdDENSmean\_OLD(:,i\_d) = splitapply(@nanmean,nansum(tmp .\* .2,2),G');

birdMTRmean\_OLD(:,i\_d) = splitapply(@nanmean,nansum(tmp.\*.2 .\* sqrt(dc(i\_d).u.^2+dc(i\_d).v.^2),2)/1000\*60\*60 ,G');

end

figure('position',[0 0 1200 300]); hold on;

b=bar(day\_num, [nanmean(birdDENSmean,2) nanmean(insectDENSmean,2) nanmean(weatherDENSmean,2)]./11,'stacked');

b(1).FaceColor = [162 29 49]/255;b(2).FaceColor = [127 47 141]/255;

datetick('x','keeplimits'); xlim([737104 737426]); ylabel('Reflectivity [cm^2/km^3]');

box on; grid on;

% yyaxis right;

stairs(day\_num-.5, nanmean(birdDENSmean\_OLD,2)./11,'k');

figure('position',[0 0 1000 800]);

gr\_day = 7;

gr = repelem(1:1000,gr\_day); gr=gr(1:length(day\_num));

subplot(5,1,[1 3]);hold on;

b=bar(day\_num(1:gr\_day:end), splitapply(@nanmean,[nanmean(birdDENSmean,2) nanmean(insectDENSmean,2) nanmean(weatherDENSmean,2)],gr')./11,1,'stacked');

b(1).FaceColor = [162 29 49]/255; b(1).EdgeAlpha=0;

b(2).FaceColor = [127 47 141]/255; b(2).EdgeAlpha=0;

b(3).FaceColor = [0.9290 0.6940 0.1250]; b(3).EdgeAlpha=0;

b2=bar(day\_num(1:gr\_day:end),splitapply(@nanmean,nanmean(birdDENSmean\_OLD,2),gr')./11,1,'k');

b2(1).FaceAlpha =0; b2(1).LineWidth=1.5;

% stairs(splitapply(@nanmean,day\_num,gr)-gr\_day/2, splitapply(@nanmean,nanmean(birdDENSmean\_OLD,2),gr')./11,'k');

datetick('x','keeplimits'); xlim([737104 737426]); ylabel('Reflectivity [cm^2/km^3]');

legend('Birds','Insects','Weather','Bird old'); box on; grid on; %ylim([0 1])

subplot(5,1,4);hold on;

b2=bar(day\_num(1:gr\_day:end),(splitapply(@nanmean,nanmean(birdDENSmean,2),gr')-splitapply(@nanmean,nanmean(birdDENSmean\_OLD,2),gr'))./11,1,'k');

b2(1).FaceAlpha =0; b2(1).LineWidth=1.5;

box on; grid on;datetick('x','keeplimits'); xlim([737104 737426]);

subplot(5,1,5);hold on;

gr = month(day\_num);

tt = splitapply(@nanmean,[nanmean(birdDENSmean,2) nanmean(insectDENSmean,2) nanmean(weatherDENSmean,2)],gr')./11;

tt2 = splitapply(@nanmean, nanmean(birdDENSmean\_OLD,2),gr')./11;

for i=[2 3 4 5 6 8 9 10 11 12]

subplot(5,12,48+i);

p=pie(tt(i,:)/nansum(tt(i,:)),{'','',''});

p(1).FaceColor = [162 29 49]/255; p(1).EdgeAlpha=0;

p(3).FaceColor = [127 47 141]/255; p(3).EdgeAlpha=0;

p(5).FaceColor = [0.9290 0.6940 0.1250]; p(5).EdgeAlpha=0;

hold on

p2 = pie(tt2(i,:)./nansum(tt(i,:)),{''});

p2(1).FaceAlpha=0;

axis equal

title(num2str(round(nansum(tt(i,:)),2)))

end

figure('position',[0 0 1000 800]);

gr\_day = 7;

gr = repelem(1:1000,gr\_day); gr=gr(1:length(day\_num));

subplot(5,1,[1 3]);hold on;

b=bar(day\_num(1:gr\_day:end), splitapply(@nanmean,[nanmean(birdMTRmean,2) nanmean(insectMTRmean,2)],gr')./11,1,'stacked');

b(1).FaceColor = [162 29 49]/255; b(1).EdgeAlpha=0;

b(2).FaceColor = [127 47 141]/255; b(2).EdgeAlpha=0;

%b(3).FaceColor = [0.9290 0.6940 0.1250]; b(3).EdgeAlpha=0;

b2=bar(day\_num(1:gr\_day:end),splitapply(@nanmean,nanmean(birdMTRmean\_OLD,2),gr')./11,1,'k');

b2(1).FaceAlpha =0; b2(1).LineWidth=1.5;

% stairs(splitapply(@nanmean,day\_num,gr)-gr\_day/2, splitapply(@nanmean,nanmean(birdDENSmean\_OLD,2),gr')./11,'k');

datetick('x','keeplimits'); xlim([737104 737426]); ylabel('Reflectivity Traffic Rate [cm^2/km^2/hr]');

legend('Birds','Insects','Bird old'); box on; grid on; %ylim([0 1])

subplot(5,1,4);hold on;

b2=bar(day\_num(1:gr\_day:end),(splitapply(@nanmean,nanmean(birdMTRmean,2),gr')-splitapply(@nanmean,nanmean(birdMTRmean\_OLD,2),gr'))./11,1,'k');

b2(1).FaceAlpha =0; b2(1).LineWidth=1.5;

box on; grid on;datetick('x','keeplimits'); xlim([737104 737426]);

subplot(5,1,5);hold on;

gr = month(day\_num);

tt = splitapply(@nanmean,[nanmean(birdMTRmean,2) nanmean(insectMTRmean,2)],gr')./11;

tt2 = splitapply(@nanmean, nanmean(birdMTRmean\_OLD,2),gr')./11;

for i=[2 3 4 5 6 8 9 10 11 12]

subplot(5,12,48+i);

p=pie(tt(i,:)/nansum(tt(i,:)),{'',''});

p(1).FaceColor = [162 29 49]/255; p(1).EdgeAlpha=0;

p(3).FaceColor = [127 47 141]/255; p(3).EdgeAlpha=0;

hold on

p2 = pie(tt2(i,:)./nansum(tt(i,:)),{''});

p2(1).FaceAlpha=0;

axis equal

title(num2str(round(nansum(tt(i,:)),2)))

end

  
